# Metal Coordinating Inhibitors of Rift Valley Fever virus Replication

**DOI:** 10.1101/2022.03.10.483712

**Authors:** Elizabeth Geerling, Valerie Murphy, E. Taylor Stone, Andreu Gazquez Casals, Mariah Hassert, Austin T. O’Dea, Feng Cao, Maureen J. Donlin, Mohamed Elagawany, Bahaa Elgendy, Vasiliki Pardali, Erofili Giannakopoulou, Grigoris Zoidis, Daniel V. Schiavone, Alex J. Berkowitz, Nana B. Agyemang, Ryan P. Murelli, John E. Tavis, Amelia K. Pinto, James D. Brien

## Abstract

Rift Valley Fever Virus (RVFV) is a veterinary and human pathogen and is an agent of bioterrorism concern. Currently, RVFV treatment is limited to supportive care, so new drugs to control RVFV infection are urgently needed. RVFV is a member of the Bunyavirales order, and replication of these viruses depends on the viral endonuclease activity of the viral L protein. Screening for RVFV replication inhibitors among compounds with divalent cation-coordinating motifs similar to known viral nuclease inhibitors identified 31 novel RVFV inhibitors with selective indexes from 5 – 402 and 50% effective concentrations of 0.54 – 56 µM in Vero cells, primarily α-Hydroxytropolones and N-Hydroxypyridinediones. Inhibitor activity and selective index was validated in the human cell line A549. To evaluate specificity, select compounds were tested against another Bunyavirus, La Crosse Virus (LACV). Conservation of the enzymatic activity such as the cap-snatching mechanism among the Bunyavirales implies that the α-Hydroxytropolone and N-Hydroxypyridinedione chemotypes hold potential for development into treatments for related pathogens, including Hantaan Virus, Severe fever with thrombocytopenia syndrome virus, Crimean-Congo Hemorrhagic Fever Virus, and LACV. Keywords: Rift Valley Fever Virus 1, La Crosse virus 2, Cap-snatching endonuclease 3, Replication inhibitors 4, α-Hydroxytropolones 5, N-Hydroxypyridinediones 6.

## 1. Introduction

Bunyavirales is a large order of enveloped viruses with a segmented, negative-polarity, single-stranded RNA genome and includes multiple members that pose a significant risk including Rift Valley Fever Virus (RVFV), La Crosse virus (LACV), Hantaan Virus, Severe Fever with Thrombocytopenia virus (SFTSV), and Crimean-Congo Hemorrhagic Fever Virus (CCHFV) [1,2]. RVFV is an arbovirus transmitted to animals by mosquito vectors, and is traditionally endemic in eastern and southern Africa, but has recently expanded its range throughout sub-Saharan Africa and parts of the Middle East. RVFV is a serious veterinary pathogen, causing Rift Valley Fever in domestic animals including cattle, horses, sheep, goats, and camels. Rift Valley Fever is characterized by fever, hemorrhage, diarrhea, death, and nearly complete spontaneous abortions in infected animals, and often has a severe economic impact on affected herds. Veterinary outbreaks of RVFV infections can reach epidemic proportions, particularly in rainy years.

Humans can be infected by RVFV via contact with infected animal body fluids or tissues, by breathing aerosols contaminated with RVFV, or less frequently via mosquito bites; human to human transmission of RVFV is rare [2]. Most infections are either asymptomatic or cause mild fever with hepatic involvement. However, 8-10% of infections can become severe, where symptoms can include lesions to the eye, causing blindness in 50% of ocular cases (1-10% of all infections), encephalitis, gastrointestinal dysfunction, jaundice, joint/muscle pain, hemorrhagic fever, disorientation/hallucination, and partial paralysis. Hemorrhagic fever is rare (∼1% of cases) but has a ∼50% fatality rate. Human RVFV infections can be diagnosed by ELISA or RT-PCR assays, but treatment is limited to supportive care [2]. There is a live-attenuated veterinary vaccine approved for RVFV (MP12), which has undergone a phase I clinical trial for use in humans.

In addition to RVFV, LACV is an arbovirus found throughout the midwestern children and is commonly underdiagnosed because of the lack of available therapeutics [3,4]. Currently there are 50-150 cases of neuroinvasive disease reported annually [5]. Since 2011, LACV has continued to spread beyond the midwest and into the northeast, mid-Atlantic and southern states, resulting in 700 cases of neuroinvasive disease since 2011. Similar to RVFV, there are no antivirals or vaccines available to treat LACV infection or prevent disease.

Transcription of bunyavirus mRNAs uses a “cap-snatching” mechanism to provide the 5’ cap for the viral mRNAs [1,6,7]. Cap-snatching involves cleavage of a short, capped RNA oligonucleotide from the 5’ end of host cell mRNAs and uses the oligonucleotide to prime viral mRNA transcription. RNA cleavage to acquire the host-derived mRNA cap is catalyzed by a viral endonuclease located in the amino-terminus of the viral L protein. The endonuclease active site has an H..PD..D/E..K motif that binds Mg++ ions that are essential for catalysis and hence viral replication [1]. This is analogous to the Mg++-binding motif of the Influenza Virus PA cap-snatching endonuclease [8] and bears significant similarity to the D..E..D..D or D..D..E motifs found in ribonucleases H and viral integrases [9].

Inhibiting viral endonucleases can block viral replication, as has been shown for the HIV integrase [10], the HIV ribonuclease H [11], the Hepatitis B Virus (HBV) ribonuclease H [12], and the influenza virus cap-snatching enzyme [13,14]. The most common mode of inhibition is for small molecules to chelate the Mg++ ions in viral nuclease active sites, with specificity and affinity modulated by additional contacts between the inhibitors and the enzymes [15-17], and sometimes also by contacts with the nucleic acid substrate [18]. This metal-chelating mechanism is used by the HIV integrase inhibitors Bictegravir, Dolutegravir, Elvitegravir, and Raltegravir, and the Influenza Virus PA cap-snatching endonuclease inhibitor Baloxavir marboxil. The US Food and Drug Administration has approved 62 drugs that act by coordinating active-site cations in metalloenzymes as of 2017 [19], making active site metal ion chelation a well-established drug mechanism.

In these studies, we hypothesized that metal chelating compounds similar to inhibitors of the HIV and HBV ribonucleases H, the HIV integrase, and the Influenza Virus PA endonuclease would inhibit RVFV and LACV replication. This hypothesis is based on i) the inhibitory mechanism employed by metal chelating compounds against metallonucleases, ii) the essential nature of the L protein endonuclease for viral replication [7], iii) the structural similarities of Mg++-dependent viral endoribonucleases [9,20], and iv) the successes in developing drugs for HIV and Influenza virus that act by chelating the catalytic Mg++ ions of endonucleases.

## 2. Materials and Methods

### 2.1. Compound acquisition and synthesis

Commercially acquired compounds are indicated by the vendor’s name and catalog number in Tables 1 and 2.

**Table 1.**
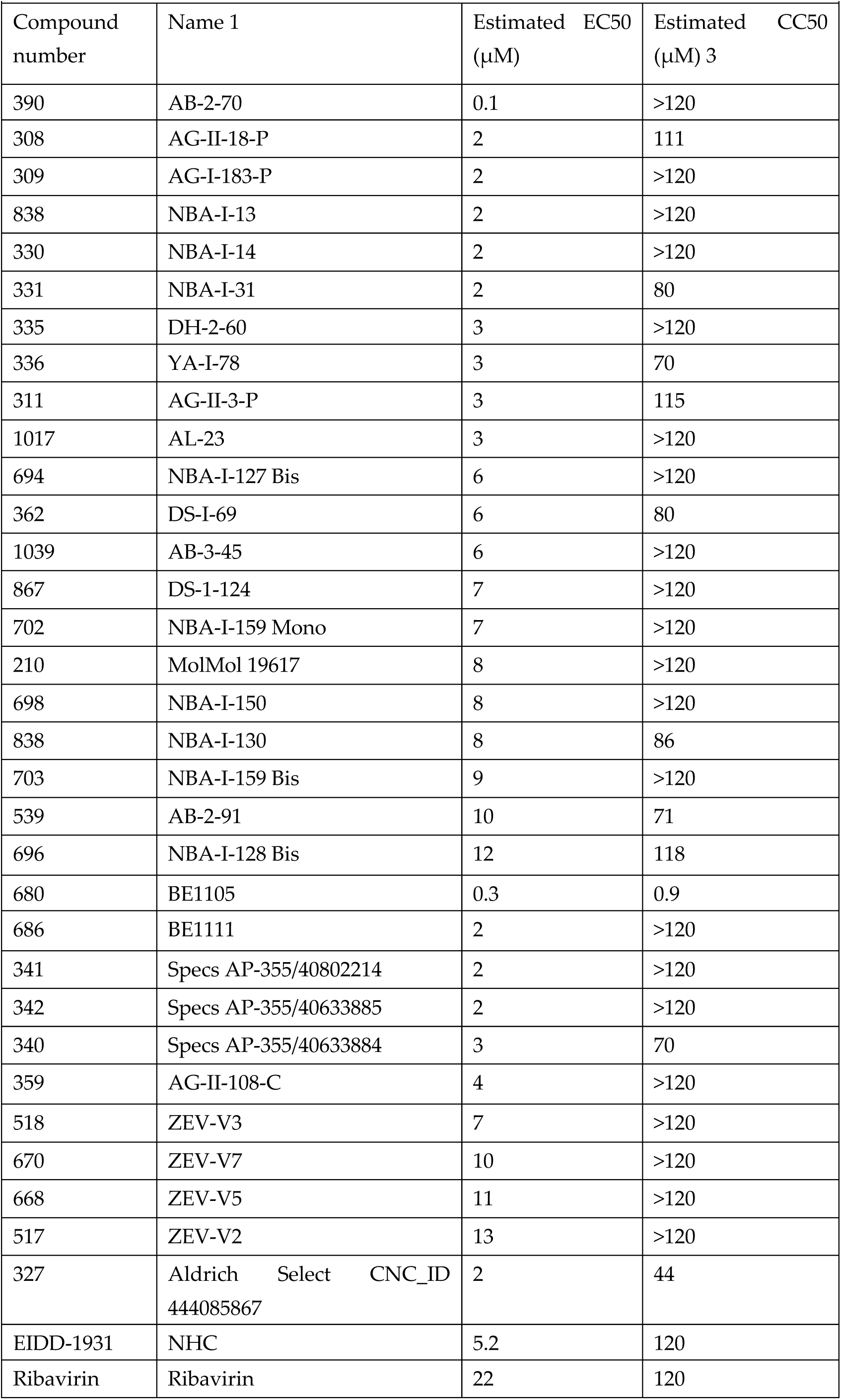
Top hits in the primary antiviral screen.

**Table 2.** EC50 and CC50 of select compounds against RVFV replication in Vero and A549 cells.

|  |  |  |  |  |  |  |  |  |
| --- | --- | --- | --- | --- | --- | --- | --- | --- |
| Compound number | Name | Chemotype | RVFV Vero (EC50, μM) | Vero (CC50 μM) | SI (Vero) | RVFV A549 (EC50, μM) | A549 (CC50 μM) | SI (A549) |
| 308 | AG-II-18-P | αHT | 1.2 | >120 | 25.8 | 8.4 | 119.0 | 14.2 |
| 309 | AG-I-183-P | αHT | 1.6 | >120 | 75.0 | 5.9 | 98.1 | 16.8 |
| 320 | NBA-I-13 | αHT | 34 | >120 | 49.5 | 90.3 | 154.1 | 1.7 |
| 327 | Aldrich Select CNC_ID 444085867 | DHN | 39.7 | 42.9 | 18.5 | 19.6 | 27.5 | 1.4 |
| 330 | NBA-I-14 | αHT | 10.4 | >120 | 56.2 | 42.6 | >240 | 5.6 |
| 331 | NBA-I-31 | αHT | 15.8 | 80.4 | 36.6 | 17.7 | >240 | 13.6 |
| 335 | DH-2-60 | αHT | 40.8 | >120 | 78.0 | >120 | 96.2 | 0.3 |
| 390 | AB-2-70 | αHT | 11.9 | >120 | 29.8 | 9.4 | 224.2 | 23.9 |
| 515 | ZEV-E2 | HPD | 24.3 | >120 | 71.3 | 150.7 | 93.6 | 0.6 |
| 517 | ZEV-V2 | HPD | >120 | >120 | 91.0 | 95.8 | 71.8 | 0.7 |
| 518 | ZEV-V3 | HPD | 20.0 | >120 | 91.4 | 17.2 | 16.5 | 1.0 |
| 668 | ZEV-V5 | HPD | 19.2 | >120 | 76.8 | 35.7 | 82.0 | 2.3 |
| 670 | ZEV-V7 | HPD | 14 | >120 | 69.0 | 15.3 | >240 | 15.7 |
| 680 | BE1105 | TTP | 6.3 | >120 | 19.0 | 11.6 | >240 | 20.7 |
| 686 | BE1111 | TTP | 6.3 | >120 | 19.0 | 11.6 | 67.6 | 8.6 |
| 704 | NBA-I-160 | αHT | 14.3 | >120 | 21.6 | 20.0 | 43.0 | 2.2 |
| 840 | NBA-I-155-Mono | αHT | 31.6 | >120 | 21.5 | 15.4 | 141.6 | 9.2 |
| 867 | DS-1-124 | αHT | 8.7 | >120 | 5.2 | 33.0 | >240 | 7.3 |
| 1039 | AB-3-45 | αHT | 8.9 | >120 | 8.7 | 15.5 | >240 | 15.5 |

TTP Compounds were synthesized as described in [21].

αHT compounds were synthesized as described in [22]. For 265, 308, and 311 see [23]. For 169 and 362 see [24]. For 385, 694, 696, 698, 700, 702, 704, 703, 838, and 840 see [25]. For 321, 336, 358, and 359 see [26]. For 388, 389, 390 and 539 see [27]. For 330 and 331 see [28]. For 113, 118 and 120 see [29]. For 260 see [30]. For 111 see [22]. For 335 see [31].

The novel HPD compounds were synthesized in a three-step procedure outlined in Figure 2. Synthetic validation is in Supplemental Data File 1. Briefly, the key structure 5-acetyl-1-(benzyloxy)-6-hydroxy-4-methylpyridin-2(1H)-one (ZEV1) was synthesized with an improved yield of 75% by refluxing a mixture of O-benzyl hydroxylamine (1 eq) and diketene (2 eq) in the presence of triethylamine (1 eq) in dry toluene. Next, the benzyl group was cleaved by catalytic hydrogenation over 10% palladium on carbon leading to compound ZEV2. 5-Acetyl-1,6-dihydroxy-4-methylpyridin-2(1H)-one (ZEV2) was coupled with the appropriate substituted aniline using sulfuric acid as catalyst in absolute ethanol at reflux. The desired compounds were obtained in good yields ranging from 60% to 70%, with the only exception being 668 which was isolated in an overall yield of 25%. Compounds were diluted to 10 mM in DMSO and stored in single-use aliquots in opaque tubes at -20oC.

### 2.2. Cells and Viruses

RVFV strains MP-12 and ZH501, as well as LACV strain original (BEI NR-540) were passaged in Vero E6 cells (ATCC® CRL-1586™) before clarification by centrifugation at 3,000 rpm for 30 minutes and stored at -80°C until further use. RVFV isolates were a kind gift of Drs. M. Buller and A. Hise. MP-12 is a BSL-2 vaccine strain of RVFV and is not classified as a select agent allowing easier assay development. RVFV ZH501 is a select agent strain of RVFV and was handled within the Saint Louis University Virus inhibition assays and toxicity assays, described below, were completed in both Vero E6 and A549 cells (ATCC® CCL-185™). Unless otherwise specified, all cells were cultured in Dulbecco’s Modified Eagle Medium (Sigma-D5796-500ML) containing 1% HEPES (Sigma-H3537-100ML) and 5% FBS (Sigma-F0926) at 37°C, 5% CO2. Studies with infectious RVFV-ZH501 viruses were approved by the SLU IBC and were conducted in our select agent registered BSL-3 laboratory.

### 2.3. Focus Forming Assay (FFA)

FFA assays are used to quantify infectious virus and are the basis for the antiviral compound inhibition assay. Briefly, 100ul of Vero E6 cells at a concentration of 3×105 cells/ml were plated in a 96-well flat bottom plate resulting in a confluency of 90-95% the day prior to the assay. To quantify viral stocks, tenfold serial dilutions of virus supernatants were then made in a 96-well round bottom plate containing 5% DMEM media before being added to the Vero cell monolayer and allowed to adsorb for one hour in an incubator with 37°C, 5% CO2. Following virus adsorption, a solution of 2% methylcellulose (Sigma-M0512-250G) was diluted 1:1 in 5% DMEM and warmed to room temperature. The methylcellulose-media mixture overlay was added to the plate by adding 125 µL of overlay media to each well and returned to an incubator with 37°C, 5% CO2 for 24 hours. Plates were then fixed in a solution of 5% paraformaldehyde (PFA) diluted in tissue culture grade 1X PBS, then washed in 1X PBS for 15 minutes. Foci were visualized by an immunostaining protocol using anti-nucleocapsid protein antibody (1D8) diluted 1:5000 to detect RVFV and anti-Gc protein antibody (4C12A1) diluted 1:5000 to detect LACV with FFA staining buffer (1X PBS, 1mg/ml saponin (Sigma: 47036)) as a primary detection antibody overnight at 4°C. The anti-nucleocapsid protein antibody (1D8) was obtained from Joel Dalrymple and Clarence J Peters (USAMRIID) via BEI resources. The secondary antibody consisted of goat anti-mouse conjugated horseradish peroxidase (Sigma: A-7289) diluted 1:5,000 in FFA staining buffer and allowed to incubate for 2 hours at room temperature. Foci were visualized using KPL TrueBlue HRP substrate and allowed to develop for 10-15 minutes, or until blue foci are visible. The reaction was then quenched by washing with Millipore water. RVFV ZH501 foci assays were measured within the BSL3 facility, while RVFV MP12 and LACV viral foci were quantified with an automated ELISPOT machine (CTL universal S6) using the Immunospot software suite.

### 2.4 Antiviral compound efficacy assay

RVFV (MP-12 and ZH501)-Vero cells were plated at 3×105 cells per mL in a flat bottom 96 well plate. Twenty-four hours later they were infected with RVFV at a multiplicity of infection of 0.005. Compound was added and plates were incubated for 1 hour at 37°C and then cells were overlayed with methylcellulose. After 24 hours plates were fixed, RVFV infectious foci were stained and quantified as described for the FFA above.

RVFV (MP-12)-A549 cells, a human lung epithelial cell line, were plated at 3×105 cells per mL in 96 well plates and incubated at 37°C for 24 hours. Cells were then infected with RVFV strain MP-12 at a multiplicity of infection of 0.005. Plates were incubated for 1 hour at 37°C and then cells were overlayed with methylcellulose. After 24 hours plates were fixed, RVFV infectious foci were stained and quantified as described for the FFA above.

LACV (original) Vero cells were plated at 2×105 cells per mL in a flat bottom 96 well plate. Twenty-four hours later they were infected with LACV at a multiplicity of infection of 0.01. Compound was added and plates were incubated for 1 hour at 37°C and then cells were overlayed with methylcellulose. After 24 hours plates were fixed, LACV infectious foci were stained using the murine anti-LACV Gc antibody 4C12A1, detected with an anti-mouse HRP secondary antibody and quantified as described for the FFA above.

Fifty percent effective concentrations (EC50) for key hits were determined by screening for suppression of viral growth using an eight point, 2.5-fold dilution series of the compounds starting at 100 µM. EC50 values were calculated by non-linear curve fitting in GraphPad Prism v7.

### 2.5. Compound Cytotoxicity

Initial compound cytotoxicity was estimated in Vero cells in the primary antiviral compound assays by staining the virally infected, compound treated cells were stained with crystal violet after viral foci had been counted. After cells were stained with crystal violet wells were washed 2x with water. Then the crystal violet dye was extracted using 50% ethanol and absorbance was measured at 570nm in an ELISA plate reader. Estimated 50% cytotoxic concentration (CC50) values were derived by non-linear curve fitting in GraphPad Prism of the four data points derived from the primary screen.

Quantitative CC50 values were measured in two systems. Compound cytotoxicity in A549 cells was determined using the CytoTox-GloTM Cytotoxicity Assay (Promega) according to the manufacturer’s instructions. Briefly, A549s were seeded at 3×105 cells per mL in 96 well plates and incubated for 24 hours at 37°C. An eight point, 3-fold dilution series of compounds was added to the cellular monolayer starting with the highest concentration, 600 μM, in addition to DMSO and PBS as controls. Plates were incubated for 48 hours at 37°C, then the AAF-GloTM Reagent was added to the cells for 15 minutes and luminescence was measured to determine the number of dead cells. Lysis buffer was added next for 15 minutes, then luminescence was measured to determine the total cell number. The dead cell number was then subtracted from the total cell number to generate the viable cell number.

Second, cytotoxicity was measured in the HepG2-derived hepatoblastoma cell line HepDES19 [32]. Cells were treated with a range of compound concentrations in a final DMSO concentration of 1% for three days and mitochondrial function was measured by MTS assays as described [33]. CC50 values were then calculated by non-linear curve fitting in GraphPad Prism v8.

### 2.6 RNaseH inhibition reactions

Activity of human RNaseH was measured using a FRET assay in which the RNA:DNA heteroduplex was formed by annealing an 18 nucleotide-long RNA oligonucleotide with a fluorescein label at the 3’ end to a complementary DNA oligonucleotide with an Iowa Black quencher at the 5’ end. RNaseH activity cleaves the RNA, permitting the fluorescein diffuse away from the quencher, increasing fluorescence. The oligonucleotides employed were:

DNA: 5’-IABkFQ-AGC TCC CAG GCT CAG ATC-3’ (IABkFQ: Iowa Black quencher)

RNA: 5’-GAU CUG AGC CUG GGA GCU FAM-3’ (FAM: Fluorescein fluorophore). Recombinant human RNaseH 1 was purified from E. coli as described in [34].

Enzyme and substrate (12.5 nM) were combined in 100 mM NaCl, 50 mM HEPES pH 8.0, 2 U RNase OUT (ThermoFisher), and test compound in a final concentration of 1% DMSO. Reactions were started by adding MgCl2 to 5 mM and incubating at 37oC for 90 min. with detection of fluorescence every 2 min. in a plate reader. The initial rate was determined for each compound concentration, and IC50 values were determined from the reaction rates by non-linear curve fitting in GraphPad Prism.

## 3. Results

### 3.1. Development and validation of a live virus Bunyavirus antiviral compound efficacy assay

Based on our previous work optimizing a focus forming assay (FFA) for Zika virus and SARS-CoV-2, [35-37] we developed an antiviral compound efficacy assay for RVFV and LACV. We chose to develop a FFA because it is a high-throughput cell-based assay that can capture a range of information about viral growth and replication such as number of infection foci and foci morphology, critical information for the development of antiviral compounds. To establish and validate the assay, we needed to identify an appropriate cell line, define primary antibodies capable of detecting viral antigen, and determine an optimal cell concentration. Previous work from a number of laboratories demonstrated that Vero E6 cells, a non-human primate African green monkey kidney epithelial cell line, was highly sensitive to RVFV infection and are widely available [38].

To identify an appropriate primary antibody to detect viral antigen, we stained a serial dilution of virally infected Vero E6 cells. We evaluated 5 total monoclonal antibodies, with 2 antibodies recognizing the Gc glycoprotein (4B6, 3D11), 2 antibodies recognizing the Gn glycoprotein (7B6, 3C10), and one antibody recognizing the nucleocapsid protein (1D8) from the Joel M. Dalrymple-Clarence J. Peters USAMRIID antibody collection (Figure 1A). The murine anti-flavivirus mAb 4G2 was used as a negative control. The mAb 1D8 had the best signal to noise ratio and was used throughout the rest of these studies.

**Figure 1.**
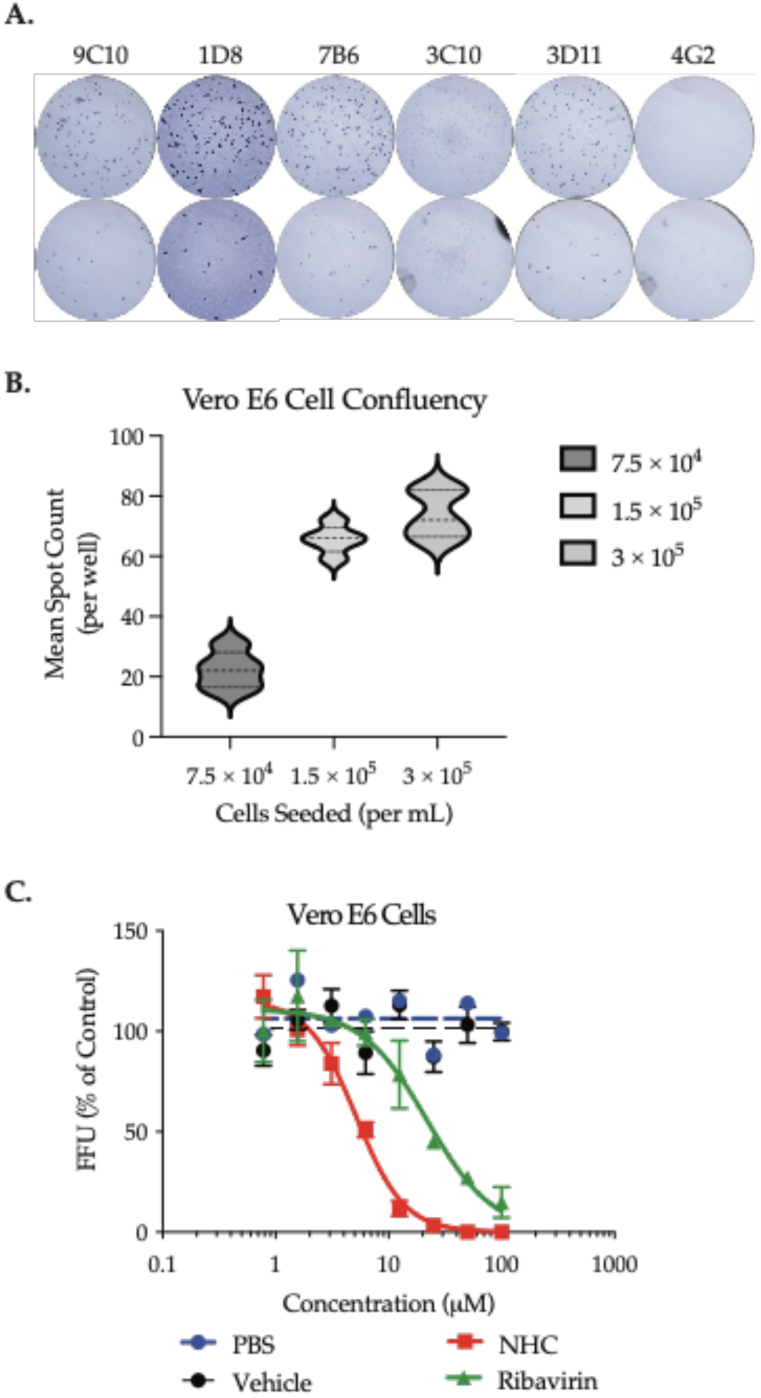
Development and validation of RVFV antiviral screen. **A**. Identification of optimal monoclonal antibodies (mAb) for the detection of RVFV by mAb staining of a serial dilution of RVFV strain MP-12 in a FFA. **B**. Impact of cell number on the sensitivity of antiviral compound screen. **C**. Evaluation of the sensitivity of the antiviral compound screen based upon the evaluation of ribavirin, a known antiviral for RVFV. The data is the cumulation of three independent experiments with technical duplicates.

**Figure 2.**
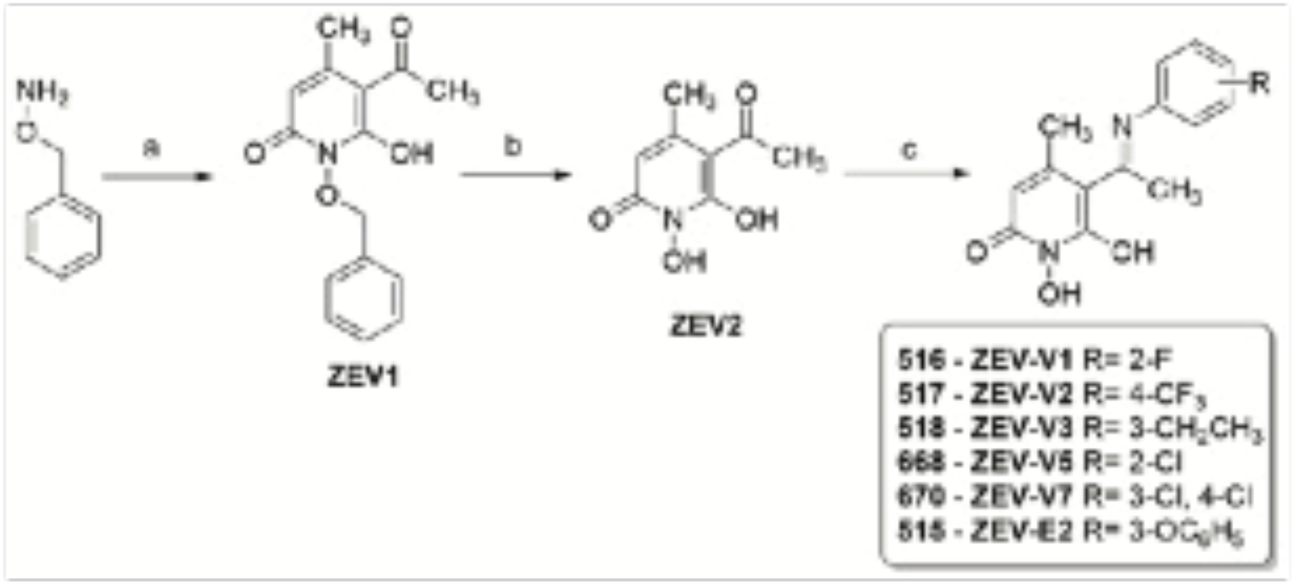
Generation of novel *N-Hydroxypyridinedione* compounds based upon ZEV1 (516). General synthetic route of HPDs, reagents and conditions; *(a) diketene*, TEA (1.0 eq), dry toluene, 4.5 h, 65 °C, Ar; (*b*) H_2_, Pd/C (10%), 40 psi, 20 min, rt; (*c*) substituted aniline, conc. H_2_SO_4_, EtOH, 4-24 h, 60 °C, Ar.

In order for RVFV to form distinct foci and for the assay to have the highest level of sensitivity, it is critical to plate cells at an optimal density. We examined the impact of cell density on foci formation by plating identical dilutions of RVFV virus stocks on 96-well plates seeded with differing numbers of E6 cells (7.5 × 10^4^, 1.5 × 10^5^ or 3 × 10^5^ cells/mL). At these concentrations, the monolayers were ∼70, 80 and 90 percent confluent, respectively, and the same virus dilution resulted in 1 × 10^3^ FFU/mL of RVFV MP-12. We observed the highest sensitivity when either 1.5 × 10^5^ or 3 × 10^5^ cells/mL were seeded in comparison to 1 × 10^5^ cells/mL (Figure 1B). We have previously tested higher cell densities for FFAs measuring flavivirus and coronavirus replication, and we have noted that cell concentrations higher than 3 × 10^5^ cells/mL results in an overly confluent monolayer with more cells than can adhere to the wells, which can lead to highly variable titer information (data not shown).

To validate the assay design, we completed an antiviral compound inhibition assay by plating Vero E6 cells at 3×105 cells/ml in a 96 well flat bottom plate, infected wells with sufficient virus to form ∼70-80 foci per well and treated cells with serial dilutions of ribavirin, or β-D-N4-Hydroxycytidine N4-Hydroxycytidine (NHC/EIDD-1931) starting at 100uM as positive controls, 100uM DMSO as a vehicle control, PBS or untreated as negative controls. The ratio of foci in comparison to vehicle control was completed and the effective concentration 50 (EC50) ribavirin had an EC50 of 22.0 µM, while NHC was 5.16 µM (Figure 1C). The resulting EC50 for ribavirin is similar to work by other groups [39,40].

### 3.2 Primary screening

Primary screens were conducted that evaluated 397 compounds either with known metal-chelating motifs or motifs similar to metal-chelating ones. The most common chemotype among the compounds screened was the troponoids (tropones, tropolones (TRP), thiotropolones (TTP), and α-hydroxytropolones (αHTs)), but the compound set also included a wide range of other chemotypes such as the N-hydroxypyridinediones (HPD), flavonoids, N-hydroxynapthyridinones, dihydronapthalenes (DHN), dioxobutanoic acids, hydroxyxanthanones, thienopyrimidinones, pyridinepiperazinthieonpyrimidins, N-biphenyltrihydroxybenzamides, and aminocyanothiophenes. Almost half of the compounds screened were αHTs as the library from which they were drawn was assembled in support of anti-HBV ribonuclease H screens and the αHTs are a leading chemotype in that effort [26,41,42].

Cells were infected with RVFV MP-12, treated with 60, 20, 6.7, or 2.2 µM of compound, and the number of RVFV foci was determined 24 hours later. Antiviral efficacy was calculated as an estimated 50% effective concentration (EC50) from the number of RVFV foci detected in comparison to vehicle control (Supplemental Table 1). Following detection of the RVFV foci, cytotoxicity concentrations (CC50) were estimated by qualitatively assessing monolayer integrity by staining the cell monolayers with crystal violet and measuring the optical density. Screening hits were defined as compounds that i) had an estimated EC50 determined from the four-point screening assay of <20 µM and ii) by measuring monolayer integrity using crystal violet, as commonly done for cell based bioassays (Table 1).

Forty-seven screening hits were identified (Supplemental Table 1). Forty of 174 troponoids screened (22%) were hits, with 35 of them being αHTs, four being TRPs, and two being TTPs. In contrast, only eight of 223 non-troponoids (3.6%) were hits. Seven of these eight hits were HPDs. The hit rate among the 24 HPDs screened (29%) was similar to that among the αHTs. The remaining screening hit was a DHN.

### 3.3 Synthesis of novel N-hydroxypyridinediones

Based on the positive primary screening results of compound 516 (ZEV-V1) and in order to provide greater insight into the structural features that may enhance the activity of this compound, we synthesized five novel HPD analogues bearing different electron withdrawing or electron donating groups on different positions on the aromatic ring (Figure 2, Scheme 1).

The key intermediate for the synthesis of the target compounds, 5-acetyl-1-(benzyloxy)-6-hydroxy-4-methylpyridin-2(1H)-one (ZEV1), was obtained by refluxing a mixture of O-benzyl hydroxylamine (1 eq) and diketene (2 eq) in the presence of triethylamine (1 eq) in dry toluene. Subsequently, the benzyl group was cleaved by catalytic hydrogenation over 10% palladium on carbon to afford the target compound ZEV2 almost quantitatively. 5-Acetyl-1,6-dihydroxy-4-methylpyridin-2(1H)-one (ZEV2) was coupled with the appropriate substituted aniline using sulfuric acid as catalyst in absolute ethanol at reflux. Novel HPD compounds explored here include 515-518, 668 and 670; synthetic details and structure elucidation data is in Supplemental Data File 1. Structures of new compounds are in Supplemental Data File 2.

### 3.3 Secondary screen of compounds against RVFV

Based upon the activity of the primary screen of αHTs and HPDs and because these compound families’ antiviral activity against RVFV is novel, we defined the dose-response curve for 35 αHTs, and 6 HPDs, and 6 additional compounds selected to broaden the chemical diversity assessed. This also served to spot-check compounds with poorer estimated EC50 values. In these studies, Vero cells were treated with a range of compound concentrations from 60 to 0.024 µM at the time of infection and their ability to prevent virus replication was measured by comparing the number of viral infection foci to wells treated with vehicle control. EC50 ranged from 1.6 to >120 µM (Supplemental Table 2). With 24 of the top 25 compounds belonging to the αHTs and one compound belonging to the HPDs. Compound AG-II-18-P (308) is a thiophene substituted αHT which had an EC50 of 1.2 µM, while the closely related furan counterpart (309) had an EC50 of 1.6 µM.

To better understand the potential cellular cytotoxicity in the context of infection, after foci were enumerated to determine the EC50 values, crystal violet staining of the cell monolayers was quantified by absorbance using a plate reader. The CC50 ranged from 22.1 to >120 µM in Vero cells, with 37 compounds having a CC50 of >120 µM. Second, cytotoxicity was assessed in the HepG2 derivative HepDES19 [32] to model longer-duration compound exposure in the liver cells. Cells were treated with a range of compound concentrations for three days and mitochondrial function was measured using MTS assays. CC50 values ranged 1.8 to >100 µM in the hepatoblastoma cells (Supplemental Table 2).

To validate compound efficacy and cytotoxicity we completed additional dose response curve experiments in A549 cells. We selected A549 cells, a human alveolar basal epithelial cell line because of the evidence that natural infection of RVFV leads to infection of epithelial cells within the kidney, liver, and spleen [43]. There was a strong concordance between the EC50 value defined in Vero cells and the EC50 concentration defined in A549 cells (Table 2 and Figure 3). In A549 cells, both compound AG-II-18-P (308), a thiophene substituted αHT, and its furan counterpart (309) had the lowest EC50 values of 8.4 and 5.9 µM, respectively. To quantify cytotoxicity independent of viral infection in the same cell line as efficacy was determined, cell permeability was quantified by measuring protease release. In these assays, A549 cells were cultured and plated as for the efficacy assays, and cells were incubated with compound for two days and cell permeability measured. For the αHTs, CC50 values ranged from 43 to >240 µM, and they ranged from 16.5 to > 240 µM for the HPDs.

**Figure 3.**
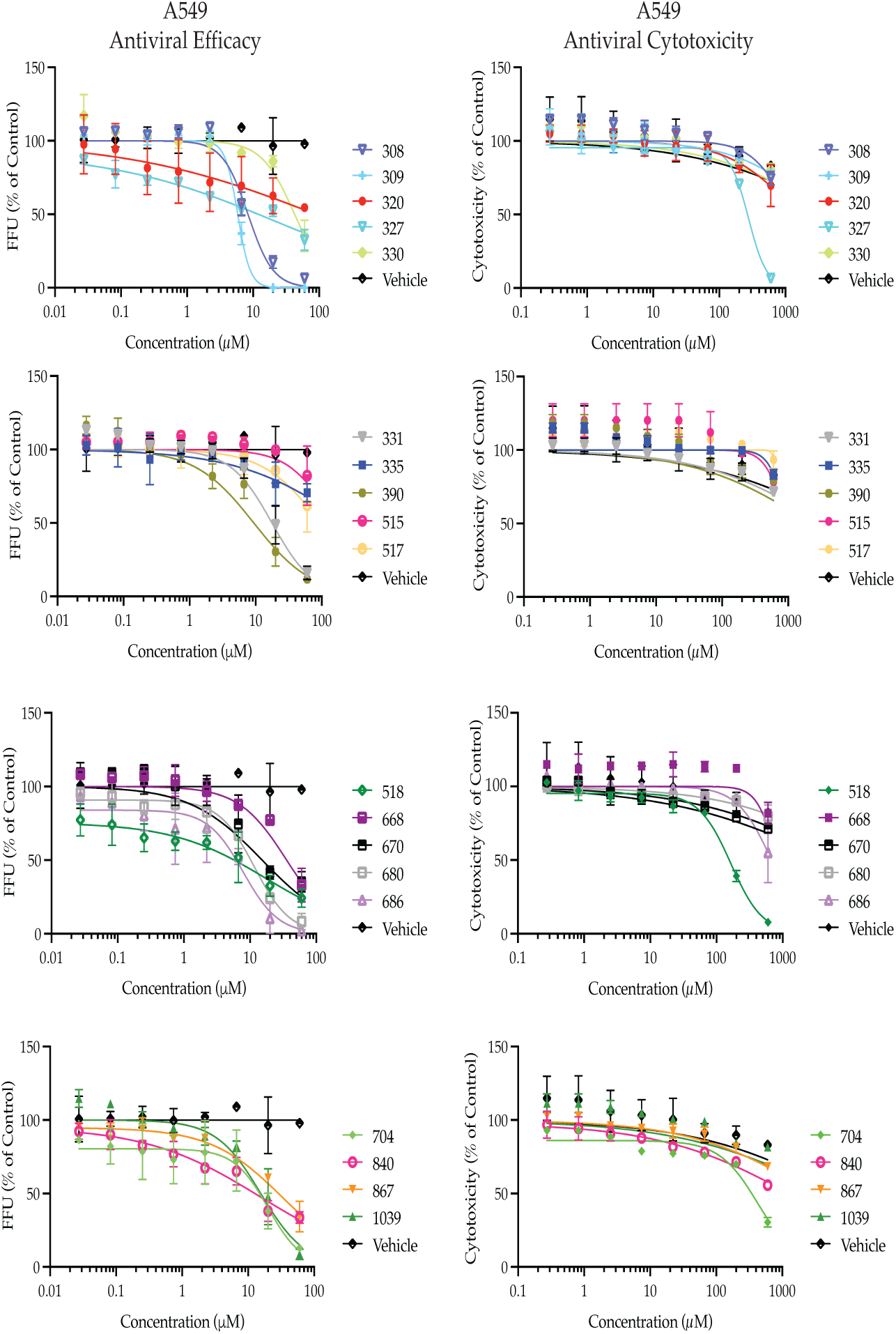
In vitro dose-response and cytotoxicity of compounds against RVFV (MP12). A549 cells were infected with RVFV MP12 then treated with decreasing concentrations of compound. The reduction in virus concentration was measured by FFA at twenty-four hours post infection. Data is representative of three individual experiments with two biological replicates. Error bars represent std dev.

In order to determine efficacy for wild type isolates of RVFV, we measured the efficacy of the αHT AG-II-18-P (308) and the nucleoside analogues NHC and ribavirin as a positive control against the highly pathogenic strain ZH501 in Vero cells (Figure 4). The assay design is identical to Figure 1, with the exception that the foci assay was fixed after 18 hours, because of the increased rate of replication. In this assay, 308 had an EC50 of 54 µM, while NHC and ribavirin had EC50s of 30.5 and 73.6 µM.

**Figure 4.**
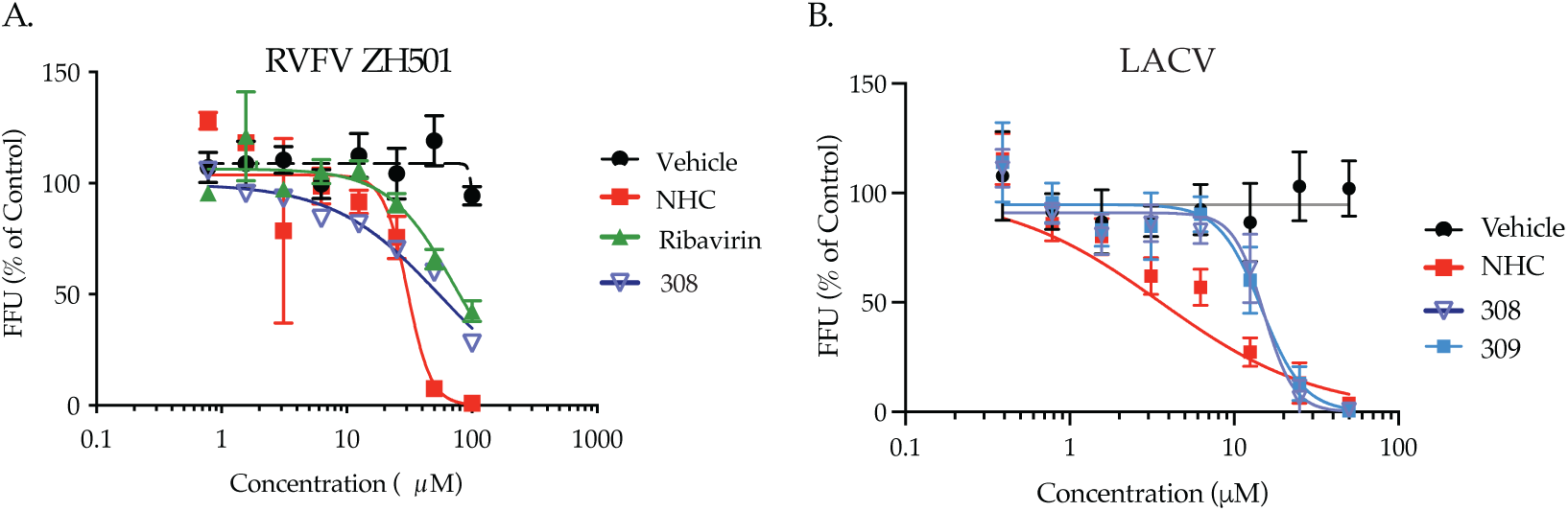
Antiviral effect of compounds on Bunyavirus replication. Vero cells were infected with either RVFV ZH501 (A) or LACV (B) then treated with decreasing concentrations of antiviral compound. Viral growth was measured by FFA. Data represents three independent experiments completed with biological replicates. Error bars represent std dev.

### 3.4 Efficacy of compounds against RVFV

To determine if the αHTs AG-II-18-P (308) and AG-I-183-P (309) and the nucleoside analogues NHC were specific to RVFV or would have a broader activity, we investigated efficacy of these compounds against LACV in Vero E6 cells. As with RVFV MP-12. we observed a dose-dependent decrease in viral titers in the antiviral efficacy assays with 308 having an EC50 of 15.1 µM, 309 had a EC50 of 14.6 µM, while NHC had EC50 of 3.6 µM. (Figure 4 and Table 3). These assays demonstrated the efficacy of these two compounds against two different bunyaviruses.

**Table 3.** EC50 and CC50 against La Crosse virus replication.

| Compound number | Compound name | Chemotype | La Crosse Vero (EC50, μM) | Vero (CC50 μM) | SI (Vero) |
| --- | --- | --- | --- | --- | --- |
| 308 | AG-II-18-P | αHT | 15.1 | 100 | 6.6 |
| 309 | AG-I-183-P | αHT | 14.6 | >120 | 8.2 |
| NHC | β-D-N4-Hydroxycytidine | Nuc | 3.6 | >120 | 33.3 |

### 3.5. Selective indexes

Selective indexes (CC50/EC50) were calculated for all compounds with EC50 values based on cytotoxicity in both Vero cells after one day compound treatment and in A549 cells after two days of treatment. SIs ranged from <1 to 402 in Vero cells and <1 to 23.9 in A549 cells (Supplemental Table 2). Confirmed hits were defined as compounds with TIs > 5 in Vero cells (n = 31) because this indicates that the reduction in RVFV foci was not a result of non-specific killing of the Vero cells in which the screening was done. Structures of all compounds in Tables 1 and 2 are in Supplemental File 2, and structures of all confirmed hits are in Supplemental File 3

## 4. Discussion

Most primary hit compounds against RVFV were either αHTs or HPDs, but TRP, TTP, and DHN hits were also found. This distribution of hits is partially due to sampling bias based on the disproportionate number of αHTs in the compound collection screened. Sampling bias, however, does not fully explain the hit distribution because only a minority of the αHTs screened were active, and because there were many chemotypes in the compound collection where hits were not found. These included dioxobutanoic acids, hydroxyxanthanones, thienopyrimidinones, pyridinepiperazinthieonpyrimidins, N-biphenyltrihydroxybenzamides, and aminocyanothiophenes. EC50 values of the 47 compounds for which quantitative data were obtained ranged from 1.16 to >120 µM (Table 2). Selective indexes for these compounds in Vero cells in which the screening was conducted ranged from 1 to 402 (average = 21). Thirty-one compounds were confirmed hits (TIs > 5 in Vero cells, 8.8% of the compounds screened) indicating that they were due to bona fide inhibition of RVFV rather than secondary effects of cytotoxicity. However, the increased CC50 values in the A549 (16.5 to >240) and HepDES19 cells (<1 to 48) indicate that compounds active versus RVFV replication can have cytotoxicity in human lung and liver-derived cells that must be addressed during subsequent hit-to-lead medicinal chemistry campaigns. One avenue for reducing cytotoxicity, at least for the αHTs, may be to reduce the lipophilicity and number of aromatic rings in the molecules because these parameters correlate with αHT toxicity [26]. RVFV infections proceed rapidly in vivo, so optimizing these hits to achieve toxicity profiles suitable for a one- to two-week treatment regimen in RVFV-infected patients or animals will likely be enough to yield usable drugs.

Identification of primary screen hits among the αHT, HPD, TRP, TTP, and DHN chemotypes indicates that a range of compounds can inhibit RVFV replication, but the lack of hits among the other chemotypes screened implies that there is specificity to RVFV inhibition. This is further supported by the efficacy of the αHT, HPD classes of compounds against LACV. The potential for specific inhibition of RVFV was confirmed by the wide range of inhibition patterns observed during counter-screening (Supplemental Table 2). For example, 362 had an EC50 of 2.6 µM vs. RVFV, an IC50 against human ribonuclease H1 of 212 µM, was inactive against E. coli growth, and was modestly effective against C. neoformans (MIC80 = 24 µM). In contrast, 260 was poorly active or inactive in all of the counter-screens, and 389 had good to moderate activity against all microbes screened, but was nearly inactive against human RNase H1. This indicates that although these chemotypes can have broad anti-microbial activity, individual compounds can be selective through specific interactions with their various targets.

The 174 troponoids (αHTs, TRPs, TTPs) evaluated yielded 28 confirmed hits with TI values ≥ 5 (Supplemental Table 2 and Supplemental File 3). Inhibition of RFVF by tropolones appears to require an intact metal ion chelating trident on the compound, which implies an interaction with two closely-spaced divalent cations on the target molecule due to the compounds’ known mechanisms of metal chelation[44]. For example, while the tropolone natural products β- and γ-thujaplicin were inactive against the virus at concentrations upwards of 100 µM, the αHT natural product β–thujaplicinol was active, with an EC50 of 13.8 µM. These trends extended to synthetic αHTs, of which 13 different molecules had EC50 values under 10 µM, the most potent of which had an EC50 of 0.54 µM (308). These more potent molecules had a broad range of appendages, such as ketone (308, 362, 359, 358, 330), amide (1017, 867, 388, 389, 1039), thioether (694, 696), sulfoxide (336) and aryl (330), and also included a 3,7-dihydroxytropolone (362), demonstrating tolerance to a variety of functional groups. Apart from the αHTs, three additional tropolones had measurable EC50 values (the TTPs 340, 341, and 342, EC50 = 25-55 µM), each of which had a carbonyl appendage α to the tropolone oxygens that could provide an alternative third contact point.

Seven HPD primary hits were found among the 24 HPDs screened. Three of these were confirmed hits, 518, 668, and 670, with EC50 values ranging from 14-20 µM. This is insufficient to generate a meaningful structure activity relationship, but trends can be inferred. The oxygen trident of the HPD scaffold is essential for its activity, as the loss of any one of these oxygens results in inactive compounds. In each case it is presumed based on data with other HPDs against HBV [33] that strong ionic interactions, along with charge-assisted hydrogen bonds anchor the chelator moiety of HPDs (N-Hydroxyimide group) to the RVFV endonuclease catalytic ensemble comprised of the two positively charged Mg++ ions. Lastly, aromatic substitutions at the imine nitrogen are tolerated that carry modifications including halogen electron withdrawing groups and an alkyl electron donating group.

The mechanism(s) of action of the RVFV inhibitors are unknown and could involve inhibition of viral and/or cellular proteins. However, inhibiting one or more mono- or di-metalloenzymes needed for viral replication by chelating their active site cations is a likely mechanism. The rationale for implicating metal chelation comes from the compound structures and their known activities against HIV and presumed activity against HBV [10-12]. The αHTs and HPDs have metal chelating tridents suitable for binding to the Mg++ ions in di-metalloenzyme active sites, and the αHTs are known to work by this mechanism against the HIV ribonuclease H and/or integrase [11,15]. However, the failure of many compounds with known ability to inhibit divalent cation containing metalloenzymes (∼70% of the αHTs did not inhibit RVFV growth) indicates that metal chelation by itself is insufficient, presumably because additional compound:target interactions are needed to provide sufficient binding affinity to inhibit viral replication. One potential target for these inhibitors is the RVFV cap-snatching endonuclease activity of the viral L protein [6,7]. This is because it is a di-metalloenzyme that catalyzes a reaction similar to that of the cap-snatching endonuclease PA endonuclease from influenza virus.

## 5. Conclusions

Screening for RVFV replication inhibitors among compounds selected for their similarity to inhibitors of viral nucleases identified 31 novel RVFV inhibitors. The frequent efficacy of the αHT and HPD compounds screened against RVFV replication indicates that these two scaffolds are promising candidates for optimization into anti-RVFV drugs for use against human and/or veterinary infections. The mechanism(s) of action for these inhibitors are not known, but it is likely one target would be the RVFV cap-snatching endonuclease. Cytotoxicity was observed in human hepatoblastoma cells, indicating that identifying and mitigating the causes of cytotoxicity will be key to optimizing these hits. The conservation of the cap-snatching mechanism among Bunyavirales and the activity of these compounds versus LACV implies that these hits hold potential for development into treatments for related pathogens, including Hantaan Virus, Severe fever with thrombocytopenia syndrome virus, and Crimean-Congo Hemorrhagic Fever Virus.

## 6. Patents

AP, GZ, JB, JT, and RM are inventors on a pending US patent application covering use of these compounds to treat Bunyavirus infections.

Supplementary Materials: The following supporting information can be downloaded at: www.mdpi.com/xxx/s1, Supplemental Table 1: Primary screening data; Supplemental Table 2: Quantitative analysis of RVFV screen; Supplemental Data File 1: Experimental methods for compound generation and validation of compounds; Supplemental Data file 1: Structures of all compounds in Tables 1 and 2.; Supplemental Data file 2: Structures of all confirmed hit compounds.

### Author Contributions

Conceptualization, J.T., A.O.D., J.D.B, and A.K.P.; methodology, E.G., E.T.S., V.M., M.H., A.O.D., F.C., M.D., M.E., V.P., E.G., S. S. A.B., N.A.; formal analysis, E.G., J.T., A.O.D., J.B.; investigation, A.O.D., F.C., M.D., M.E., V.P., E.G., S. S. A.B., N.A.; resources, J.T., B.E., G.Z., R.M.; data curation, J.T., M.D., F.C., G.Z.; writing—original draft preparation, E.G., J.T., J.D.B., G.Z.; writing—review and editing, E.G., J.T., J.B., R.M., G.Z., B.E.; visualization, X.X.; supervision, J.D.B., J.T., B.E., G.Z., R.M., A.K.P.; project administration, J.T., J.B.,A.K.P.; funding acquisition, J.T., J.D.B., A.K.P., G.Z. R.M. All authors have read and agreed to the published version of the manuscript.

### Funding

This work was funded by the USA National Institutes of Health [grants R01 AI1222669, R01 AI148264, NIH R21 AI124672, and SC GM111158], the Special Account for Research Grants from the National and Kapodistrian University of Athens [grant 15725], and a seed grants from the School of Medicine and the Department of Molecular Microbiology and Immunology, Saint Louis University School of Medicine. Institutional Review Board Statement: Not applicable Informed Consent Statement: Not applicable

### Data Availability Statement

All data are either presented in this manuscript and its supplementary data or are available upon request.

## Acknowledgments

We thank Qilan Li, Brienna Milleson, Alaina Knier, and Elena Lomonosova for technical assistance.

