## Supplemental files for "Metal Coordinating Inhibitors of Rift Valley Fever virus Replication"

**Supplemental Table 1. Primary screening hits against RVFV replication and select non-hits for comparison**

| Compound number | Name [1] | Chemotype [2] | Estimated $EC_{50}$ ($\mu$M) | Estimated $CC_{50}$ ($\mu$M)3 | Preliminary SI | Chemist |
|---|---|---|---|---|---|---|
| **α-Hydroxytropolones** | | | | | | |
| 390 | AB-2-70 | αHT | 0.1 | | | Murelli |
| 308 | AG-II-18-P | αHT | 2 | 111 | 63 | Murelli |
| 320 | NBA-I-13 | αHT | 2 | | | Murelli |
| 330 | NBA-I-14 | αHT | 2 | | | Murelli |
| 331 | NBA-I-31 | αHT | 2 | | | Murelli |
| 335 | DH-2-60 | αHT | 3 | | | Murelli |
| 336 | YA-I-78 | αHT | 3 | | | Murelli |
| 311 | AG-II-3-P | αHT | 3 | 115 | 36 | Murelli |
| 1017 | AL-23 | αHT | 3 | >120 | 40 | Murelli |
| 359 | AG-II-108-C | TRP | 4 | | | Murelli |
| 694 | NBA-I-127 Bis | αHT | 6 | >120 | | Murelli |
| 362 | DS-I-69 | αHT | 6 | | | Murelli |
| 1039 | AB-3-45 | αHT | 6 | >120 | | Murelli |
| 867 | DS-1-124 | αHT | 7 | >120 | | Murelli |
| 702 | NBA-I-159 Mono | αHT | 7 | >120 | | Murelli |
| 210 | MolMoll 19617 | αHT | 8 | >120 | | Purchased |
| 698 | NBA-I-150 | αHT | 8 | >120 | | Murelli |
| 838 | NBA-I-130 | αHT | 8 | 86 | 10 | Murelli |
| 703 | NBA-I-159 Bis | αHT | 9 | >120 | 13 | Murelli |
| 539 | AB-2-91 | αHT | 10 | 71 | 7 | Murelli |
| 696 | NBA-I-128 Bis | αHT | 12 | 118 | 9 | Murelli |
| 836 | RA-I-86 | αHT | 13 | >120 | 9 | Murelli |
| 704 | NBA-I-160 | αHT | 14 | >120 | 8 | Murelli |
| 840 | NBA-I-155-Mono | αHT | 14 | >120 | 8 | Murelli |
| 389 | AB-2-66 | αHT | 16 | | | Murelli |
| 711 | JS-116 | αHT | 17 | >120 | 7 | Murelli |
| 710 | JS-112 | αHT | 17 | >120 | 7 | Murelli |
| 809 | RA-I-82 | αHT | 17 | 64 | 3 | Murelli |
| 799 | AJF-1.033 | αHT | 17 | 62 | 3 | Murelli |
| 876 | JS-166 | αHT | 19 | >120 | 6 | Murelli |
| 1019 | AL-22 | αHT | 19 | >120 | 6 | Murelli |
| 388 | RA-1-30 | αHT | 19 | | | Murelli |
| 234 | AB-1-51 | αHT | 19 | >120 | 6 | Murelli |
| 920 | RA-1-104 | αHT | 19 | >120 | 6 | Murelli |
| **Thiotropolones** | | | | | | |
| 680 | BE1105 | TTP | 0.3 | 0.9 | 3 | Elgendy |
| 686 | BE1111 | TTP | 2 | >120 | | Elgendy |
| **Tropolones** | | | | | | |
| 341 | Specs AP-355/40802214 | TRP | 2 | | | Purchased |
| 342 | Specs AP-355/40633885 | TRP | 2 | | | Purchased |
| 340 | Specs AP-355/40633884 | TRP | 3 | | | Purchased |
| **Dihydronapthalene** | | | | | | |
| 327 | Aldrich Select CNC_ID 444085867 | DHN | 2 | | | Purchased |
| **N-Hydroxypyridinediones** | | | | | | |
| 518 | ZEV-V3 | HPD | 7 | >120 | | Zoidis |
| 670 | ZEV-V7 | HPD | 10 | >120 | | Zoidis |
| 668 | ZEV-V5 | HPD | 11 | >120 | | Zoidis |
| 517 | ZEV-V2 | HPD | 13 | >120 | | Zoidis |
| 208 | Sun B8155 | HPD | 14 | >120 | | Purchased |
| 515 | ZEV-E2 | HPD | 14 | >120 | | Zoidis |
| 516 | ZEV-V1 | HPD | 19 | >120 | | Zoidis |
| **Nucleoside Analogue** | | | | | | |
| EIDD-1931 | β-D-N4-HydroxycytidineN4-Hydroxycytidine | Nuc | 5.16 | >120 | | Purchased |
| | Ribavirin (Cayman Chemical) | Nuc | 22 | >120 | | Purchased |
| **Example inactive compounds** | | | | | | |
| 6 | Chembridge 7929959 | ACT | >120 | | | Purchased |
| 7 | Quercetagetin | FLV | >120 | | | Purchased |
| 8 | Sigma S439274 | HXT | >120 | | | Purchased |
| 22 | Sigma n8164 | TPD | >120 | | | Purchased |
| 47 | β-thujaplicin | TRP | >120 | | | NCI DTP |
| 48 | γ -thujaplicin | TRP | >120 | | | NCI DTP |
| 129 | Aldrichselect CNC_ID 249465147 | DOB | >120 | | | Purchased |
| 138 | Sigma H53704 | HPD-like | >120 | | | Purchased |
| 522 | ZAU5 | HPD-like | >1 | 1 | <1 | Zoidis |
| 681 | BE1106 | TTP | >2 | 2 | <2 | Elgendy |
| 700 | NBA-I-157 Bis | αHT | >120 | >120 | >1 | Murelli |
| 712 | JS-108 | αHT | >20 | 20 | <1 | Murelli |

1 Chemist's name, common name, or vendor catalog number.

[2] αHT, α-Hydroxytropolone; TRP, tropolone; TTP, thiotropolone; DHN, dihydronapthalene; HPD, N-Hydroxypyridinedione; FLV, flavenoid; DOB, dioxobutanoic acid; HXT, hydroxyxanthanone; TPD, thieopyrimidinone; ACT, aminocyanothiophene.

[3] Values of 120 indicate the data were at or above the upper limit of quantificaiton in the assay.

**Supplemental Table 2. $EC_{50}$s against RVFV replication**

| | | | | $CC_{50}$ | | SI | | Counter screens | | | |
|---|---|---|---|---|---|---|---|---|---|---|---|
| Compound number | Compound name | Chemotype | RVFV ($EC_{50}$, μM) | Vero (μM) | HepDES19 (μM) | Vero | HepDES19 | Human RNaseH1 ($IC_{50}$, μM) | *Staphylococcus spp.* ($MIC_{80}$, μM) | *E. coli* ($MIC_{80}$, μM) | *C. neoformans* ($MIC_{80}$, μM) |
| 308 | AG-II-18-P | αHT | 1.16 | >120 | 25.8 | 103 | 22.2 | 46 | 44 | 25 | 11 |
| 309 | | αHT | 1.6 | >120 | | 75 | | | | | |
| 362 | DS-I-69 | αHT | 2.6 | 77.8 | 5.4 | 29.9 | 2.1 | 212 | | - | 24 |
| 694 | NBA-I-127 Bis | αHT | 3.7 | 22.1 | 1.8 | 6.0 | <1.0 | >500 | 9 | - | 2.3 |
| 359 | AG-II-108-C | αHT | 5.1 | >120 | 19.4 | 23.5 | 3.8 | 65 | - | - | 24 |
| 696 | NBA-I-128 Bis | αHT | 6.0 | 56.5 | 14.1 | 9.4 | 2.4 | >500 | 13 | 20 | 1.5 |
| 1017 | AL-23 | αHT | 8.0 | 54.7 | 3.9 | 6.8 | <1.0 | 30 | - | - | 6.0 |
| 867 | DS-1-124 | αHT | 8.7 | >120 | 5.2 | 13.8 | <1.0 | >500 | - | - | 9.0 |
| 389 | AB-2-66 | αHT | 8.8 | >120 | 26.5 | 13.6 | 3.0 | 309 | 44 | 44 | 3.5 |
| 702 | NBA-I-159 Mono | αHT | 8.8 | 35.6 | 17.4 | 4.0 | 2.0 | 34 | 13 | 9 | 0.55 |
| 1039 | AB-3-45 | αHT | 8.9 | >120 | 8.7 | 13.5 | <1.0 | 209 | - | - | 12 |
| 336 | YA-I-78 | αHT | 9.0 | 65.8 | 45.4 | 7.3 | 5.0 | 85 | - | - | 50 |
| 358 | AG-II-83-P | αHT | 9.0 | >120 | 12.5 | 13.3 | 1.4 | 43 | | 20 | 11 |
| 388 | RA-1-30 | αHT | 9.0 | >120 | 19.8 | 13.3 | 2.2 | 166 | 67 | 67 | 11 |
| 113 | RM-YM-1-0613 | αHT | 10.3 | >120 | 47.0 | 11.7 | 4.6 | 174 | - | - | 24 |
| 330 | NBA-I-14 | αHT | 10.4 | >120 | 56.2 | 11.5 | 5.4 | 476 | 44 | - | 4.0 |
| 118 | RM-MD-2-0813 | αHT | 11.0 | >120 | 17.0 | 10.9 | 1.5 | >500 | - | - | 15 |
| 698 | NBA-I-150 | αHT | 11.7 | 42.9 | 13.7 | 3.7 | 1.2 | 261 | 20 | 44 | 38 |
| 210 | MolMoll 19617 | αHT | 11.8 | >120 | 71.0 | 10.2 | 6.0 | 85 | - | - | 15 |
| 311 | AG-II-3-P | αHT | 11.8 | >120 | 31.7 | 10.2 | 2.7 | 26 | 67 | 30 | 15 |
| 390 | AB-2-70 | αHT | 11.9 | >120 | 29.8 | 10 | 2.5 | 106 | - | - | 11 |
| 838 | NBA-I-130 | αHT | 12.7 | >120 | 18.6 | 9.4 | 1.5 | 100 | - | - | 1.0 |
| 46 | β-thujaplicinol | αHT | 13.8 | >120 | 25.0 | 8.7 | 1.8 | 58 | 20 | 44 | 8.0 |
| 670 | ZEV-V7 | HPD | 14 | >120 | 69.0 | 8.6 | 4.9 | 356 | - | - | 69 |
| 704 | NBA-I-160 | αHT | 14.3 | >120 | 21.6 | 8.4 | 1.5 | 25 | 20 | - | 1.8 |
| 265 | AG58 | αHT | 15.4 | >120 | 25.5 | 7.8 | 1.7 | 57 | 30 | - | 11 |
| 120 | RM-MD-1-0713 | αHT | 15.6 | >120 | 42.4 | 7.7 | 2.7 | 81 | - | - | 12 |
| 331 | NBA-I-31 | αHT | 15.8 | 80.4 | 36.6 | 5.1 | 2.3 | 448 | 67 | - | 11 |
| 703 | NBA-I-159 Bis | αHT | 16.1 | 53.2 | 8.2 | 3.3 | <1.0 | 61 | 13 | - | 7.0 |
| 539 | AB-2-91 | αHT | 18.6 | >120 | 3.6 | 6.5 | <1.0 | 105 | 38 | - | 4.5 |
| 111 | CM1012-6f | αHT | 19.0 | >120 | 35.5 | 6.3 | 1.9 | 48 | - | - | 36 |
| 668 | ZEV-V5 | HPD | 19.2 | >120 | 76.8 | 6.3 | 4.0 | 165 | - | - | na |
| 518 | ZEV-V3 | HPD | 20.0 | >120 | 91.4 | 6.0 | 4.6 | 436 | 67 | - | 24 |
| 196 | DH-3-37 | αHT | 23.9 | >120 | 6.0 | 5.0 | 0.3 | 408 | 67 | - | na |
| 515 | ZEV-E2 | HPD | 24.3 | >120 | 71.3 | 4.9 | 2.9 | 622 | - | - | 11 |
| 341 | Specs AP-355/40802214 | TRP | 25.2 | >120 | 17.1 | 4.8 | 0.7 | >500 | 44 | - | 24 |
| 385 | NBA-I-116A | αHT | 25.2 | >120 | 100* | 4.8 | 4.0 | 176 | - | - | 24 |
| 340 | Specs AP-355/40633884 | TRP | 27.3 | 70.0 | 18.7 | 2.6 | 0.7 | >500 | - | - | 49 |
| 260 | AG-I-84-P | αHT | 31.6 | >120 | 45.0 | 3.8 | 1.4 | 277 | - | - | 50 |
| 840 | NBA-I-155-Mono | αHT | 31.6 | >120 | 21.5 | 3.8 | <1.0 | 73 | - | - | 1.0 |
| 320 | NBA-I-13 | αHT | 33.96 | >120 | 49.5 | 3.5 | 1.5 | 37 | - | - | 49 |
| 327 | Aldrich Select CNC_ID 444085867 | DHN | 39.7 | 42.9 | 18.5 | 1.1 | 0.5 | 47 | - | - | 9.1 |
| 335 | DH-2-60 | αHT | 40.8 | >120 | 78.0 | 2.9 | 6.0 | 111 | - | - | 50 |
| 342 | Specs AP-355/40633885 | TRP | 55.5 | >120 | 30.4 | 2.2 | 0.5 | >500 | - | - | 24 |
| 208 | Sun B8155 | HPD | 120* | >120 | 14.4 | - | <1.0 | 194 | - | - | 30 |
| 517 | ZEV-V2 | HPD | 120* | >120 | 91.0 | - | <1.0 | 135 | - | - | 24 |
| 680 | BE1105 | TTP | 120* | >120 | 95.6 | - | <1.0 | 652 | 13 | 30 | 0.40 |
| 686 | BE1111 | TTP | 120* | >120 | 42.6 | - | <1.0 | >500 | 30 | - | 0.85 |

-, Inactive
na, Not available

### Table of Contents


#### I Experimental methods

#### II Preparation procedures and characterization data of compounds

#### III Copies of NMR spectra

NMR spectra of **ZEV1 ($^1$H, $^{13}$C, COSY, HSQC-DEPT, HMBC)**

NMR spectra of **ZEV2 ($^1$H, $^{13}$C, COSY, HSQC-DEPT, HMBC)**

NMR spectra of **ZEV-V1 ($^1$H, $^{13}$C, COSY, HSQC-DEPT, HMBC)**

NMR spectra of **ZEV-V2 ($^1$H, $^{13}$C, COSY, HSQC-DEPT, HMBC)**

NMR spectra of **ZEV-V3 ($^1$H, $^{13}$C, COSY, HSQC-DEPT, HMBC)**

NMR spectra of **ZEV-V5 ($^1$H, $^{13}$C, COSY, HSQC-DEPT, HMBC)**

NMR spectra of **ZEV-V7 ($^1$H, $^{13}$C, COSY, HSQC-DEPT, HMBC)**

NMR spectra of **ZEV-E2 ($^1$H, $^{13}$C, COSY, HSQC-DEPT, HMBC)**

#### I Experimental methods

During the conduct of the experimental part of the present study were used the following materials, apparatuses and techniques. Melting points were determined using a Büchi capillary apparatus and are uncorrected. NMR experiments were performed to elucidate the structure and determine the purity of the newly synthesized compounds. $^{1}H$ NMR and 2D NMR spectra (COSY, HSQC-DEPT, HMBC) were recorded on a Bruker DRX400 spectrometer (400.13 MHz, $^{1}H$ NMR) and a Bruker Ultrashield™ Plus Avance III 600 spectrometer (600.11 MHz, $^{1}H$ NMR). $^{13}C$ NMR spectra were recorded on a Bruker Avance 200 spectrometer (50.32 MHz, $^{13}C$ NMR), a Bruker DRX400 spectrometer (100.61 MHz, $^{13}C$ NMR) and a Bruker Ultrashield™ Plus Avance III 600 spectrometer (150.9 MHz, $^{13}C$ NMR). Chemical shifts *δ (delta)* are reported in parts per million (ppm) downfield from the NMR solvent, with the tetramethylsilane or solvent (DMSO-$d_6$) as internal standard. Data processing including Fourier transformation, baseline correction, phasing, peak peaking and integrations were performed using MestReNova software v.12.0.0. Splitting patterns are designated as follows: s, singlet; d, doublet; t, triplet; dd, doublet of doublets; td, triplet of doublets; m, multiplet; complex m, complex multiplet. Coupling constants (*J*) are expressed in units of Hertz (Hz). The spectra were recorded at 293 K (20 °C) unless otherwise specified. The solvent used to obtain the spectra was deuterated DMSO, DMSO-$d_6$ (quin, 2.50 ppm, $^{1}H$ NMR; septet, 39.52 ppm, $^{13}C$ NMR). Analytical thin-layer chromatography (TLC) was used to monitor the progress of the reactions, as well as to authenticate the compounds. TLCs were conducted on, precoated with normal-phase silica gel, aluminium sheets (Silica gel 60 $F_{254}$, Merck) (layer thickness 0.2 mm), precoated with reverse phase silica gel, aluminium sheets (Silica gel 60 RP-18 $F_{254}$s, Merck) and precoated aluminum oxide plates (TLC Aluminium oxide 60 $F_{254}$, neutral). Developed plates were examined under a UV light source, at wavelengths of 254 nm, or after being stained by iodine vapors. The Retention factor ($R_f$) of the newly synthesized compounds, that equals to the distance migrated over the total distance covered by the solvent, was also measured on the chromatoplates. Elemental analyses (C, H, N) were performed by the Service Central de Microanalyse at CNRS (France), and were within ±0.4% of the theoretical values. Elemental analysis results for the tested compounds correspond to >95% purity. The commercial reagents were purchased from Alfa Aesar, Sigma-Aldrich, and Merck, and were used without further purification. Solvent abbreviations: ACN, acetonitrile; AcOEt, ethyl acetate; $Et_2O$, diethyl ether; EtOH, ethanol; MeOH, methanol.

#### II Preparation procedures and characterization data of compounds

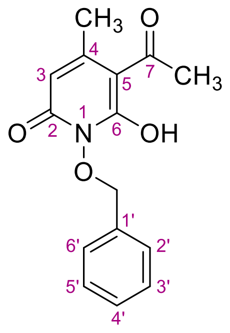

**5-acetyl-1-(benzyloxy)-6-hydroxy-4-methylpyridin-2(1*H*)-one (ZEV1)**

A stirred solution of *O*-benzylhydroxylamine (2.98 g, 24.2 mmol, 1.0 eq) and triethylamine (2.45 g, 3.38 mL, 24.2 mmol, 1.0 eq) in 19 mL dry toluene was cooled in an ice-bath, and diketene (4.07 g, 3.73 mL, 48.4 mmol, 2.0 eq) was added dropwise. After 4.5 h of stirring at 65 °C under Argon, the mixture was concentrated to dryness under reduced pressure and treated with 150 mL HCl 10%. The residue was partitioned between the aqueous phase and AcOEt (300 mL), the organic phase was extracted once more with HCl 10% (150 mL) and the combined aqueous phases were extracted once more with 150 mL AcOEt. The combined organic phases were washed with brine (3 × 200 mL), dried over anhydrous $Na_2SO_4$ and the solvent was removed in vacuo. The residual brownish solid was triturated with $Et_2O$ and AcOEt sequentially to afford the title compound **ZEV1** as a beige crystalline solid. (4.95 g, 75%). Mp 144-146 °C (MeOH, AcOEt/*n*-pentane), $R_{f\,(NP\text{-}TLC)}$ = 0.25 (AcOEt). $^1$H NMR (600.11 MHz, DMSO-$d_6$) $\delta$ (ppm) 2.34 (s, 3H, 4-C$H_3$), 2.60 (s, 3H, 7-C$H_3$), 5.05 (s, 2H, C$H_2$Ph) 5.83 (s, 1H, $H_3$), 7.37-7.44 (complex m, 3H, $H_{3'}$, $H_{4'}$, $H_{5'}$), 7.54 (dd, 2H, $J_1$=7.5 Hz, $J_2$=1.7 Hz, $H_{2'}$, $H_{6'}$); $^{13}$C NMR (50.32 MHz, DMSO-$d_6$) $\delta$ (ppm) 23.8 (4-$C$H$_3$), 29.0 (7-$C$H$_3$), 76.9 ($C$H$_2$Ph), 104.8 ($C_5$), 106.7 ($C_3$), 128.2 ($C_{3'}$, $C_{5'}$), 128.6 ($C_{4'}$), 129.3 ($C_{2'}$, $C_{6'}$), 134.9 ($C_{1'}$), 150.5 ($C_4$), 159.0 ($C_2$), 164.8 ($C_6$), 193.4 ($C_7$). Anal. Calcd for $C_{15}H_{15}NO_4$: C, 65.92; H, 5.53; N, 5.13; Found: C, 66.00; H, 5.59; N, 5.08.

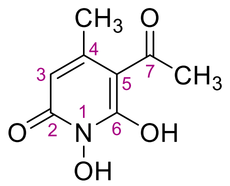

**5-acetyl-1,6-dihydroxy-4-methylpyridin-2(1*H*)-one (ZEV2)**

A solution of **ZEV1** (2.0 g, 7.32 mmol) in 120 mL MeOH was hydrogenated for 20 min at rt and 40 psi pressure, in the presence of 200 mg Pd/C (10 wt.%) as a catalyst. The catalyst was filtered off, washed with portions of hot MeOH (3 × 20 mL) and the combined filtrates were evaporated under reduced pressure. The beige crystalline product was treated with AcOEt to yield the *N*-hydroxypyridinedione **ZEV2** almost quantitatively (1.32 g, 98%). Mp 178-179 °C (MeOH/*n*-pentane, dry $Et_2O$), $R_{f\,(NP\text{-}TLC)}$ = 0.06 (AcOEt), $R_{f\,(RP\text{-}TLC)}$ = 0.85 ($H_2O$/ACN 7:3). $^1$H NMR (600.11 MHz, DMSO-$d_6$) $\delta$ (ppm) 2.32 (s, 3H, 4-C$H_3$), 2.56 (s, 3H, 7-C$H_3$), 5.80 (s, 1H, $H_3$); $^{13}$C NMR (100.61 MHz, DMSO-$d_6$) $\delta$ (ppm) 23.3 (4-$C$H$_3$), 28.0 (7-$C$H$_3$), 104.7 ($C_5$), 107.6 ($C_3$), 149.1 ($C_4$), 157.9 ($C_2$), 165.1 ($C_6$), 194.3 ($C_7$). Anal. Calcd for $C_8H_9NO_4$: C, 52.46; H, 4.95; N, 7.65; Found: C, 52.51; H, 5.00; N, 7.58.

#### 5-(1-((2-fluorophenyl)imino)ethyl)-1,6-dihydroxy-4-methylpyridin-2(1*H*)-one (ZEV-V1)

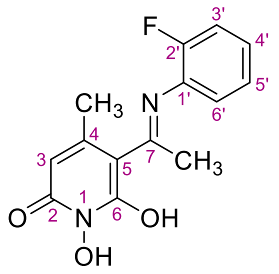

To a stirred solution of **ZEV2** (200 mg, 1.09 mmol, 1.0 eq) in absolute EtOH (4 mL) at 75 °C were added successively 2-fluoroaniline (133 mg, 116 μL, 1.2 mmol, 1.1 eq), activated 4Å molecular sieves and a drop of sulfuric acid. The mixture was refluxed for 4.5 h at 60 °C under Argon and 2.5 h at 30 °C, when a yellow solid was precipitated. The solid formed was filtered off *in vacuo* and washed with small portions of EtOH and AcOEt to afford the title compound **ZEV-V1** as a yellow solid (180 mg, 60%). Mp 190-193 °C (AcOEt/*n*-pentane), $R_{f\ (alox)}$ = 0.05 (MeOH), $R_{f\ (RP\text{-}TLC)}$ = 0.07 ($H_2O$/ACN 7:3). $^1H$ NMR (600.11 MHz, DMSO-$d_6$) $\delta$ (ppm) 2.38 (s, 3H, 4-C$H_3$), 2.46 (s, 3H, 7-C$H_3$), 5.73 (s, 1H, $H_3$), 7.30-7.37 (m, 1H, $H_{4'}$), 7.41-7.47 (complex m, 2H, $H_{3'}$, $H_{6'}$), 7.49 (t, 1H, *J*=8.0 Hz, $H_{5'}$), 10.11 (s, 1H, 1-OH), 14.78 (s, 1H, 6-OH); $^{13}C$ NMR (150.9 MHz, DMSO-$d_6$) $\delta$ (ppm) 20.4 (7-*C*$H_3$), 25.1 (4-*C*$H_3$), 99.7 ($C_5$), 110.8 ($C_3$), 116.4, 116.5 (d, $J_{C\text{-}F}$=19.5 Hz, $C_{3'}$), 124.6, 124.7 (d, $J_{C\text{-}F}$=12.2 Hz, $C_{1'}$), 125.11, 125.12 (d, $J_{C\text{-}F}$=2.8 Hz, $C_{4'}$), 128.2 ($C_{5'}$), 129.38, 129.43 (d, $J_{C\text{-}F}$=7.9 Hz, $C_{6'}$), 148.3 ($C_4$), 155.0, 156.6 (d, $J_{C\text{-}F}$=247.1 Hz, $C_{2'}$), 158.8 ($C_2$), 164.6 ($C_6$), 169.1 ($C_7$). Anal. Calcd for $C_{14}H_{13}FN_2O_3$: C, 60.87; H, 4.74; N, 10.14; Found: C, 60.91; H, 4.79; N, 10.18.

#### 5-(1-((4-(trifluoromethyl)phenyl)imino)ethyl)-1,6-dihydroxy-4-methylpyridin-2(1*H*)-one (ZEV-V2)

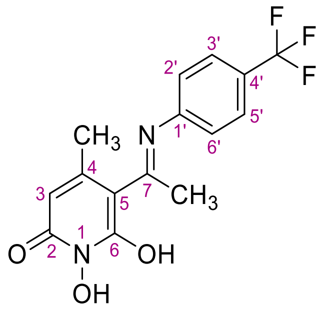

4-(trifluoromethyl)aniline (193 mg, 150 μL, 1.2 mmol, 1.1 eq) and activated 4Å molecular sieves were added to a solution of 200 mg (1.09 mmol, 1.0 eq) of **ZEV2** in 4 mL absolute EtOH at 75 °C. Then, a drop of sulfuric acid was added, and the temperature was adjusted to 60 °C. The mixture was refluxed for 4h at 60 °C under Argon. A yellow precipitate was formed, filtered off *in vacuo* and washed with EtOH to give the title compound **ZEV-V2** as a yellow crystalline solid (230 mg, 65%). Mp 191-193 °C (AcOEt/*n*-pentane), $R_{f\ (alox)}$ = 0.06 (MeOH), $R_{f\ (RP\text{-}TLC)}$ = 0.02 ($H_2O$/ACN 7:3). $^1H$ NMR (600.11 MHz, DMSO-$d_6$) $\delta$ (ppm) 2.38 (s, 3H, 4-C$H_3$), 2.51 (s, 3H, 7-C$H_3$), 5.74 (s, 1H, $H_3$), 7.56 (d, 2H, *J*=8.1 Hz, $H_{2'}$, $H_{6'}$), 7.86 (d, 2H, *J*=8.3 Hz, $H_{3'}$, $H_{5'}$), 10.13 (s, 1H, 1-OH), 14.98 (s, 1H, 6-OH); $^{13}C$ NMR (150.9 MHz, DMSO-$d_6$) $\delta$ (ppm) 20.9 (7-$CH_3$), 25.0 (4-*C*$H_3$), 100.1 ($C_5$), 110.9 ($C_3$), 123.1, 124.9 (d, $J_{C\text{-}F}$=272.0 Hz, $C_{4'}$), 126.2 ($C_{2'}$, $C_{6'}$), 126.5, 126.6 (d, $J_{C\text{-}F}$=4.1 Hz, $C_{3'}$, $C_{5'}$), 140.6 ($C_{1'}$), 148.3 ($C_4$), 158.8 ($C_2$), 164.4 ($C_6$), 168.2 ($C_7$). Anal. Calcd for $C_{15}H_{13}F_3N_2O_3$: C, 55.22; H, 4.02; N, 8.59; Found: C, 55.20; H, 4.00; N, 8.65.

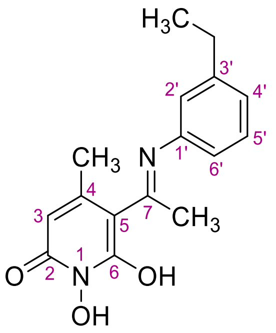


**5-(1-((3-ethylphenyl)imino)ethyl)-1,6-dihydroxy-4-methylpyridin-2(1*H*)-one (ZEV-V3)**

200 mg (1.09 mmol, 1.0 eq) of **ZEV2** was dissolved in 4 mL of absolute EtOH at 75 °C. Then, 3-ethylaniline (145 mg, 149 µL, 1.2 mmol, 1.1 eq), activated 4Å molecular sieves and a drop of sulfuric acid were added successively. After being refluxed at 60 °C for 1.5 h under Argon, a yellow solid was formed, and the stirring was continued at the same temperature for 3 h. The precipitate was filtered off *in vacuo* and washed with EtOH subsequently to afford the *N*-hydroxypyridinedione **ZEV-V3** as a yellow crystalline solid (205 mg, 66%). Mp 202-204 °C (EtOH), $R_{f\ (alox)}$ = 0.08 (MeOH), $R_{f\ (RP\text{-}TLC)}$ = 0.02 ($H_2O$/ACN 7:3). $^1$H NMR (600.11 MHz, DMSO-$d_6$) $\delta$ (ppm) 1.21 (t, 3H, $J$=7.6 Hz, $CH_2CH_3$), 2.37 (s, 3H, 4-$CH_3$), 2.48 (s, 3H, 7-$CH_3$), 2.66 (q, 2H, $J$=7.6 Hz, $CH_2CH_3$), 5.69 (s, 1H, $H_3$), 7.14 (dd, 1H, $J_1$=8.1 Hz, $J_2$=2.2 Hz, $H_{6'}$), 7.17 (s, 1H, $H_{2'}$), 7.24 (d, 1H, $J$=7.7 Hz, $H_{4'}$), 7.41 (t, 1H, $J$=7.7 Hz, $H_{5'}$), 10.06 (s, 1H, 1-OH), 14.86 (s, 1H, 6-OH); $^{13}$C NMR (150.9 MHz, DMSO-$d_6$) $\delta$ (ppm) 15.3 ($CH_2CH_3$), 20.7 (7-$CH_3$), 25.0 (4-$CH_3$), 27.8 ($CH_2CH_3$), 99.2 ($C_5$), 110.1 ($C_3$), 122.8 ($C_{6'}$), 124.8 ($C_{2'}$), 126.9 ($C_{4'}$), 129.4 ($C_{5'}$), 136.7 ($C_{3'}$), 145.5 ($C_{1'}$), 148.4 ($C_4$), 158.8 ($C_2$), 164.3 ($C_6$), 168.6 ($C_7$). Anal. Calcd for $C_{16}H_{18}N_2O_3$: C, 67.12; H, 6.34; N, 9.78; Found: C, 67.05; H, 6.39; N, 9.85.

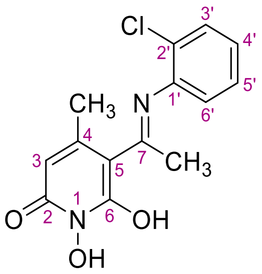


**5-(1-((2-chlorophenyl)imino)ethyl)-1,6-dihydroxy-4-methylpyridin-2(1*H*)-one (ZEV-V5)**

To a solution of **ZEV2** (200 mg, 1.09 mmol, 1.0 eq) in absolute EtOH (4 mL) at 75 °C were added successively 2-chroloaniline (153 mg, 126 µL, 1.2 mmol, 1.1 eq), activated 4Å molecular sieves and a drop of sulfuric acid. The mixture was heated under reflux at 60 °C for 24 h under Argon. The solvent was concentrated to half of its volume and the solid formed was filtered off *in vacuo*. The precipitate was washed with small portions of EtOH to afford the title compound as a yellow crystalline solid (80 mg, 25%). Mp 184-186 °C (AcOEt/*n*-pentane), $R_{f\ (alox)}$ = 0.01 (MeOH), $R_{f\ (RP\text{-}TLC)}$ = 0.05 ($H_2O$/ACN 7:3). $^1$H NMR (600.11 MHz, DMSO-$d_6$) $\delta$ (ppm) 2.39 (s, 3H, 4-$CH_3$), 2.42 (s, 3H, 7-$CH_3$), 5.73 (s, 1H, $H_3$), 7.43 (td, 1H, $J_1$=7.7 Hz, $J_2$=1.7 Hz, $H_{5'}$), 7.49 (td, 1H, $J_1$=7.6 Hz, $J_2$=1.5 Hz, $H_{4'}$), 7.54 (dd, 1H, $J_1$=7.9 Hz, $J_2$=1.7 Hz, $H_{3'}$), 7.68 (dd, 1H, $J_1$=8.0 Hz, $J_2$=1.5 Hz, $H_{6'}$), 10.12 (s, 1H, 1-OH), 14.94 (s, 1H, 6-OH); $^{13}$C NMR (150.9 MHz, DMSO-$d_6$) $\delta$ (ppm) 20.6 (7-$CH_3$), 25.2 (4-$CH_3$), 99.5 ($C_5$), 110.8 ($C_3$), 28.1 ($C_{4'}$), 128.6 ($C_{3'}$), 129.1 ($C_{2'}$), 129.2 ($C_{5'}$), 130.1 ($C_{6'}$), 134.4 ($C_{1'}$), 148.4 ($C_4$), 158.8 ($C_2$), 164.7 ($C_6$), 168.9 ($C_7$). Anal. Calcd for $C_{14}H_{13}ClN_2O_3$: C, 57.45; H, 4.48; N, 9.57; Found: C, 57.53; H, 4.50; N, 9.47.

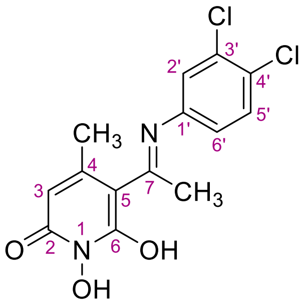


**5-(1-((3,4-dichlorophenyl)imino)ethyl)-1,6-dihydroxy-4-methylpyridin-2(1*H*)-one (ZEV-V7)**

3,4-dichlroaniline (194 mg, 1.2 mmol, 1.1 eq) was added to a solution of 200 mg (1.09 mmol, 1.0 eq) of **ZEV2** in 4 mL absolute EtOH at 75 °C. Then, activated 4Å molecular sieves and a drop of sulfuric acid were added, and a yellow solid was precipitated. The temperature was adjusted to 60 °C and the mixture was refluxed for 4 h under Argon. The precipitate was filtered off under vacuum and washed with EtOH to give the title compound **ZEV-V7** as a yellow crystalline solid (250 mg, 70%). Mp 217-219 °C (EtOH), $R_{f\ (alox)}$ = 0.01 (MeOH), $R_{f\ (RP\text{-}TLC)}$ = 0.01 ($H_2O$/ACN 7:3). $^1$H NMR (600.11 MHz, DMSO-$d_6$) $\delta$ (ppm) 2.37 (s, 3H, 4-C$H_3$), 2.48 (s, 3H, 7-C$H_3$), 5.73 (s, 1H, $H_3$), 7.35 (dd, 1H, $J_1$=8.6 Hz, $J_2$=2.5 Hz, $H_{6'}$), 7.71 (d, 1H, $J$=2.5 Hz, $H_{2'}$), 7.75 (d, 1H, $J$=8.5 Hz, $H_{5'}$), 10.10 (s, 1H, 1-OH), 14.79 (s, 1H, 6-OH); $^{13}$C NMR (150.9 MHz, DMSO-$d_6$) $\delta$ (ppm) 20.8 (7-$C$H$_3$), 25.0 (4-$C$H$_3$), 99.9 ($C_5$), 110.8 ($C_3$), 126.1 ($C_{6'}$), 127.7 ($C_{2'}$), 129.8 ($C_{4'}$), 131.2 ($C_{5'}$), 131.7 ($C_{3'}$), 137.0 ($C_{1'}$), 148.3 ($C_4$), 158.8 ($C_2$), 164.4 ($C_6$), 168.4 ($C_7$). Anal. Calcd for $C_{14}H_{12}Cl_2N_2O_3$: C, 51.40; H, 3.70; N, 8.56; Found: C, 51.44; H, 3.65; N, 8.60.

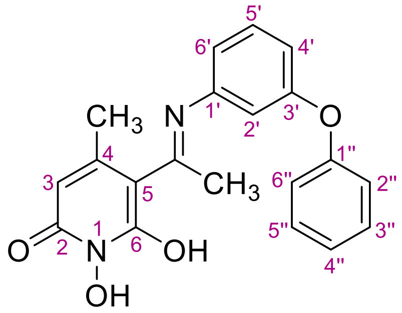


**5-(1-((3-benzylphenyl)imino)ethyl)-1,6-dihydroxy-4-methylpyridin-2(1*H*)-one (ZEV-E2)**

**ZEV2** (200 mg, 1.09 mmol, 1.0 eq) was diluted in absolute EtOH (4 mL) at 75 °C. Subsequently, 3-phenoxy aniline (223 mg, 1.2 mmol, 1.1 eq) and 4Å molecular sieves were added, followed by a drop of sulfuric acid. Reaction was refluxed at 60 °C, under Argon atmosphere. Within the first hour a yellow solid was formed and stirring was continued under the same reaction conditions for a total of 5 h. The precipitate was filtered off under vacuum and washed with EtOH to afford the desired compound **ZEV-E2** as a yellow crystalline solid (260 mg, 62%). Mp 157-159 °C (EtOH), $R_{f\ (alox)}$ = 0.06 (MeOH), $R_{f\ (RP\text{-}TLC)}$ = 0.06 ($H_2O$/ACN 7:3). $^1$H NMR (600.11 MHz, DMSO-$d_6$) $\delta$ (ppm) 2.35 (s, 3H, 4-C$H_3$), 2.48 (s, 3H, 7-C$H_3$), 5.70 (s, 1H, $H_3$), 7.00 (dd, 2H, $J_1$=4.4 Hz, $J_2$=2.1 Hz, $H_{4'}$, $H_{6'}$), 7.09 (d, 3H, $J$= 8.0 Hz, $H_{2'}$, $H_{2''}$,$H_{6''}$), 7.19 (t, 1H, $J$=7.4 Hz, $H_{4''}$), 7.42 (td, 2H, $J_1$=7.0 Hz, $J_2$=1.9 Hz, $H_{3''}$,$H_{5''}$), 7.48-7.52 (m, 1H, $J$=7.4 Hz, $H_{5'}$), 10.09 (s, 1H, 1-OH), 14.86 (s, 1H, 6-OH); $^{13}$C NMR (150.9 MHz, DMSO-$d_6$) $\delta$ (ppm) 20.8 (7-$C$H$_3$), 25.1 (4-$C$H$_3$), 99.4 ($C_5$), 110.3 ($C_3$), 115.7 ($C_{6'}$), 117.1 ($C_{4'}$), 119.0 ($C_{2''}$, $C_{6''}$), 120.5 ($C_{2'}$), 124.0 ($C_{4''}$), 130.2 ($C_{3''}$, $C_{5''}$), 130.9 ($C_{5'}$), 138.2 ($C_{1'}$), 148.4 ($C_4$), 156.0 ($C_{1''}$), 157.5 ($C_{3'}$), 158.8 ($C_2$), 164.3 ($C_6$), 168.7 ($C_7$). Anal. Calcd for $C_{21}H_{20}N_2O_3$: C, 72.40; H, 5.79; N, 8.04; Found: C, 72.52; H, 5.82; N, 8.12.

### V Copies of NMR spectra

[1]H NMR of **ZEV1** (600.11 MHz, DMSO-$d_6$)

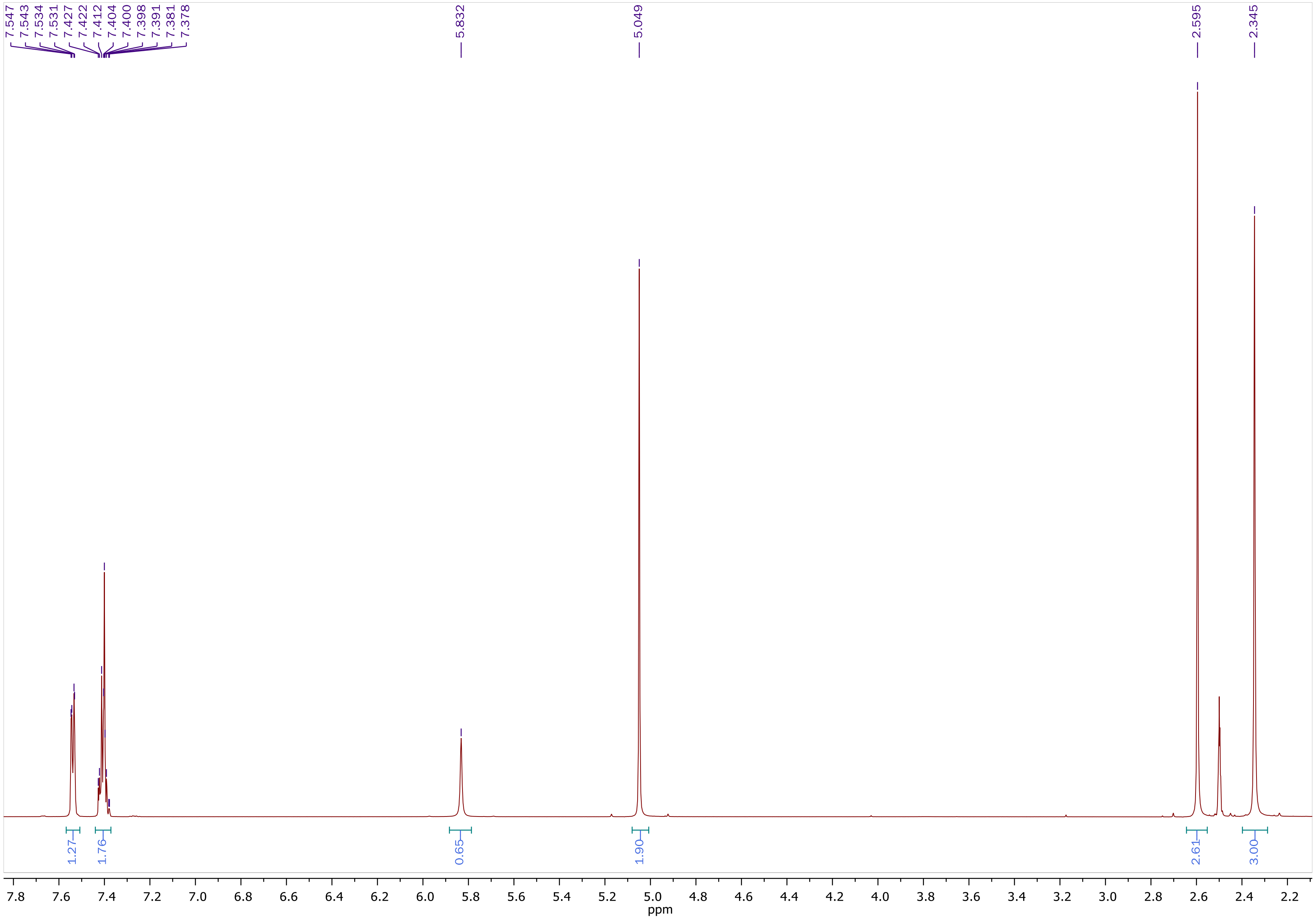

$^{13}C$ NMR of **ZEV1** (50.32 MHz, DMSO-$d_6$)

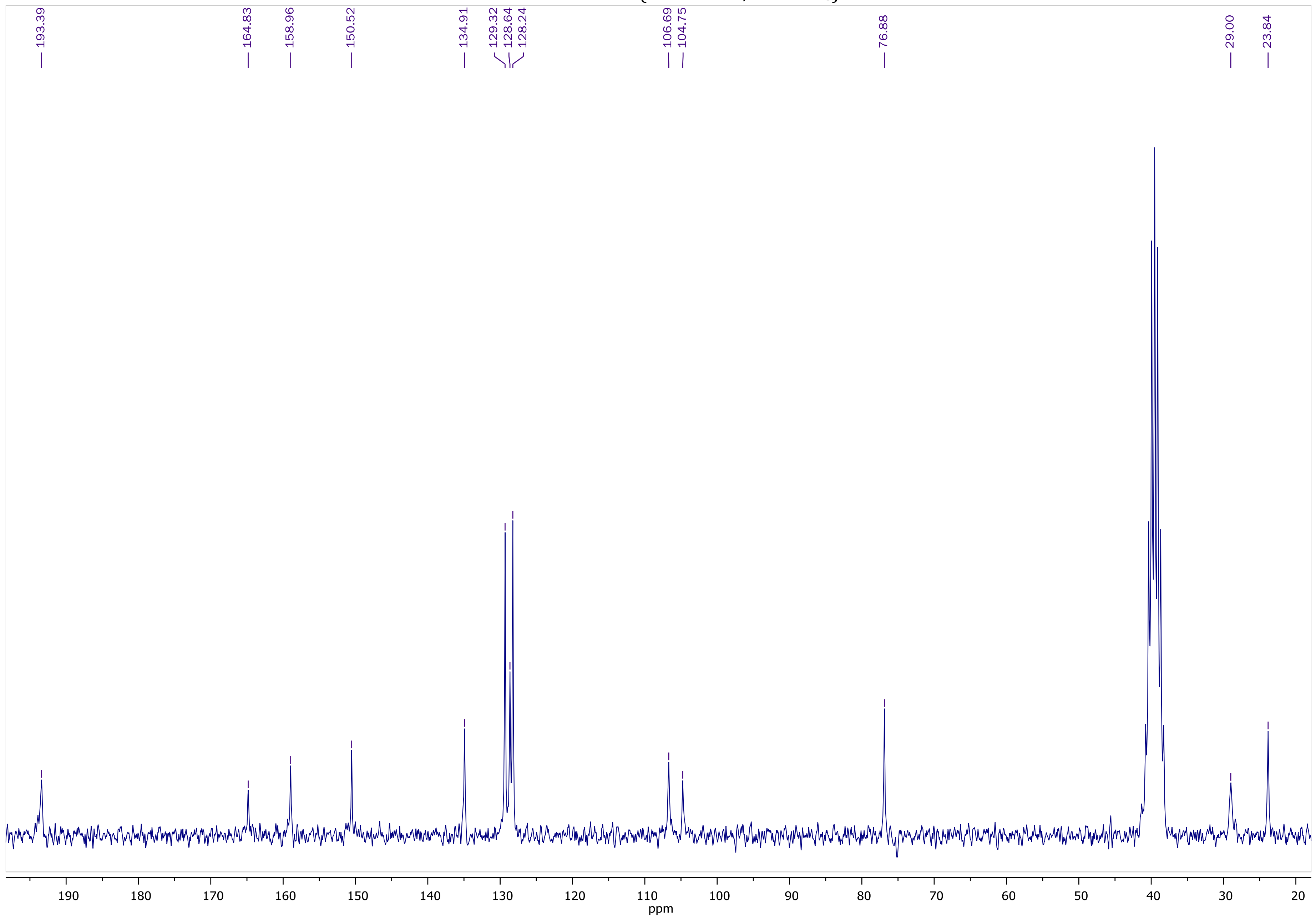

COSY NMR of **ZEV1** (400.13 MHz, DMSO-$d_6$)

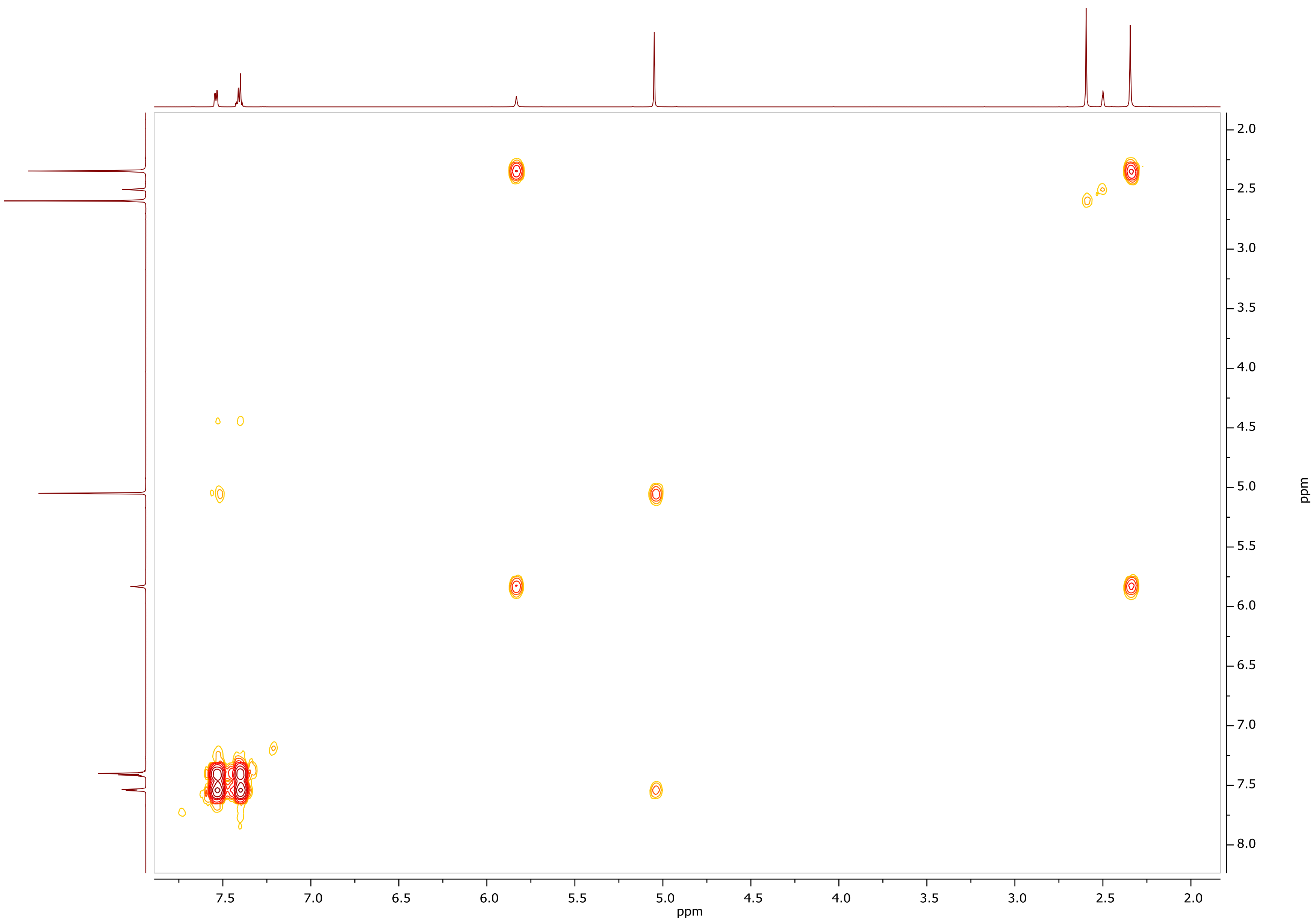

HSQC-DEPT NMR of **ZEV1** (400.13 MHz, DMSO-$d_6$)

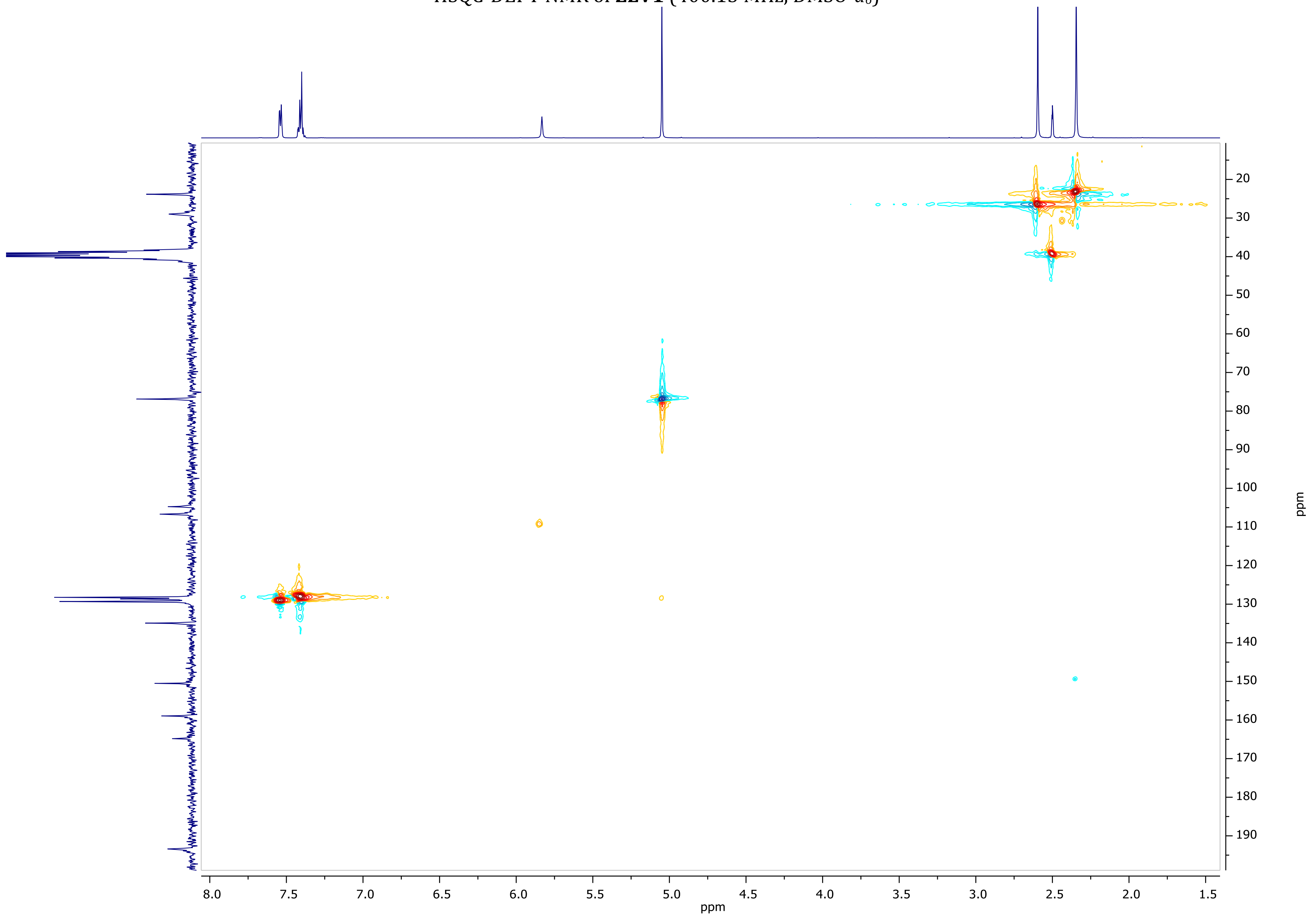

HMBC NMR of **ZEV1** (400.13 MHz, DMSO-$d_6$)

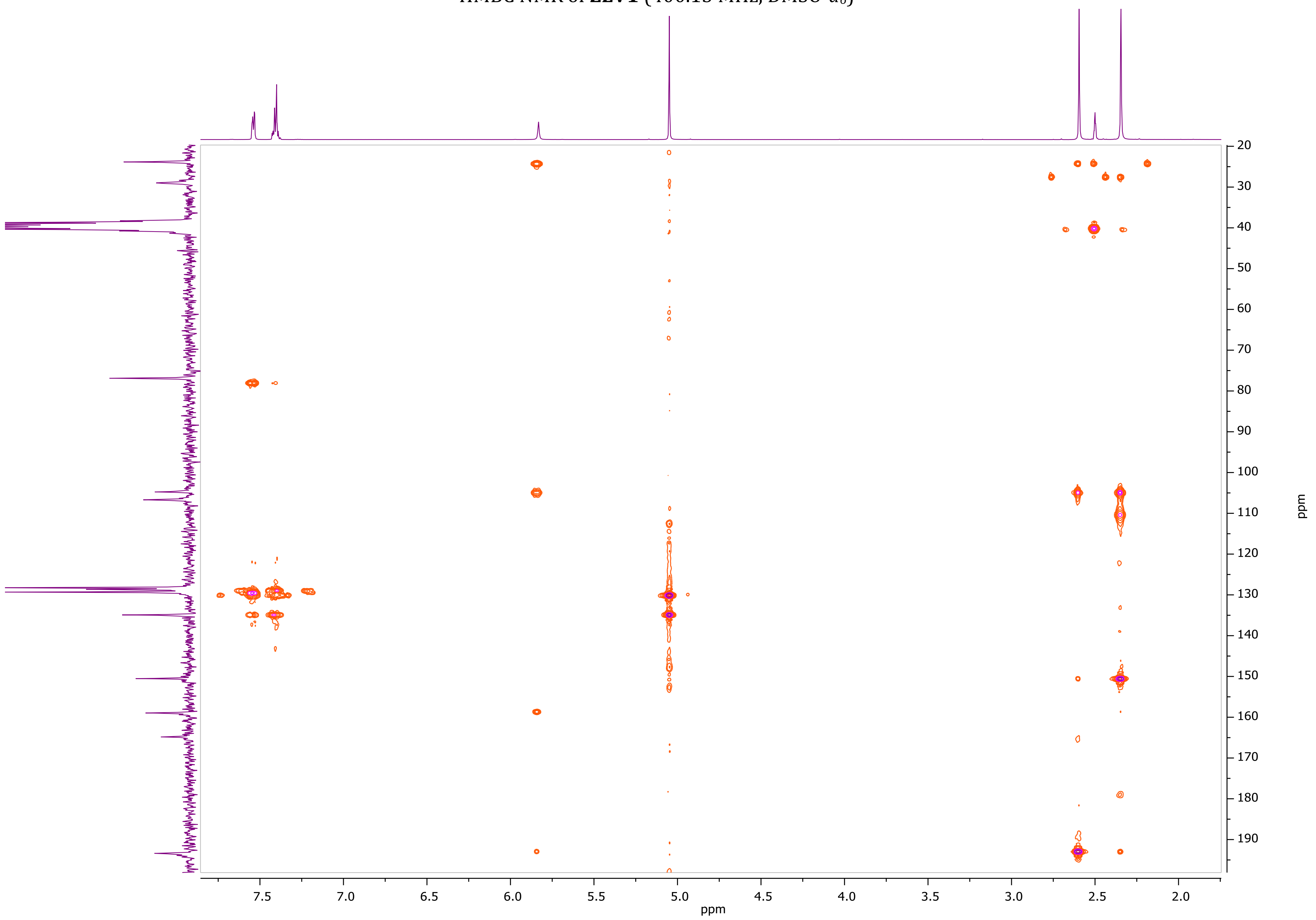

$^{1}H$ NMR of **ZEV2** (600.11 MHz, DMSO-$d_6$)

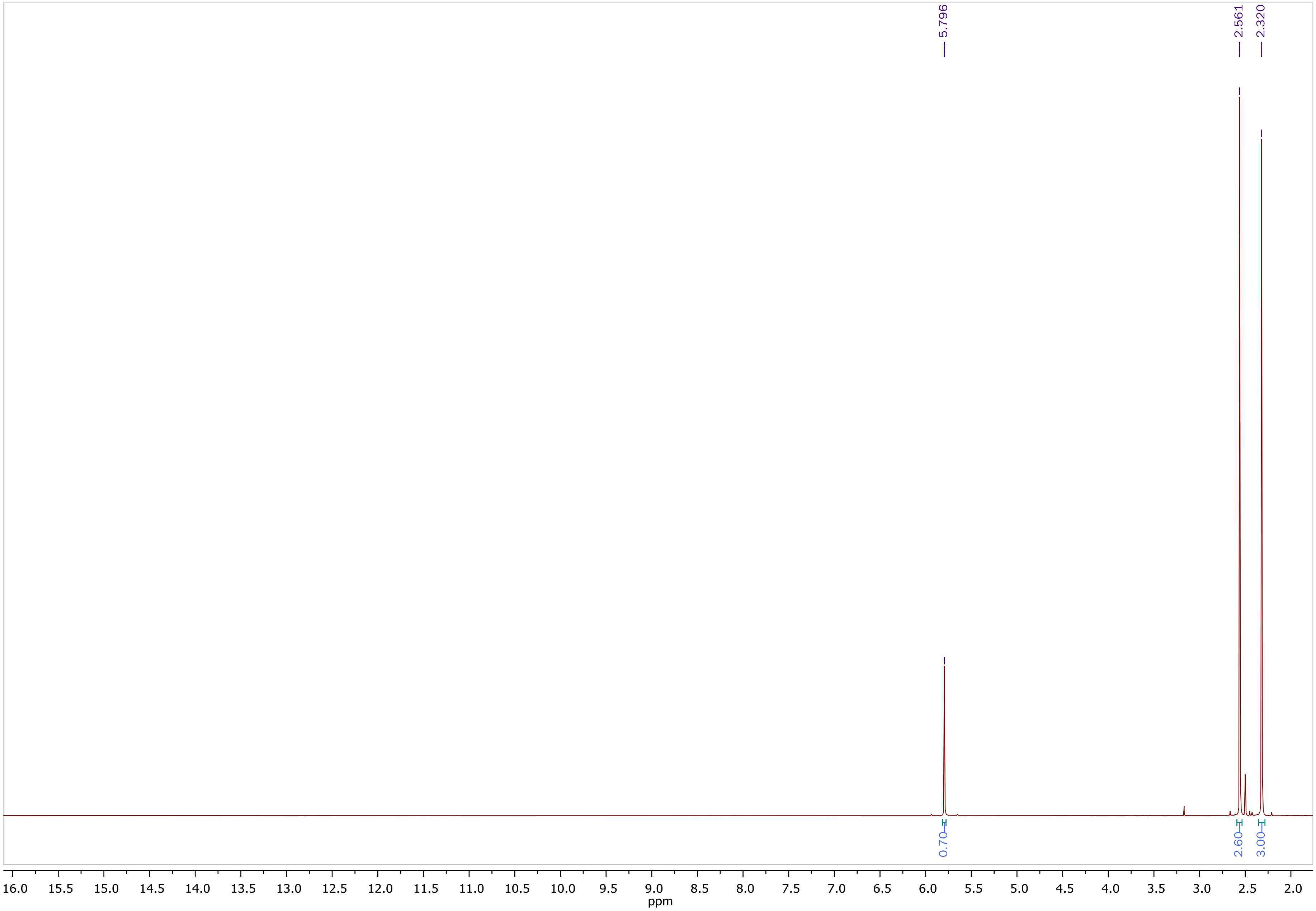

$^{13}$C NMR of **ZEV2** (100.61 MHz, DMSO-$d_6$)

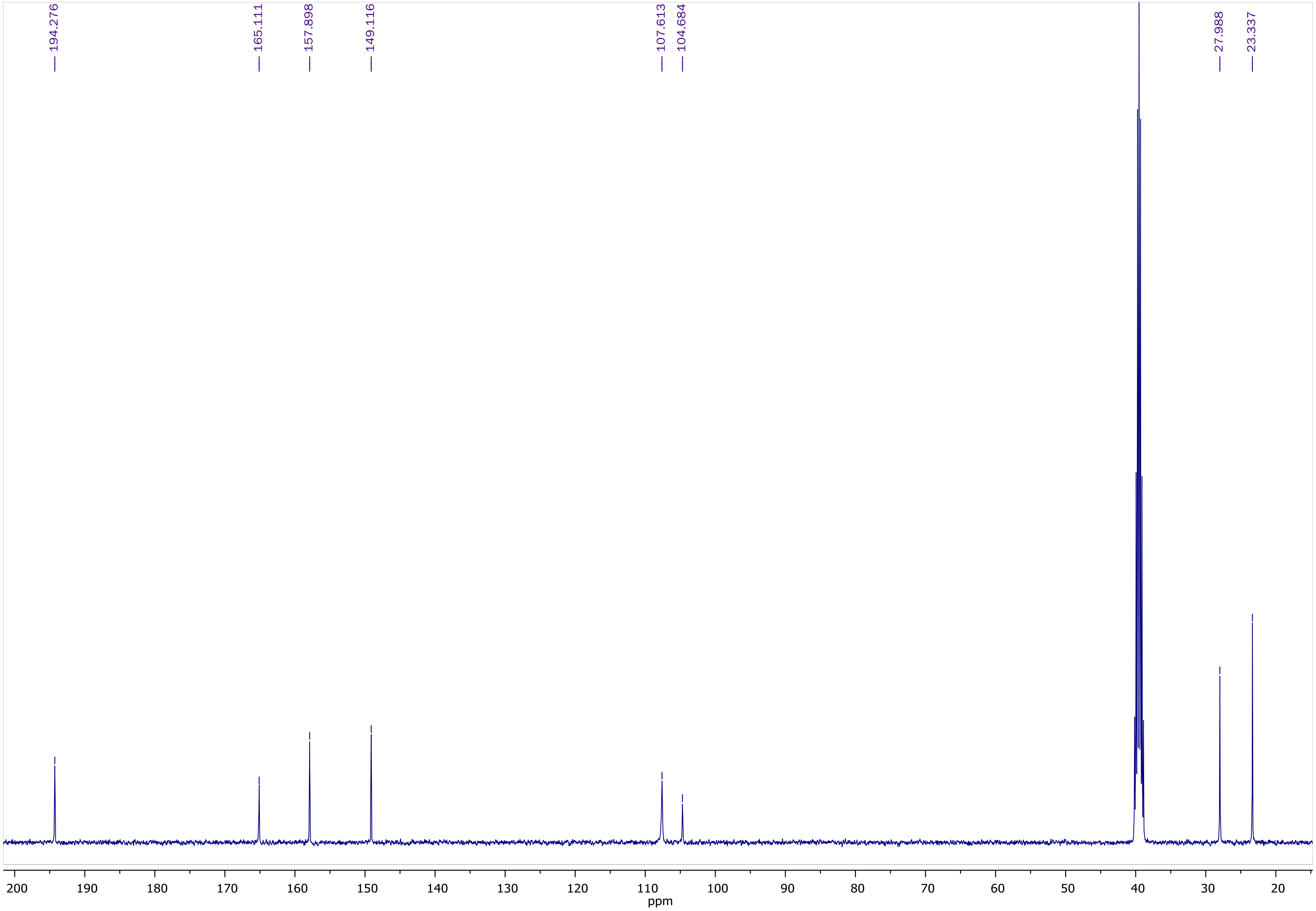

COSY NMR of **ZEV2** (400.13 MHz, DMSO-$d_6$)

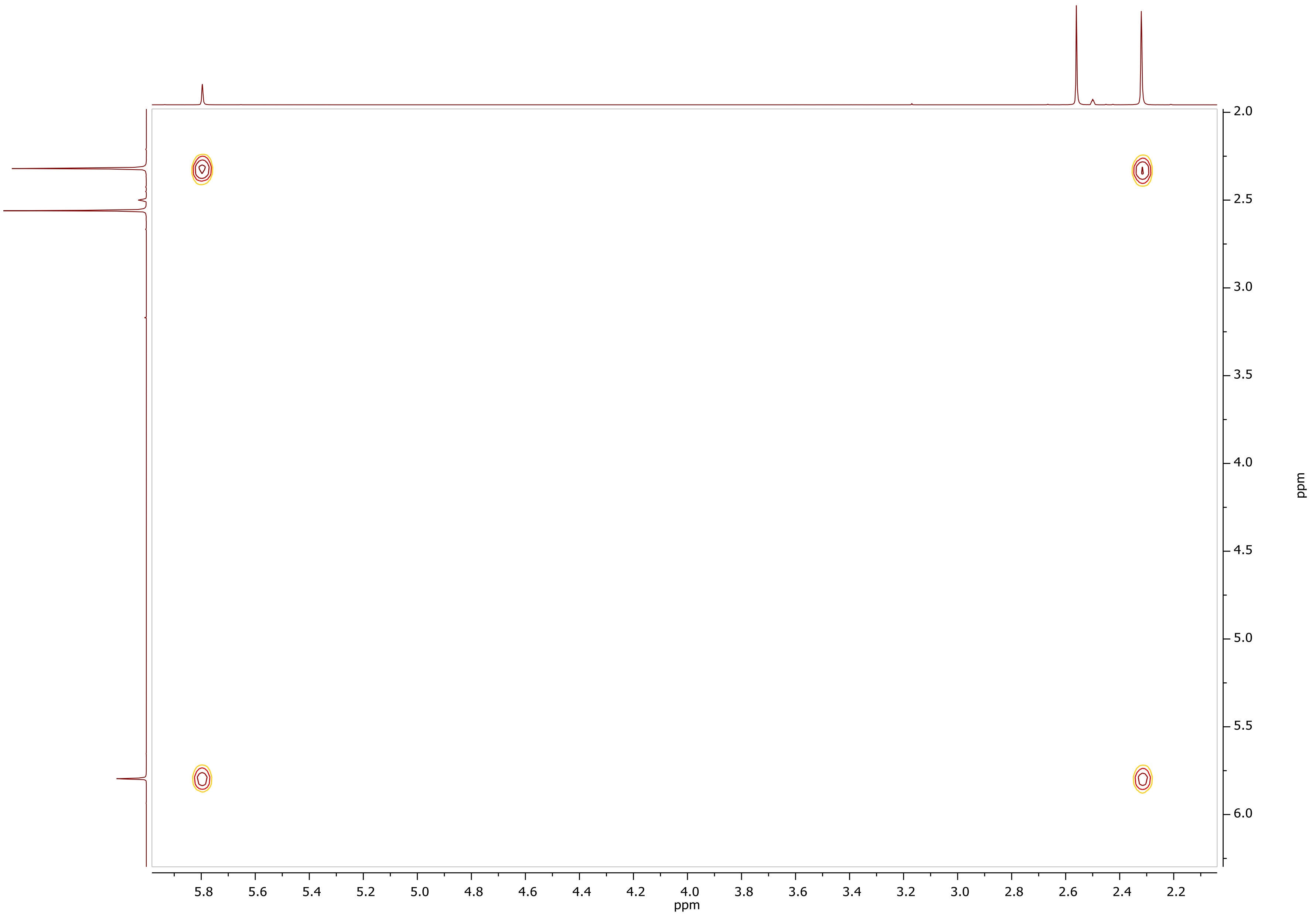

HSQC-DEPT NMR of **ZEV2** (400.13 MHz, DMSO-$d_6$)

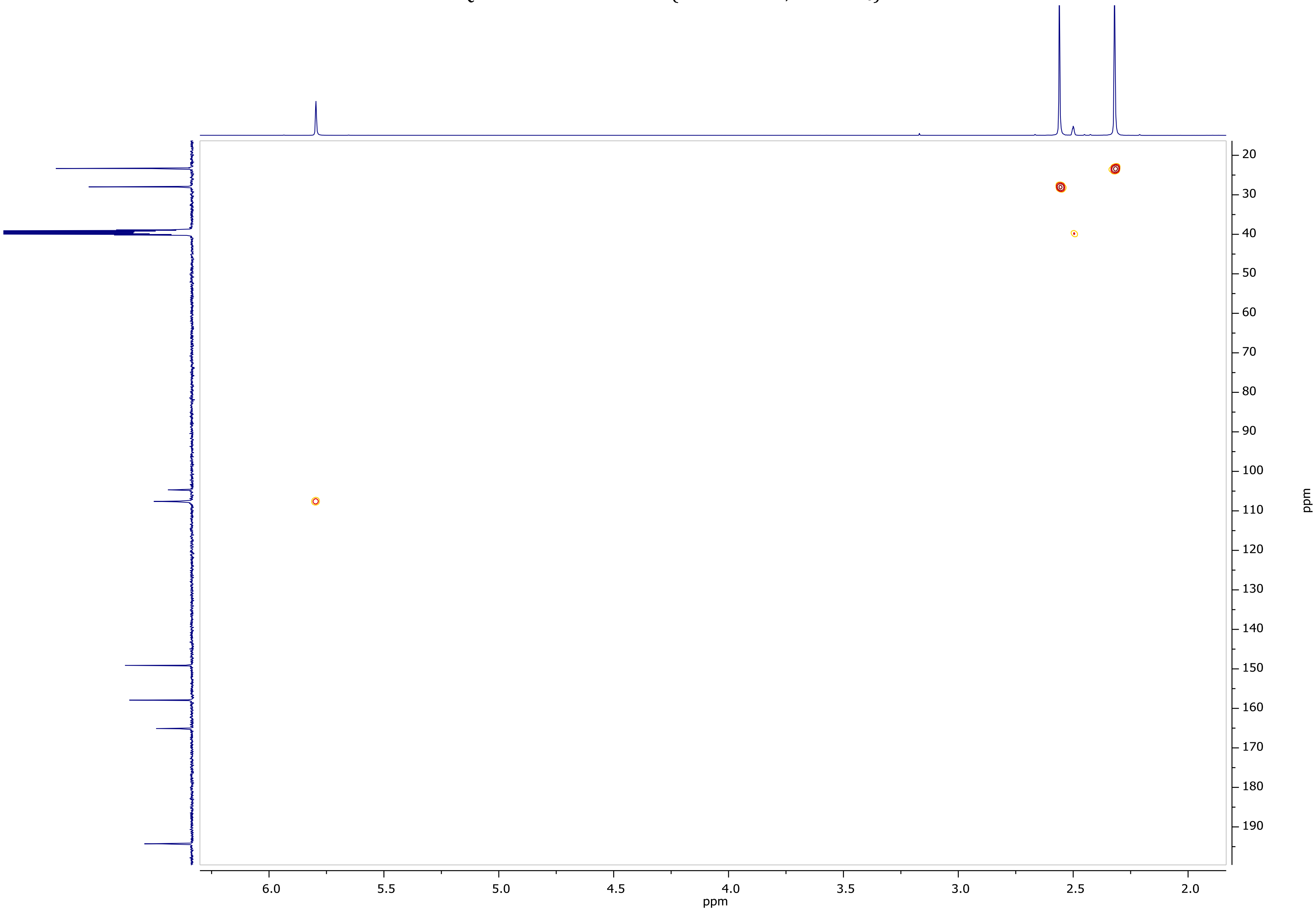

HMBC NMR of **ZEV2** (400.13 MHz, DMSO-$d_6$)

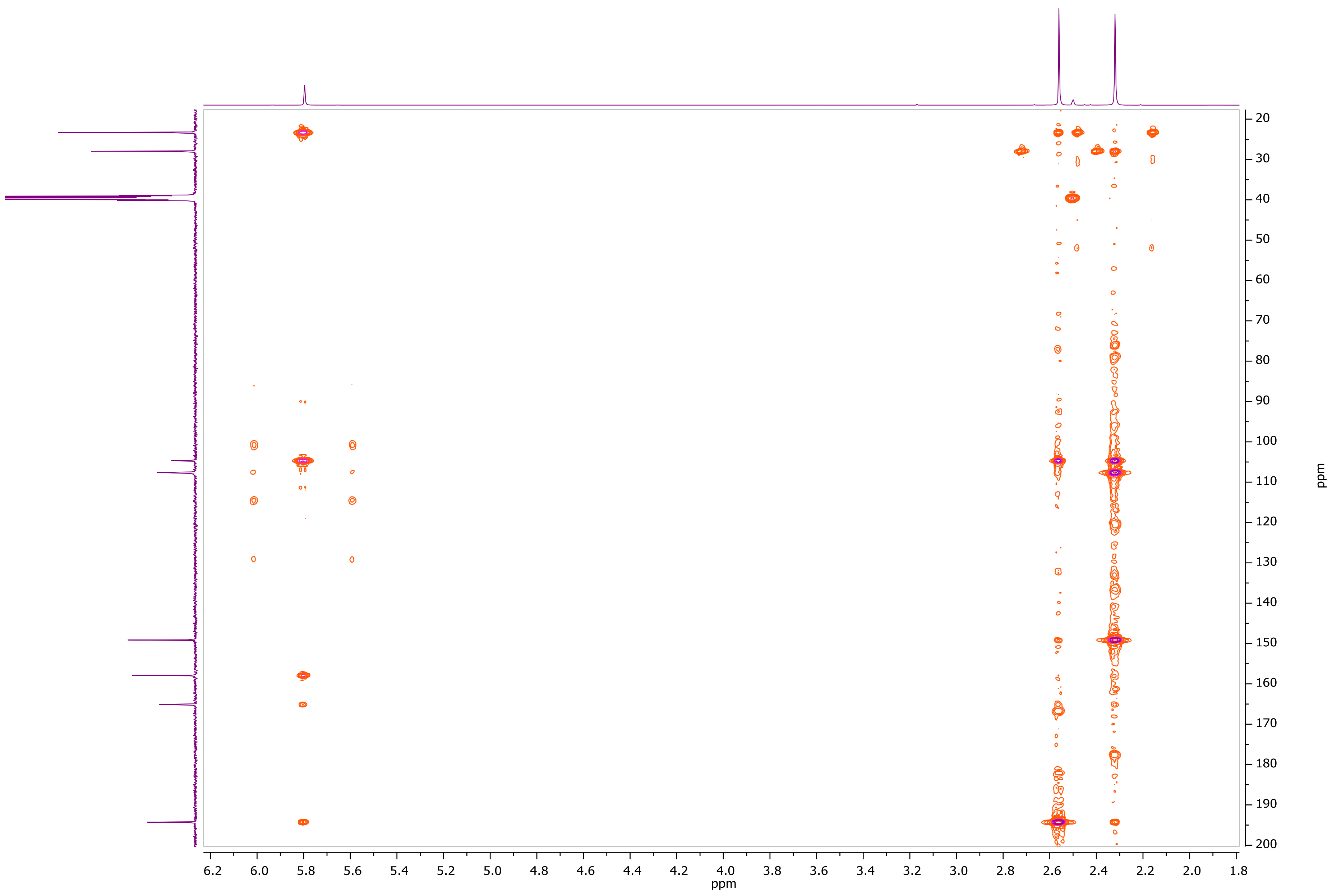

$^1$H NMR of **ZEV-V1** (600.11 MHz, DMSO-$d_6$)

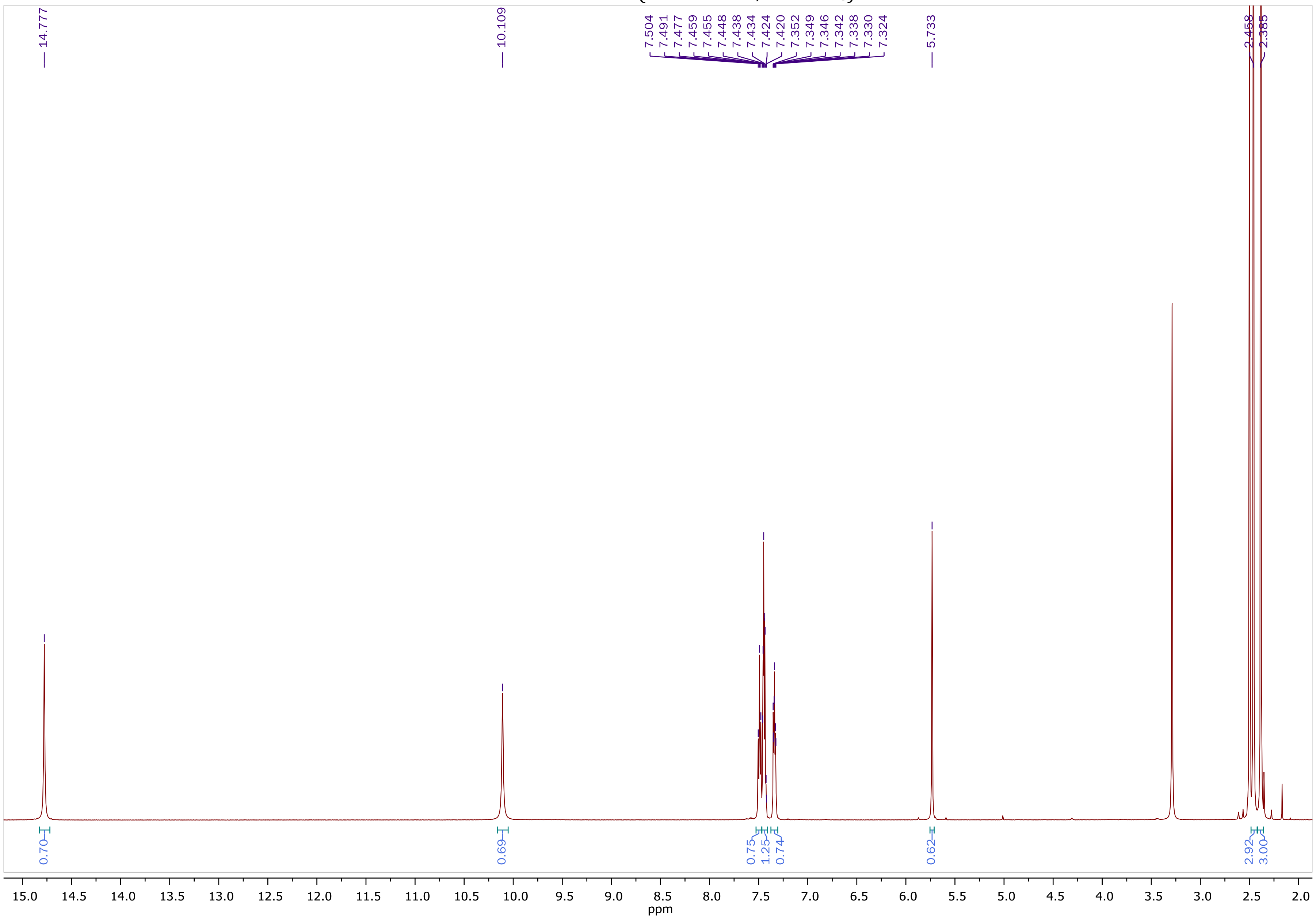

$^{13}$C NMR of **ZEV-V1** (150.9 MHz, DMSO-$d_6$)

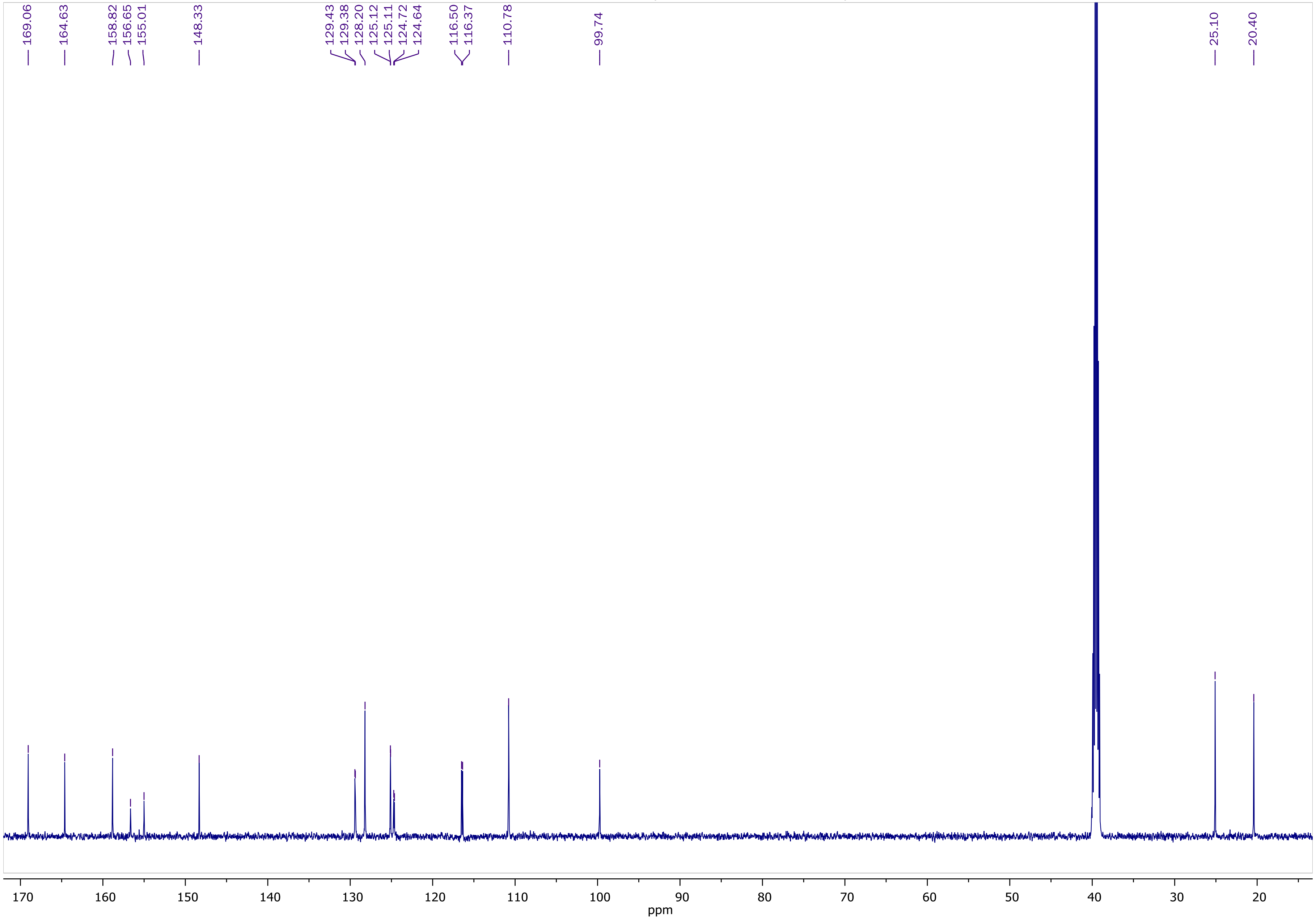

COSY NMR of **ZEV-V1** (600.11 MHz, DMSO-$d_6$)

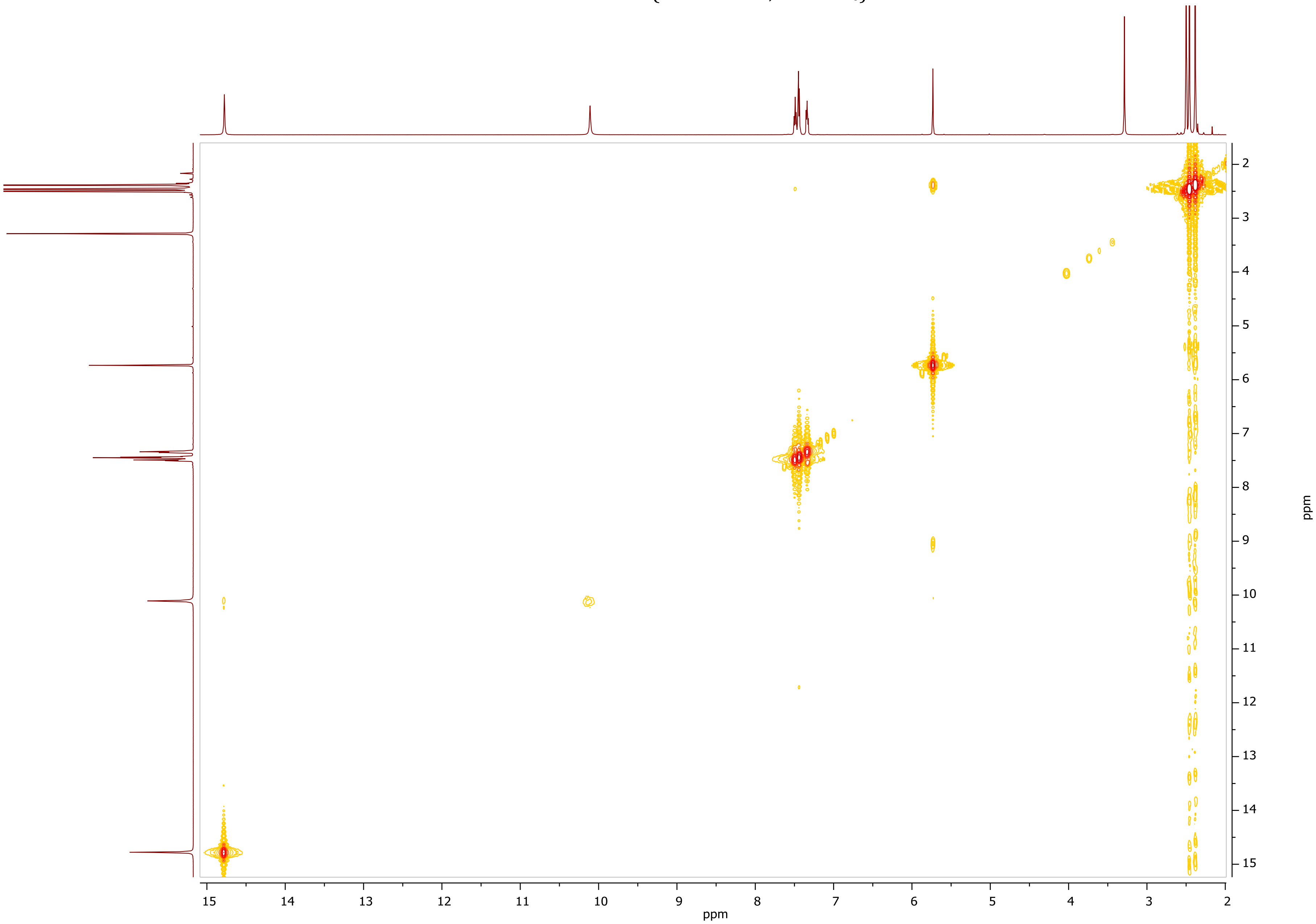

HSQC-DEPT NMR of **ZEV-V1** (600.11 MHz, DMSO-$d_6$)

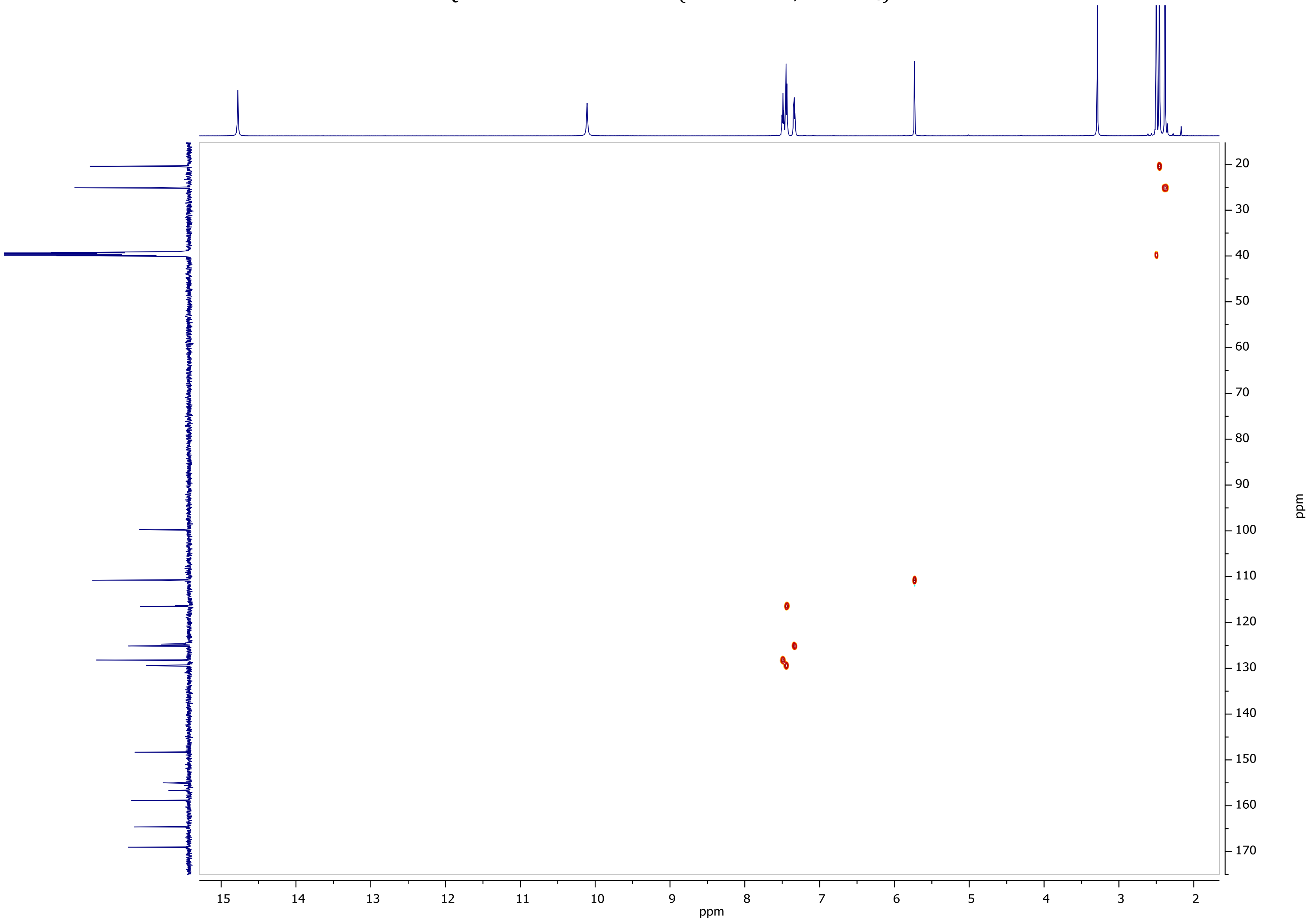

HMBC NMR of **ZEV-V1** (600.11 MHz, DMSO-$d_6$)

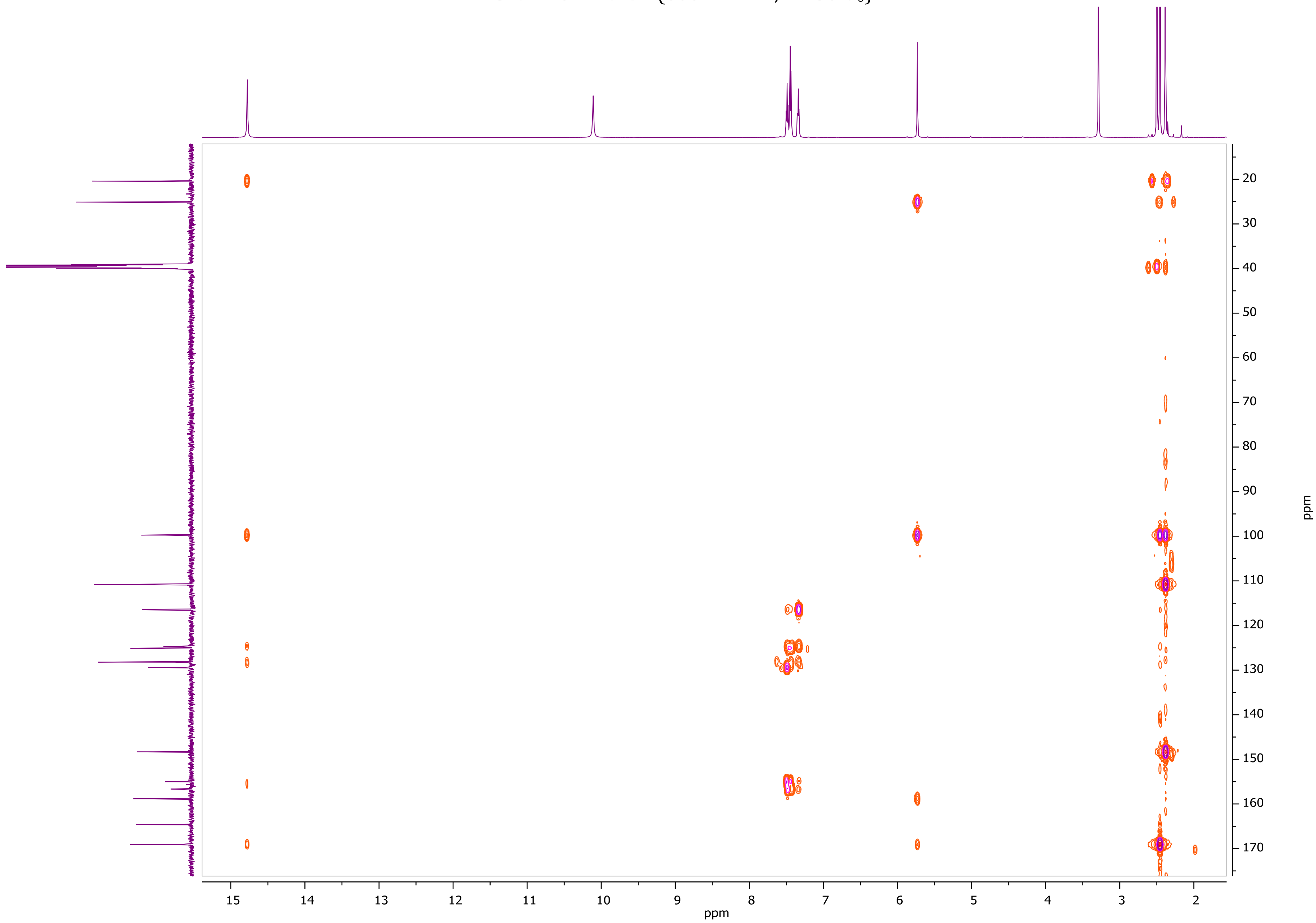

[1]H NMR of **ZEV-V2** (600.11 MHz, DMSO-$d_6$)

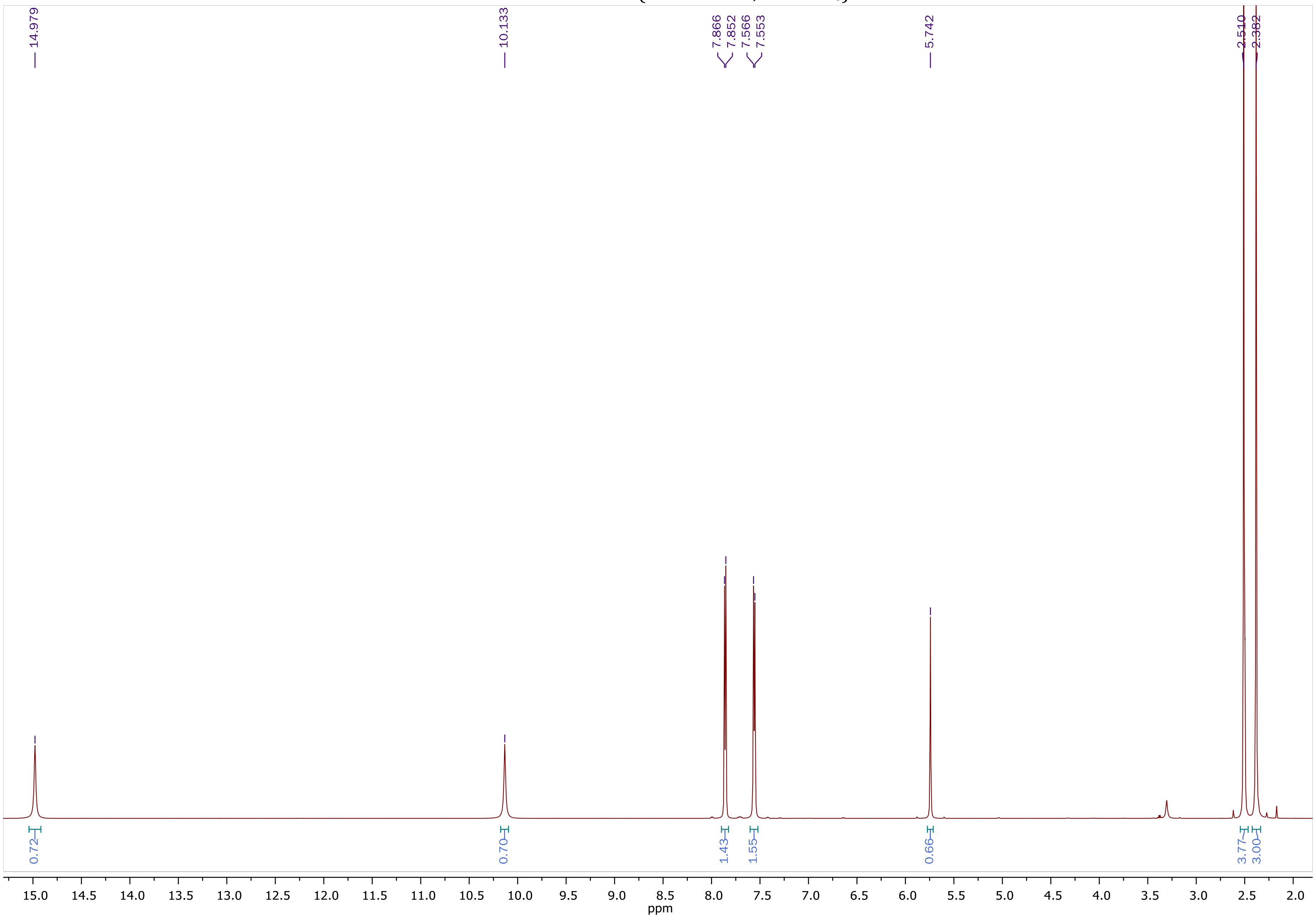

$^{13}C$ NMR of **ZEV-V2** (150.9 MHz, DMSO-$d_6$)

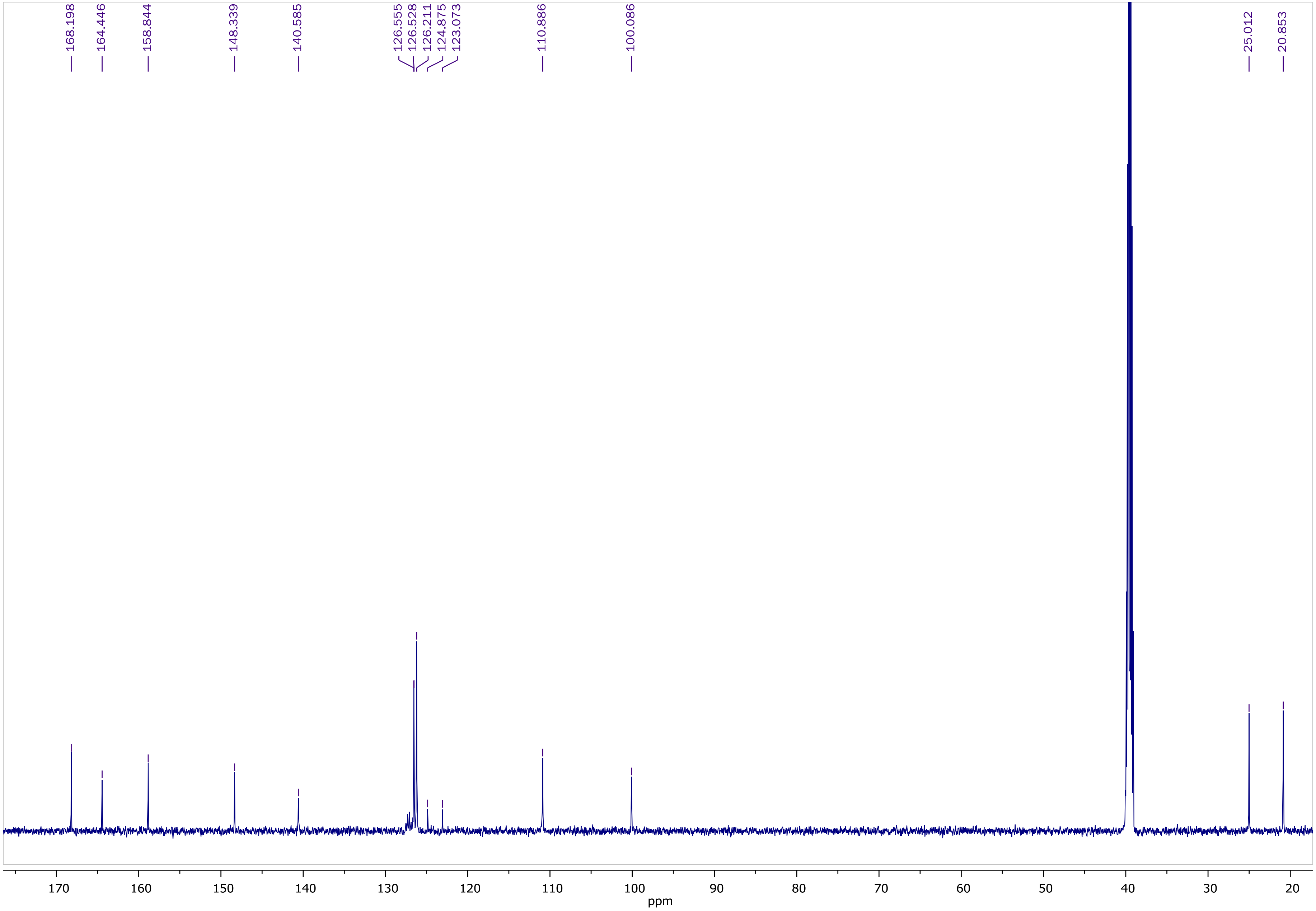

COSY NMR of **ZEV-V2** (600.11 MHz, DMSO-$d_6$)

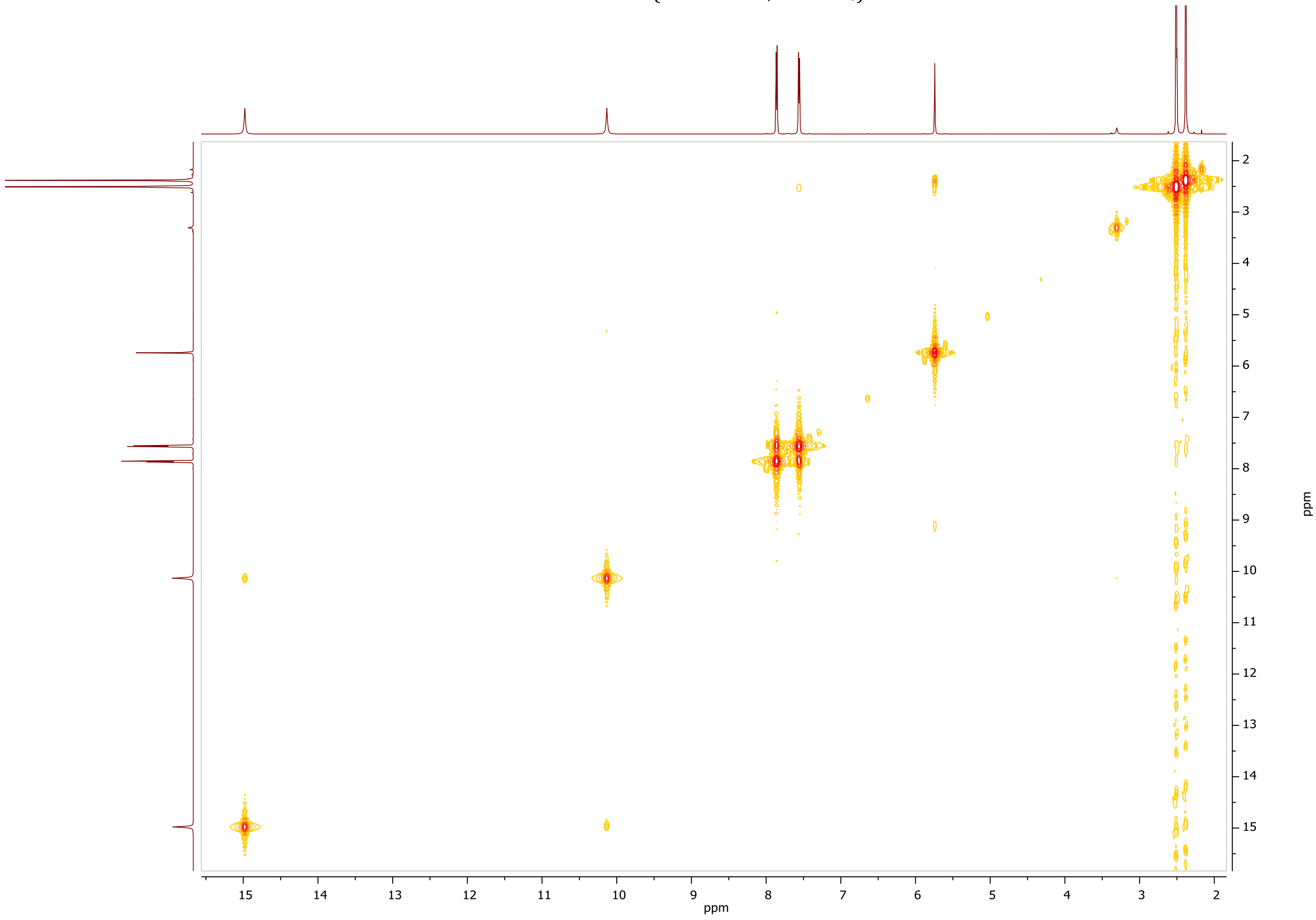

HSQC-DEPT NMR of **ZEV-V2** (600.11 MHz, DMSO-$d_6$)

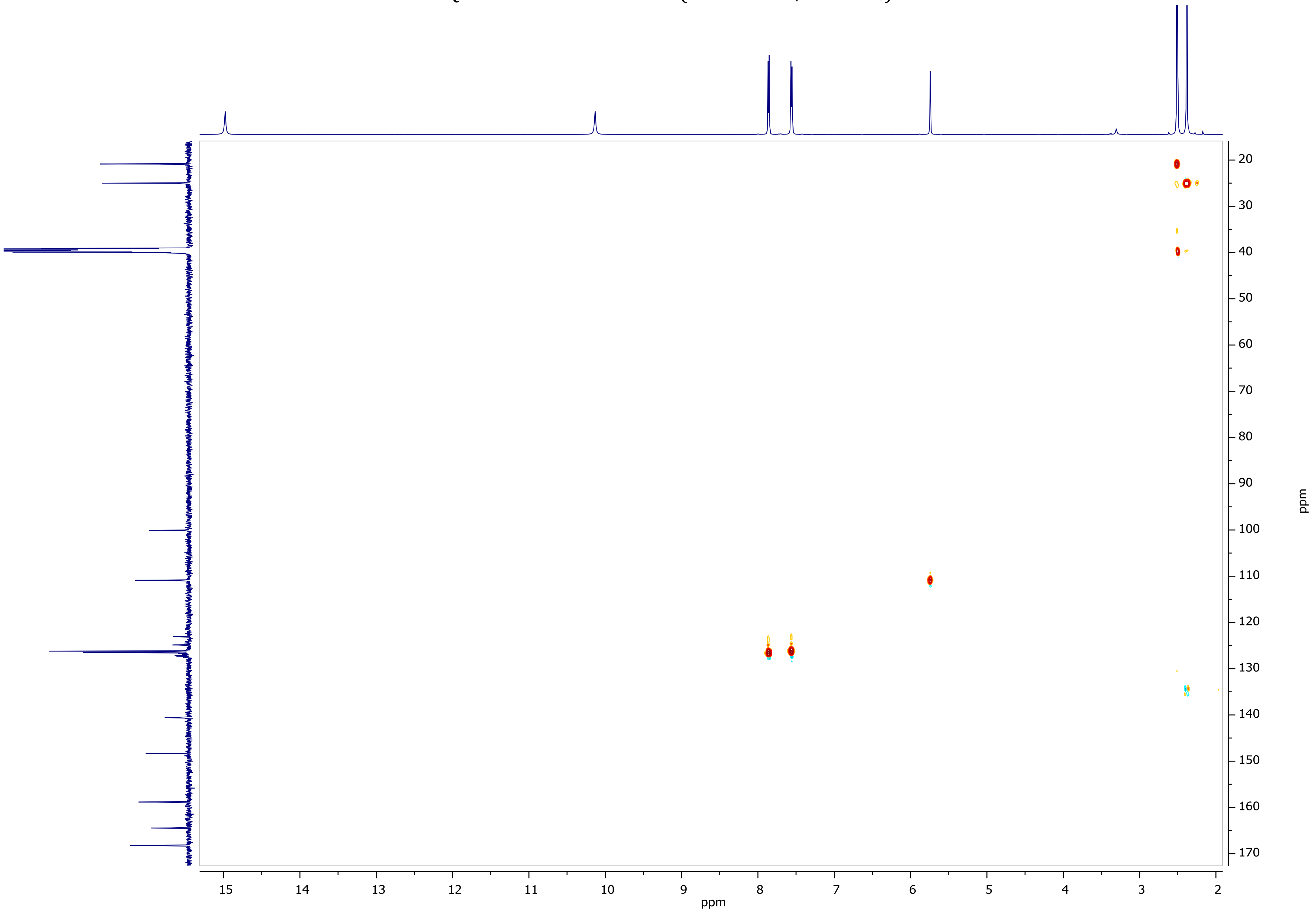

HMBC NMR of **ZEV-V2** (600.11 MHz, DMSO-$d_6$)

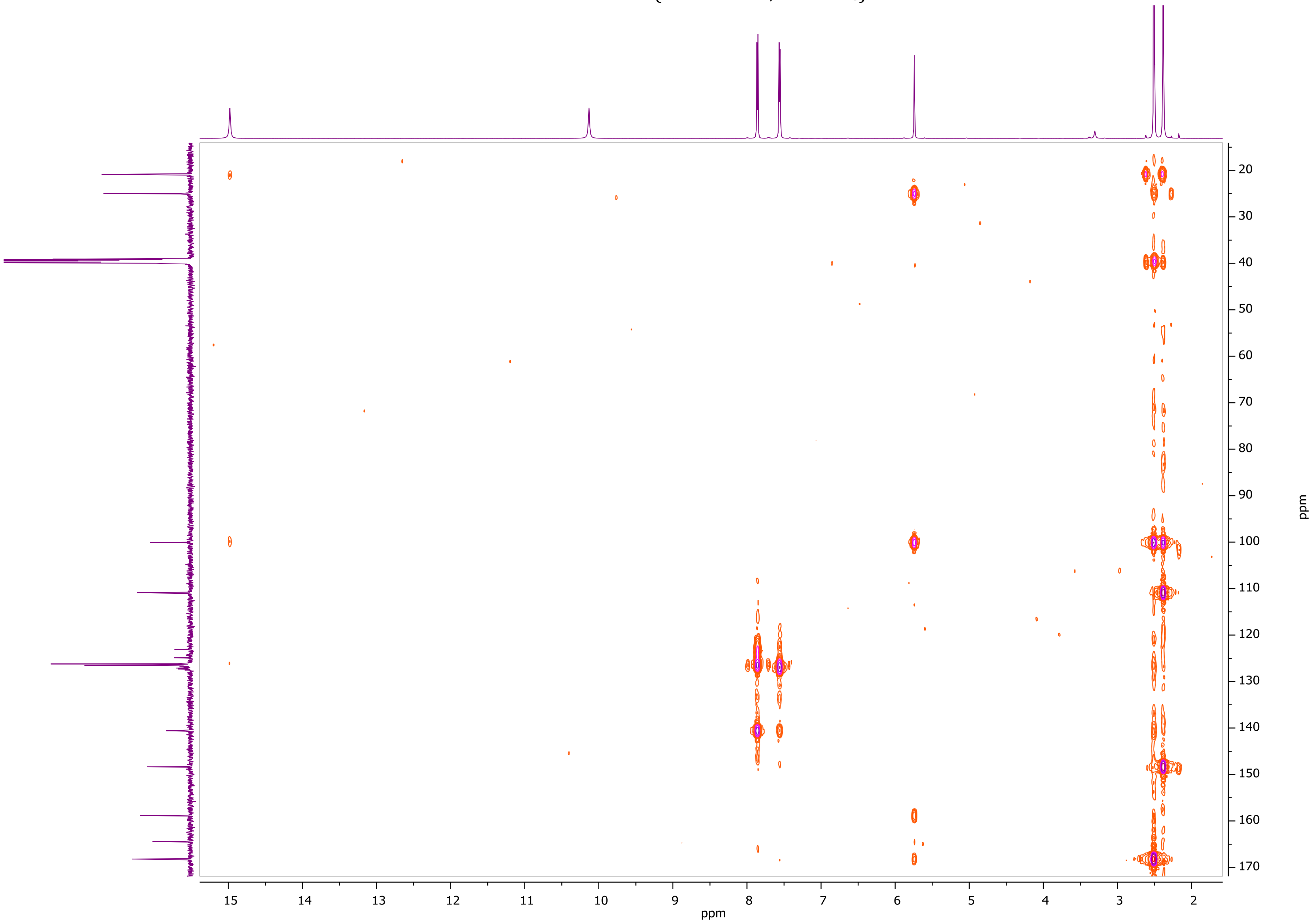

[1]H NMR of **ZEV-V3** (600.11 MHz, DMSO-$d_6$)

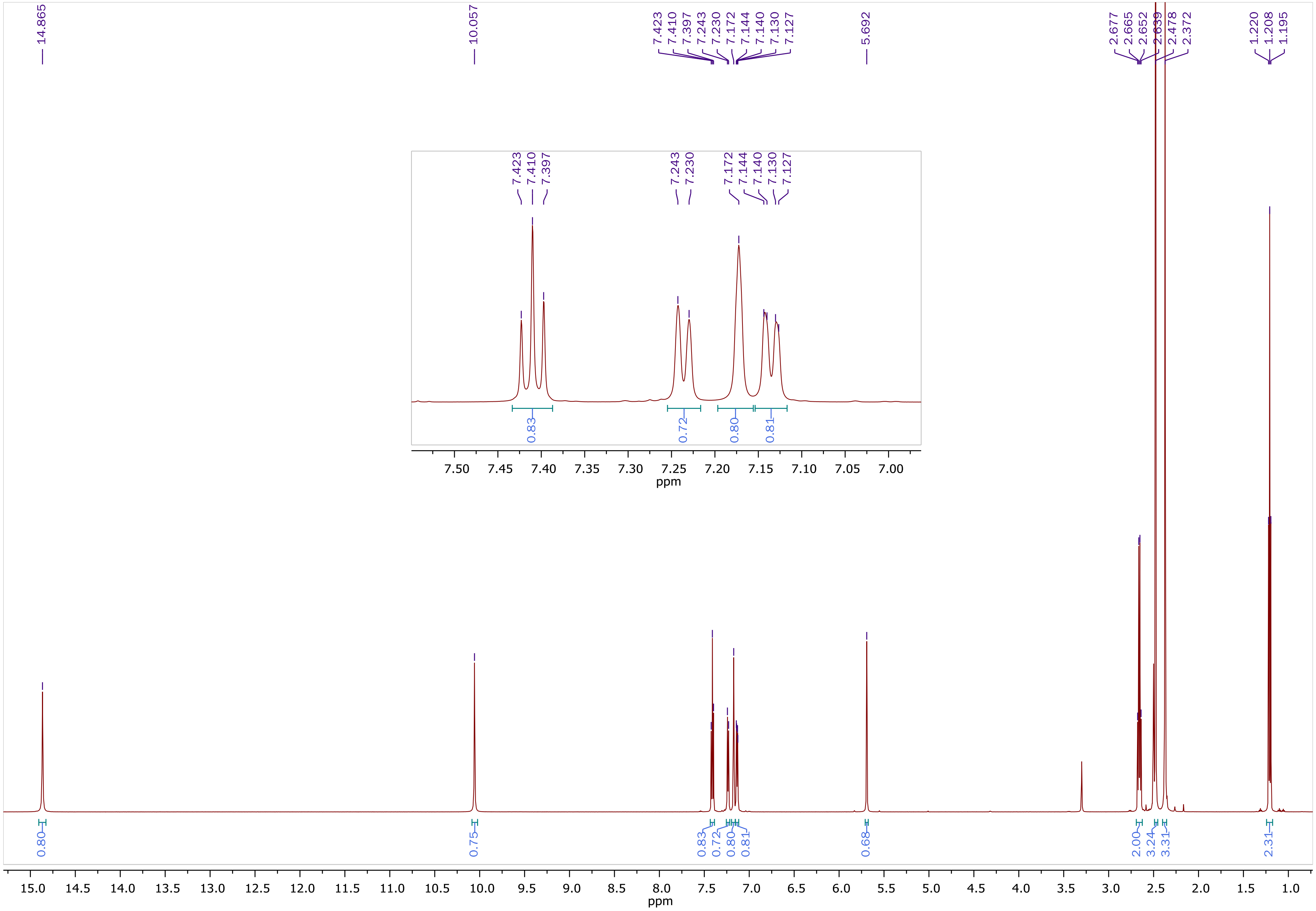

$^{13}$C NMR of **ZEV-V3** (150.9 MHz, DMSO-$d_6$)

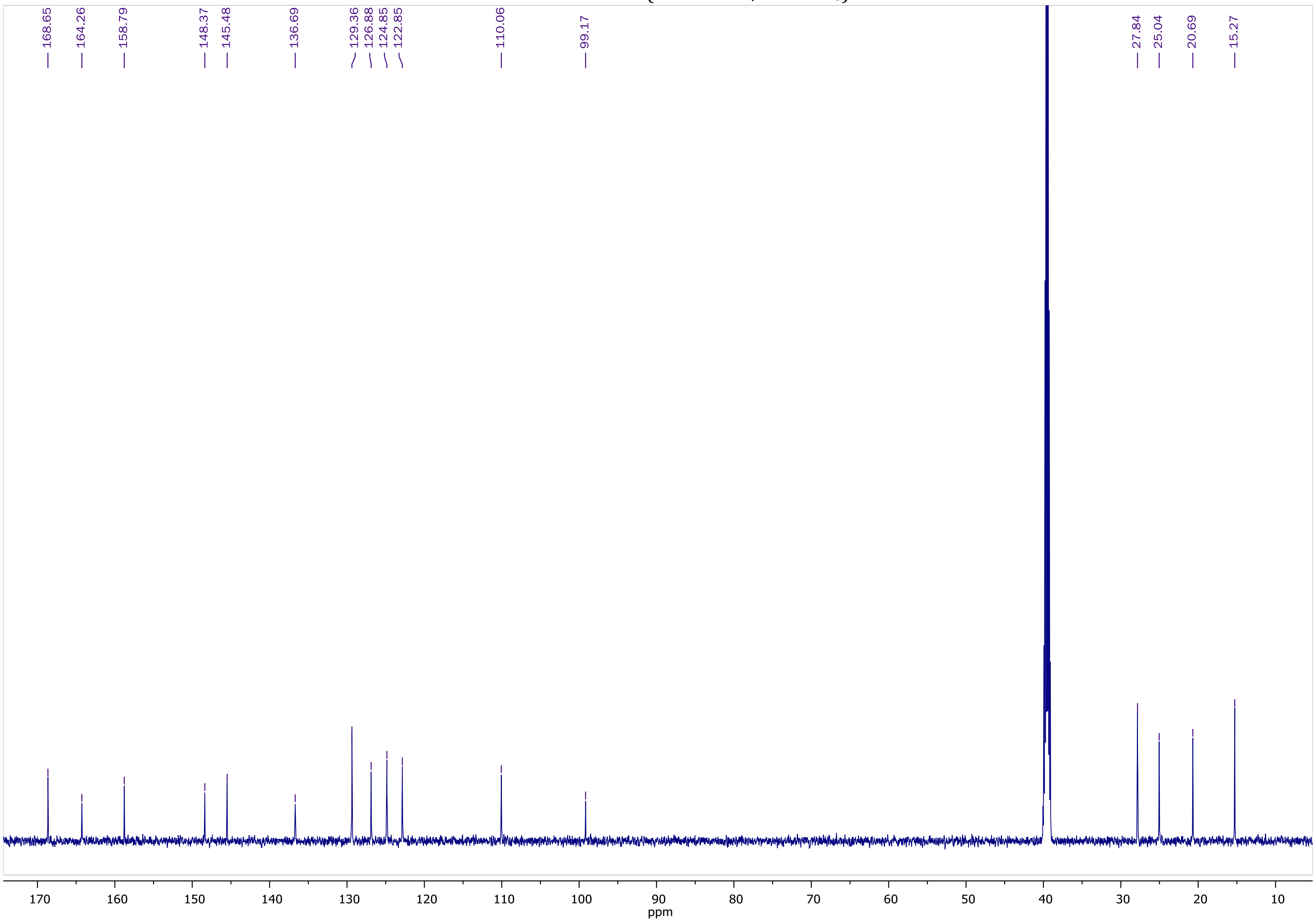

COSY NMR of **ZEV-V3** (600.11 MHz, DMSO-$d_6$)

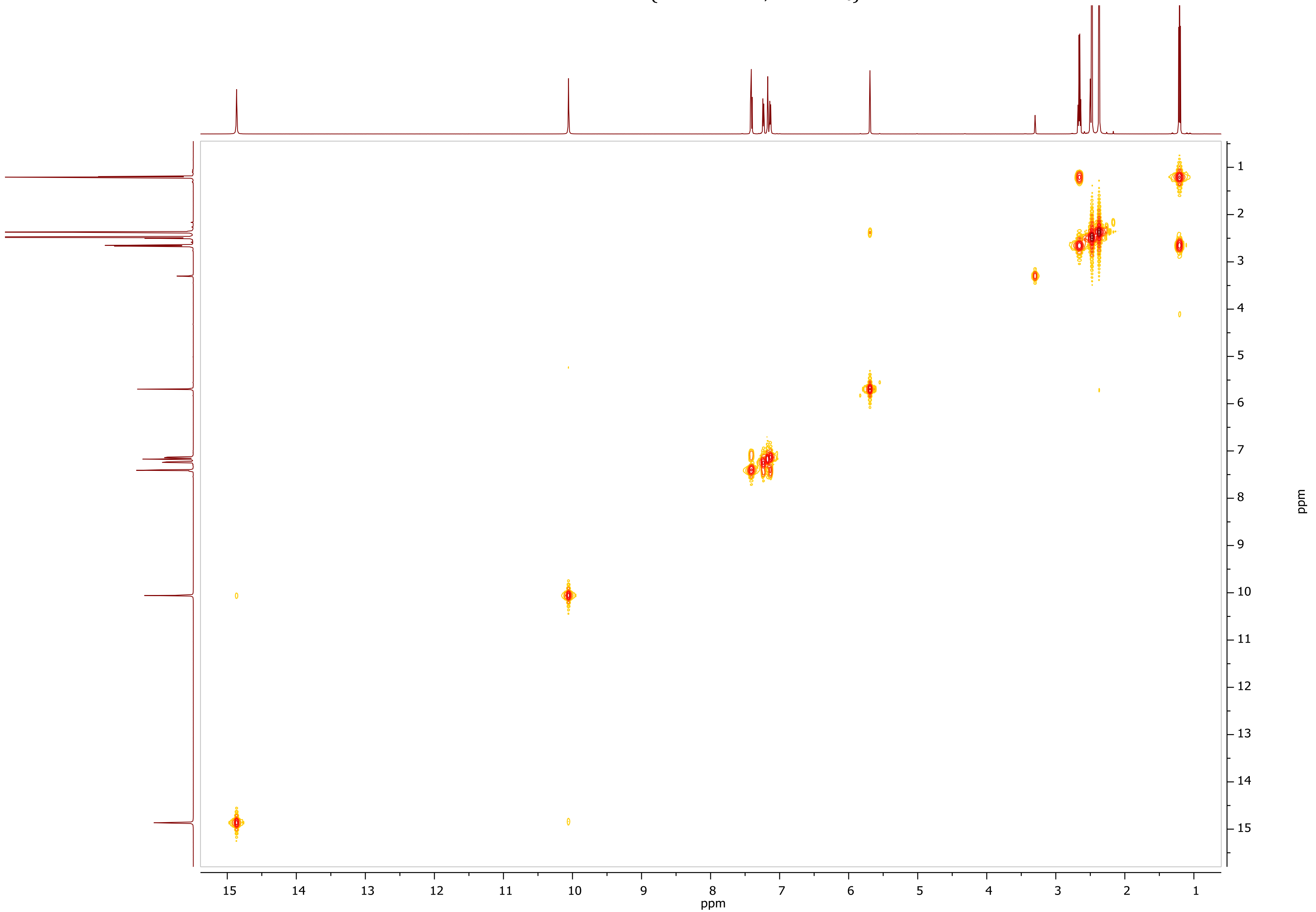

HSQC-DEPT NMR of **ZEV-V3** (600.11 MHz, DMSO-$d_6$)

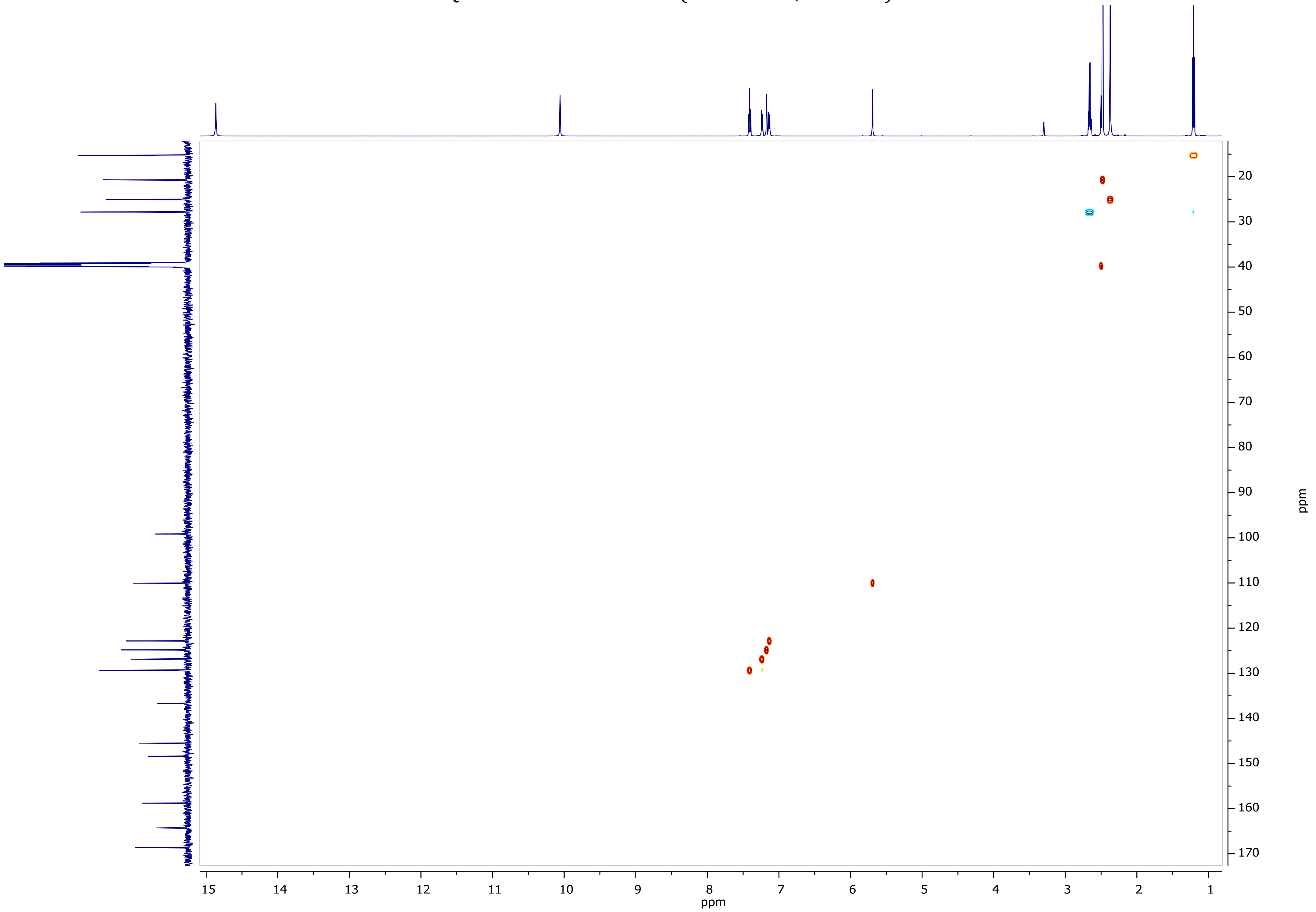

HMBC NMR of **ZEV-V3** (600.11 MHz, DMSO-$d_6$)

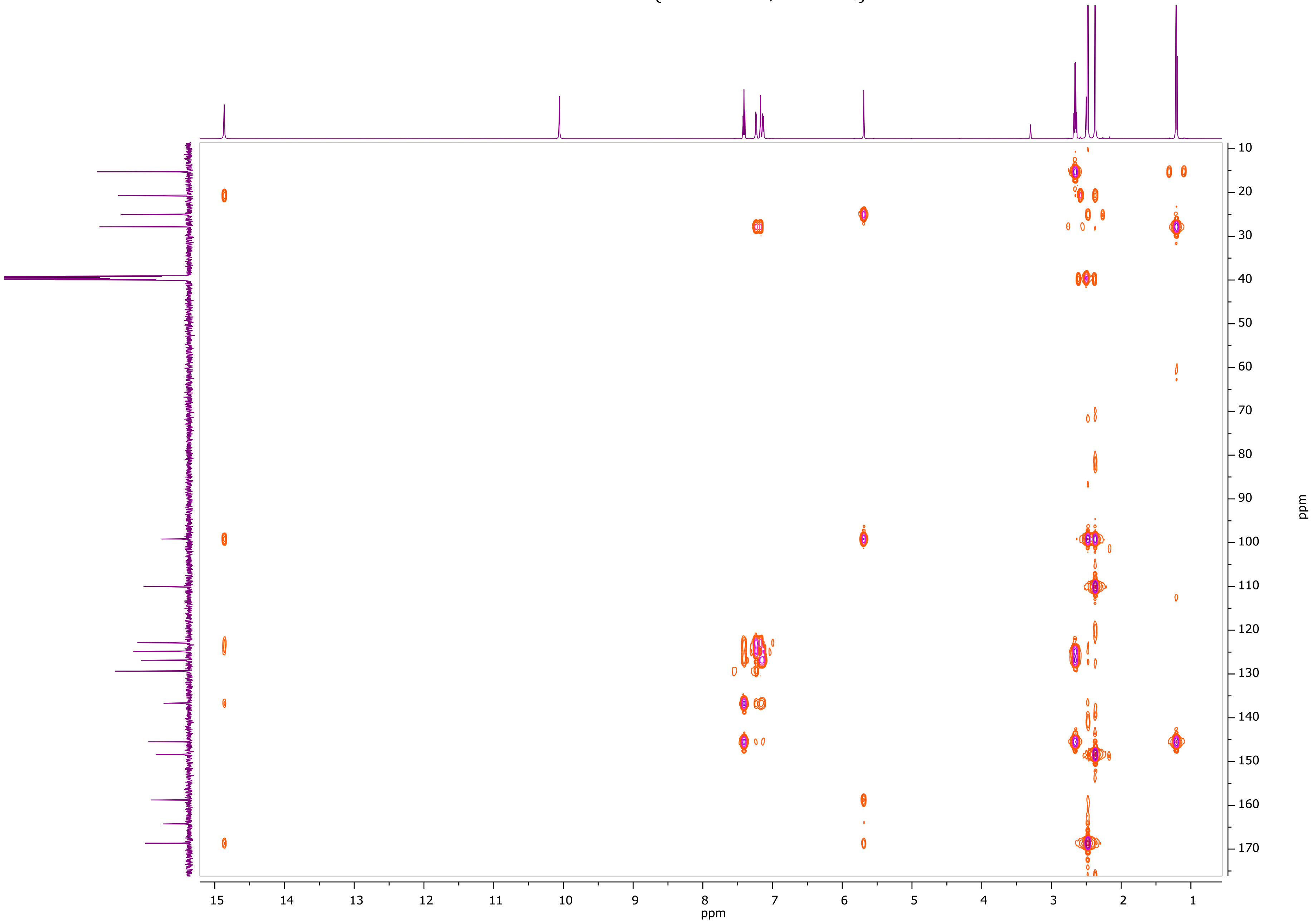

[1]H NMR of **ZEV-V5** (600.11 MHz, DMSO-$d_6$)

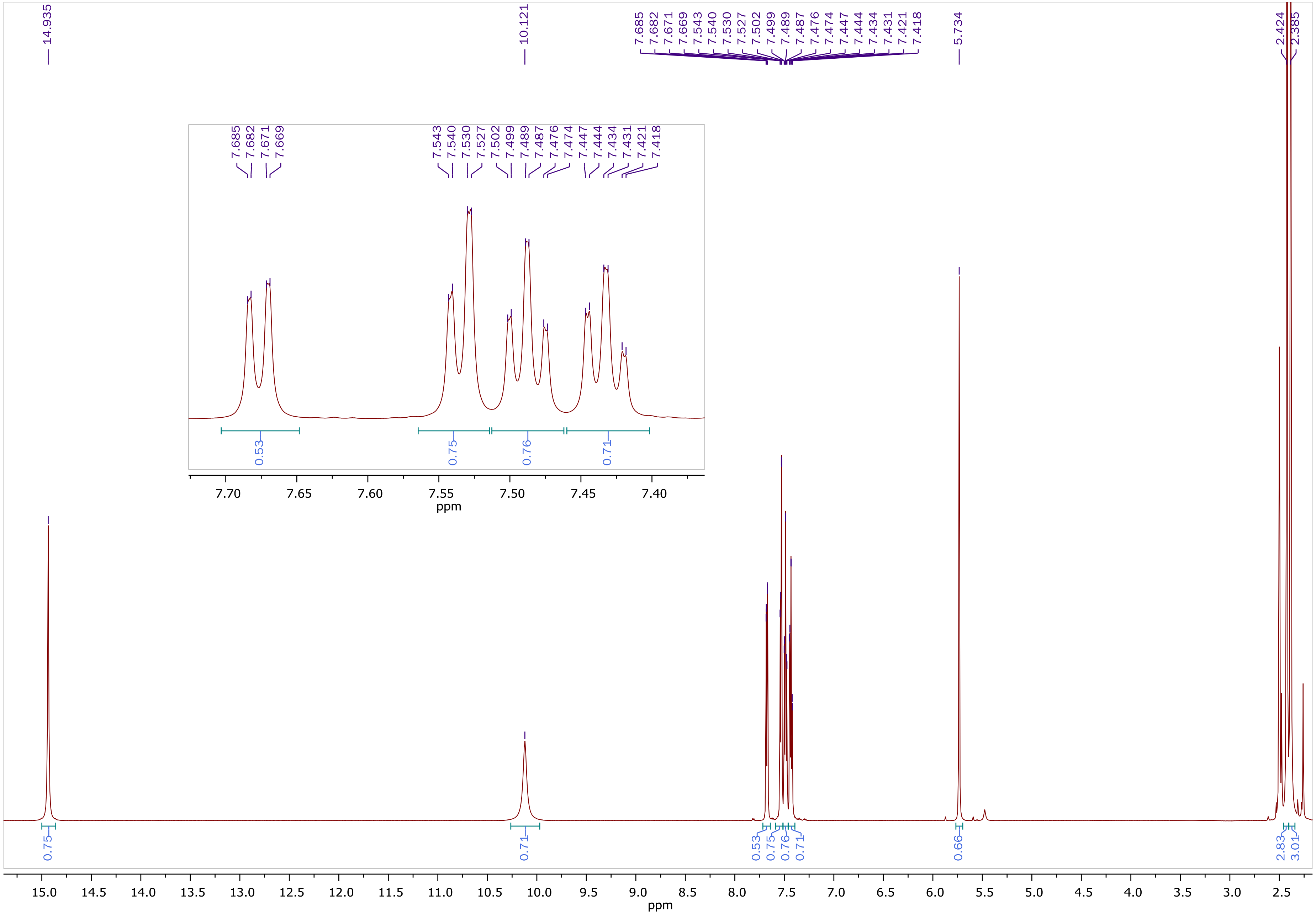

$^{13}C$ NMR of **ZEV-V5** (150.9 MHz, DMSO-$d_6$)

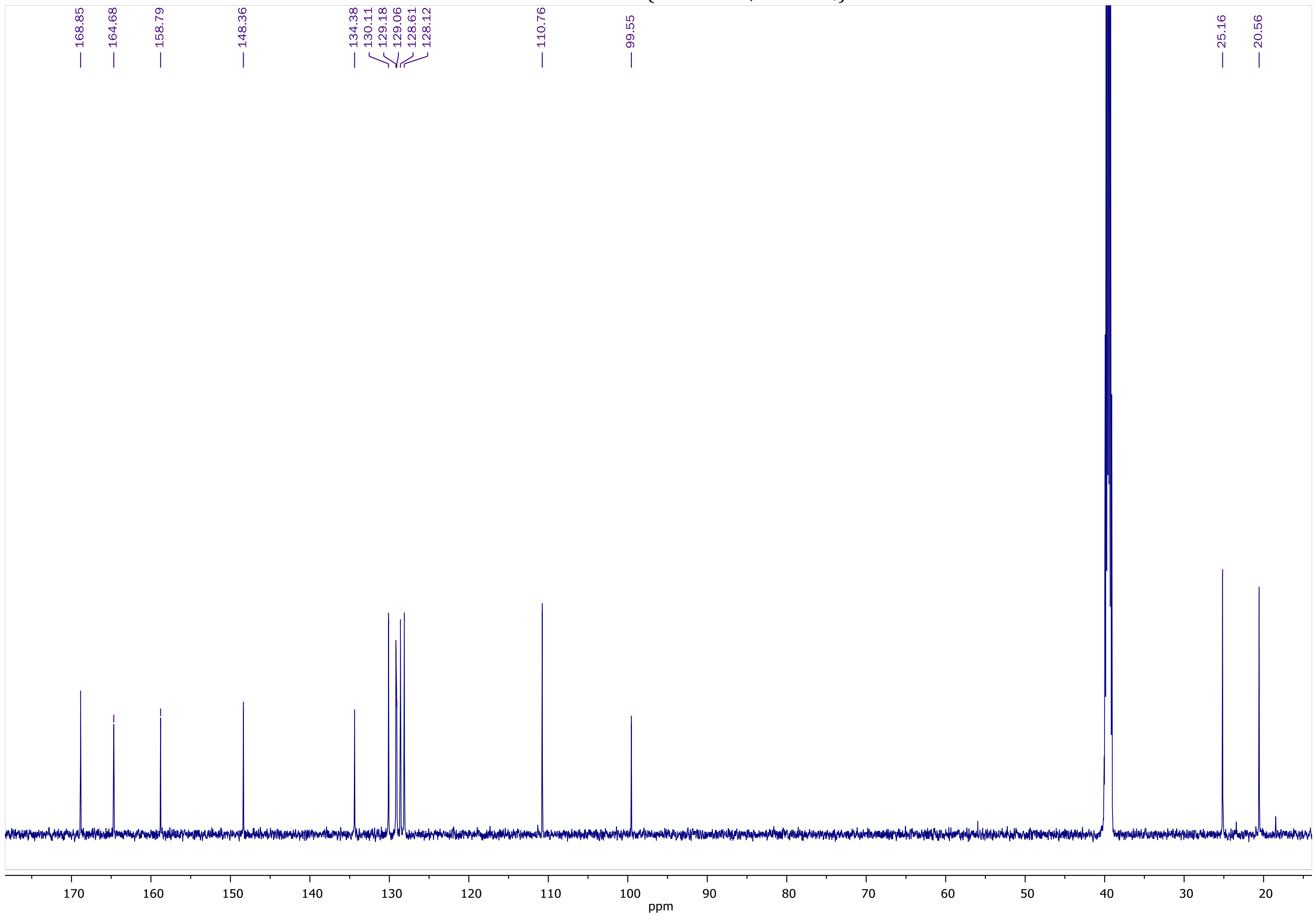

COSY NMR of **ZEV-V5** (600.11 MHz, DMSO-$d_6$)

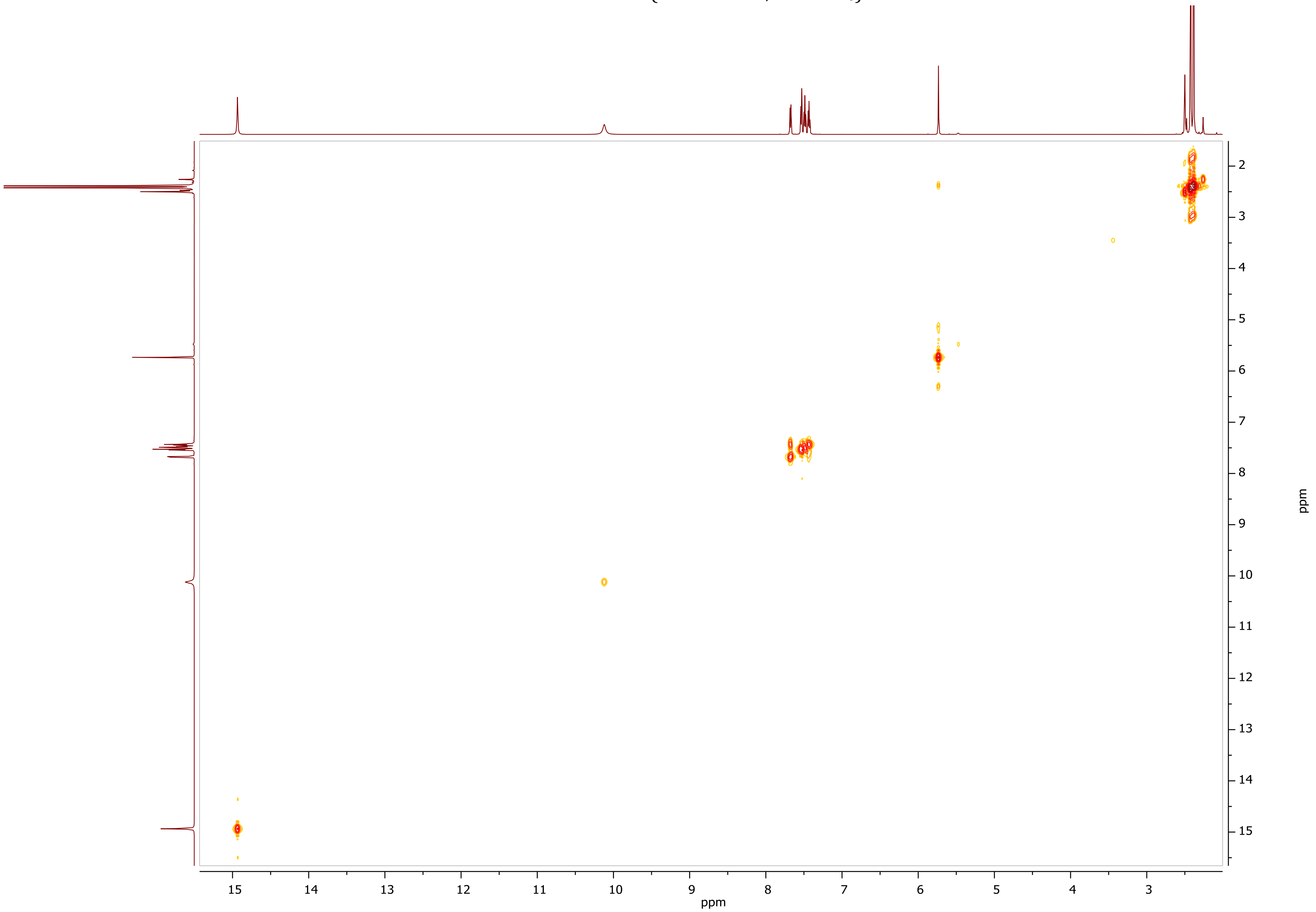

HSQC-DEPT NMR of **ZEV-V5** (600.11 MHz, DMSO-$d_6$)

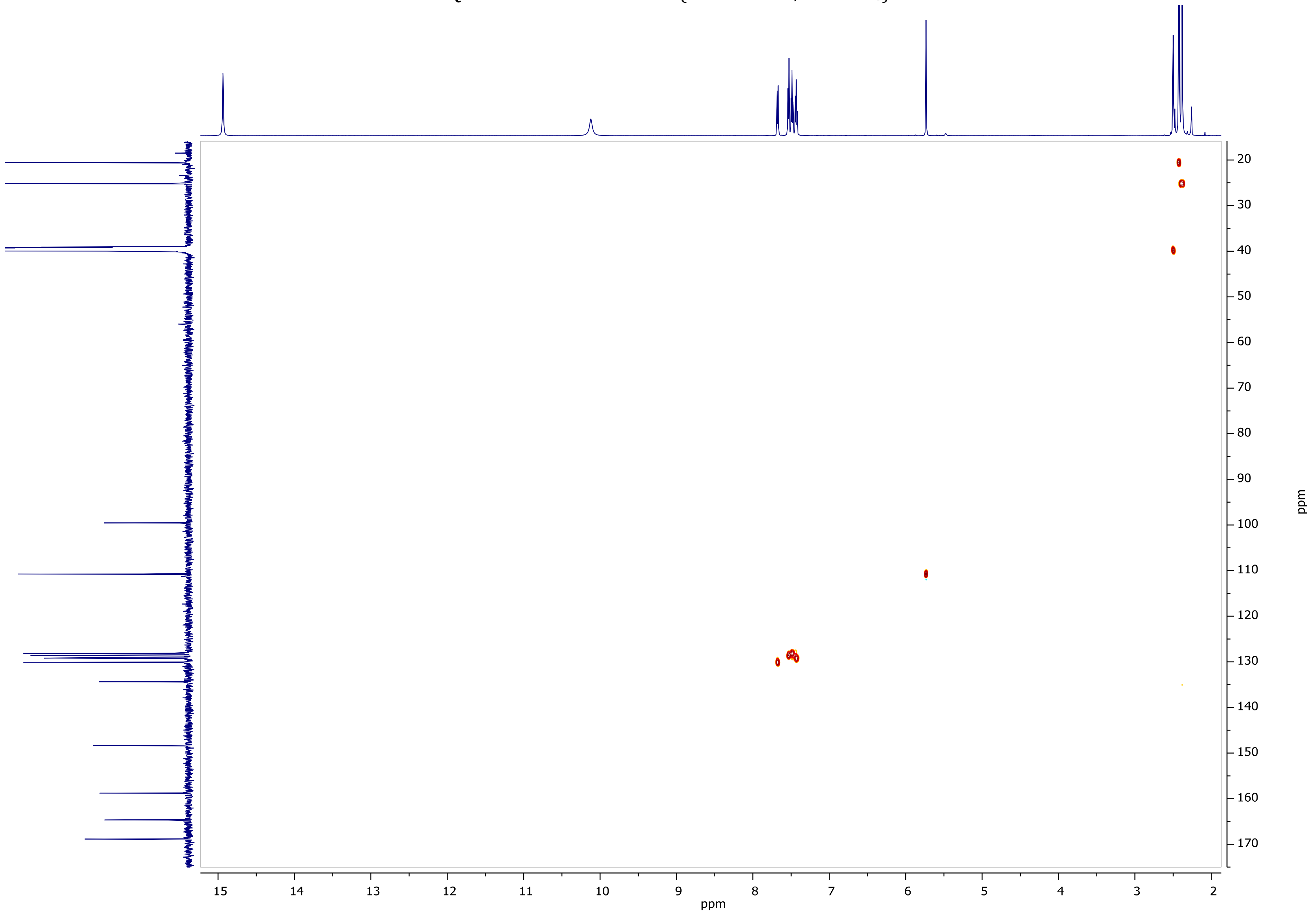

HMBC NMR of **ZEV-V5** (600.11 MHz, DMSO-$d_6$)

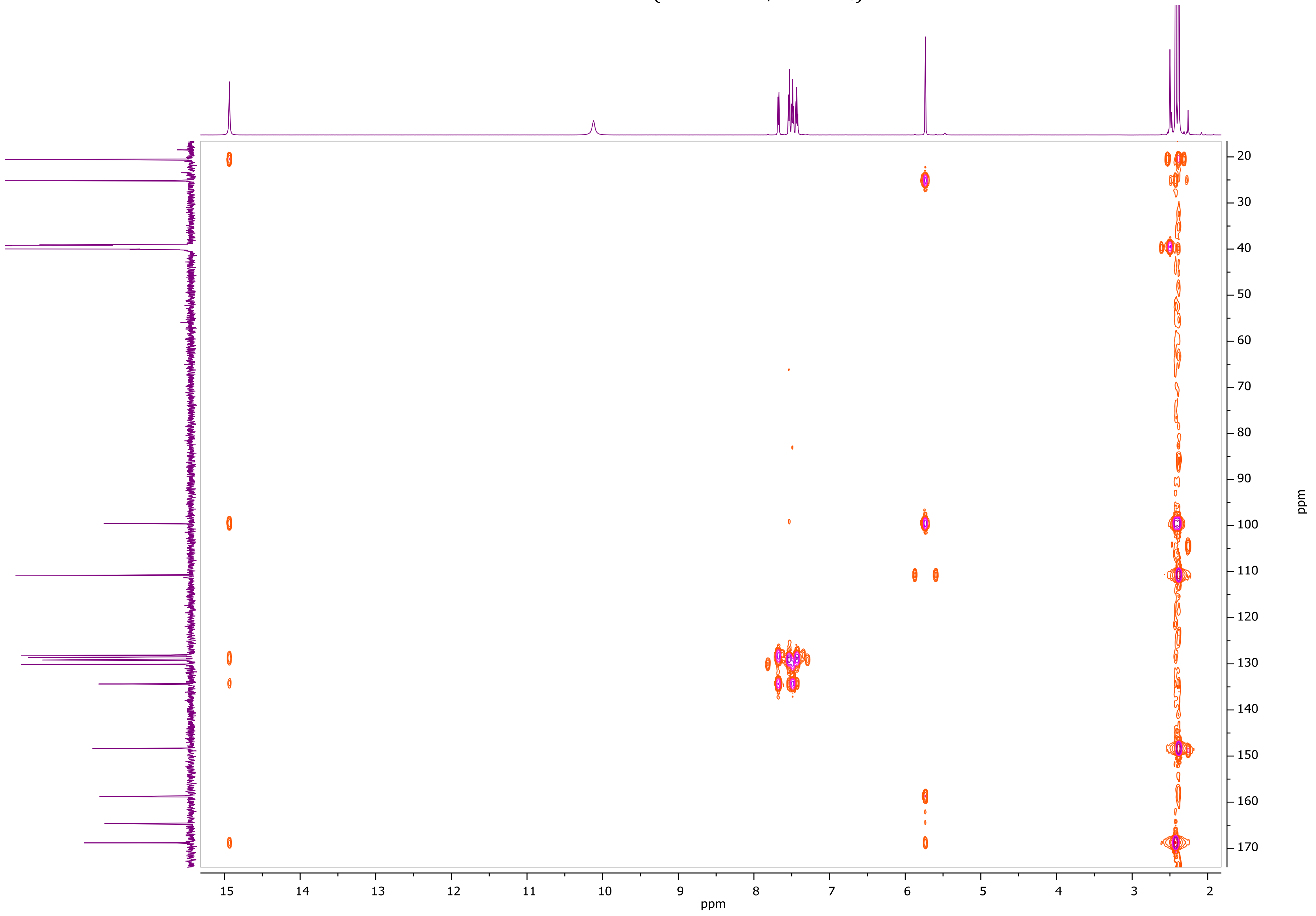

[1]H NMR of **ZEV-V7** (600.11 MHz, DMSO-$d_6$)

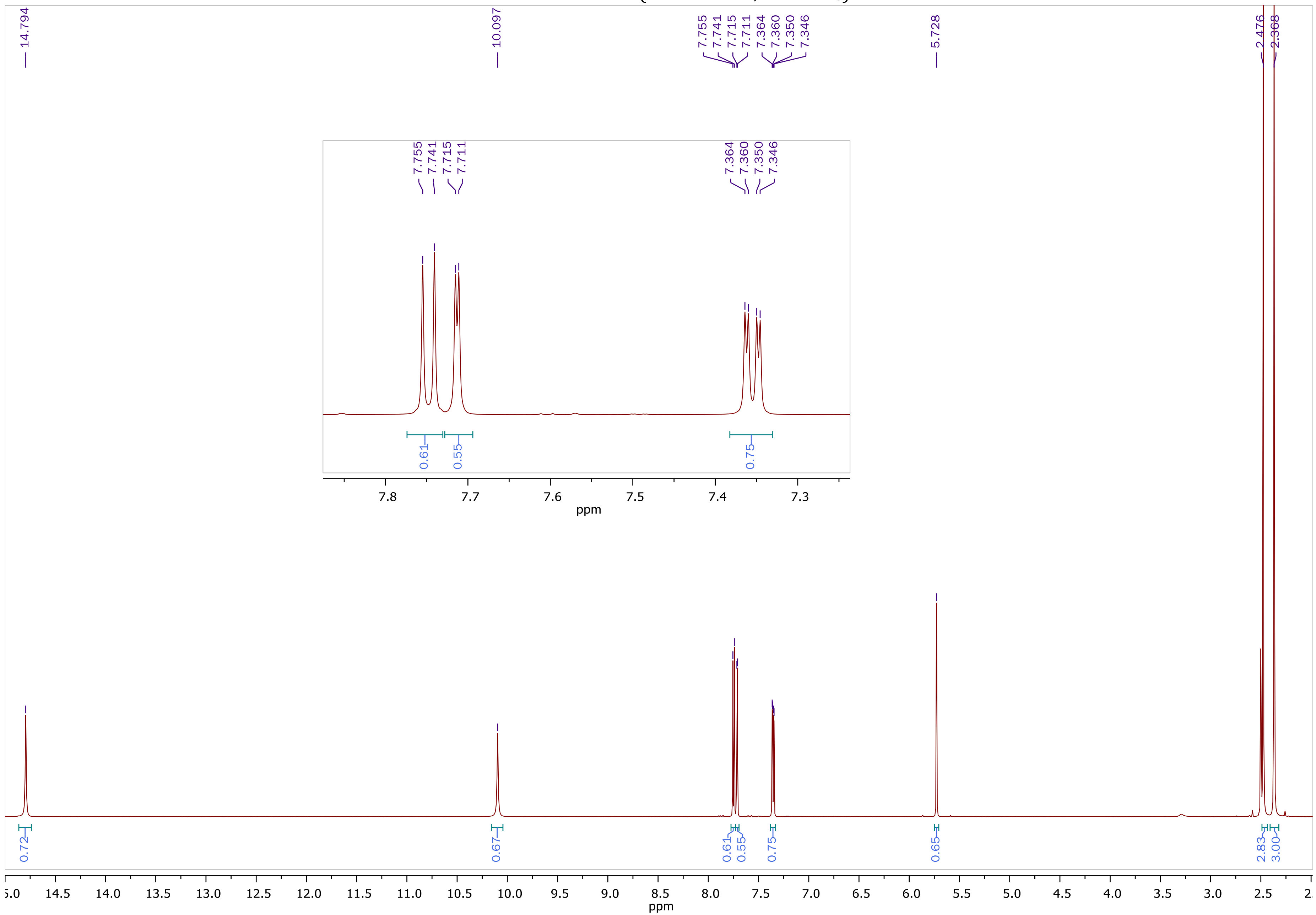

$^{13}C$ NMR of **ZEV-V7** (150.9 MHz, DMSO-$d_6$)

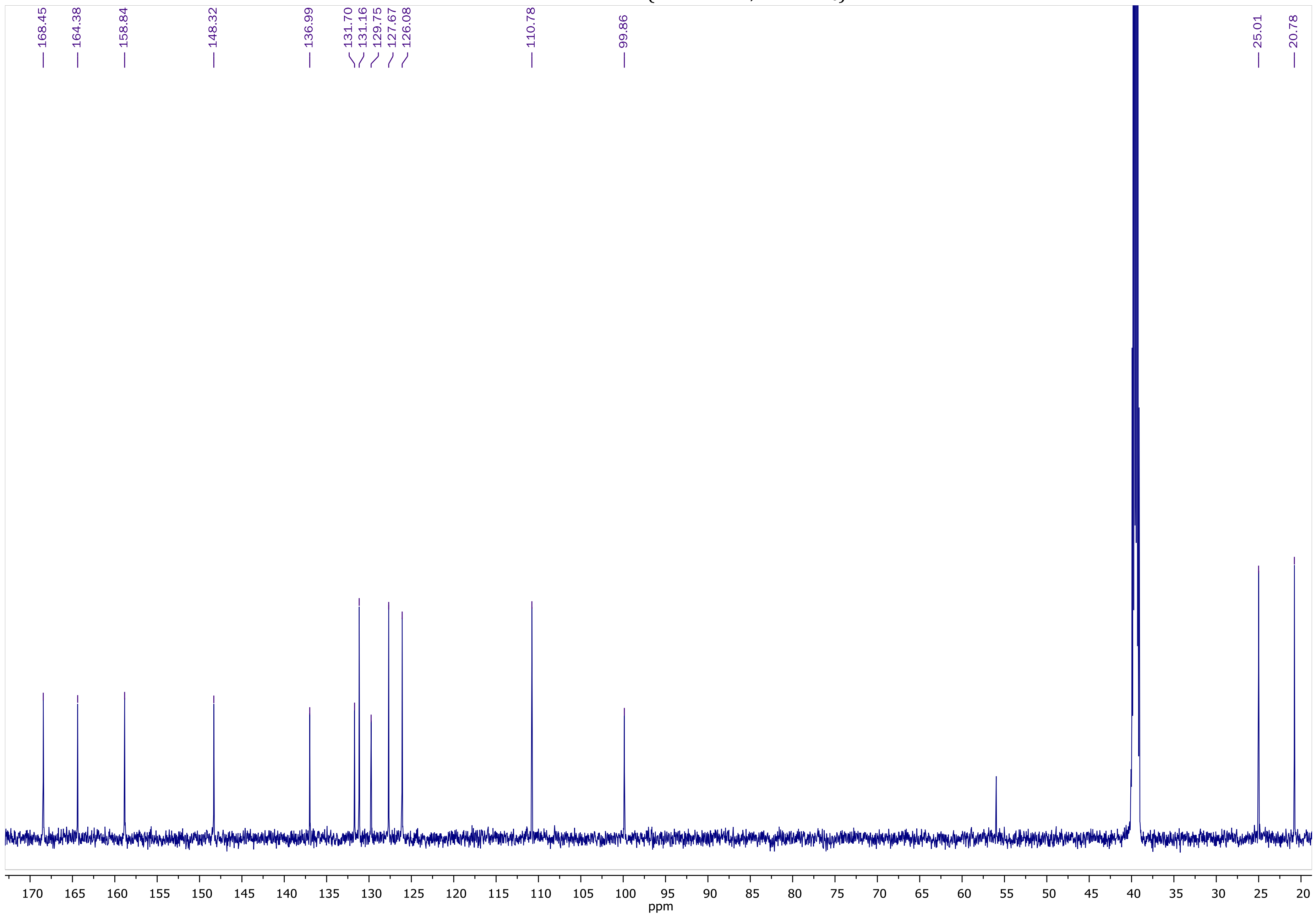

COSY NMR of **ZEV-V7** (600.11 MHz, DMSO-$d_6$)

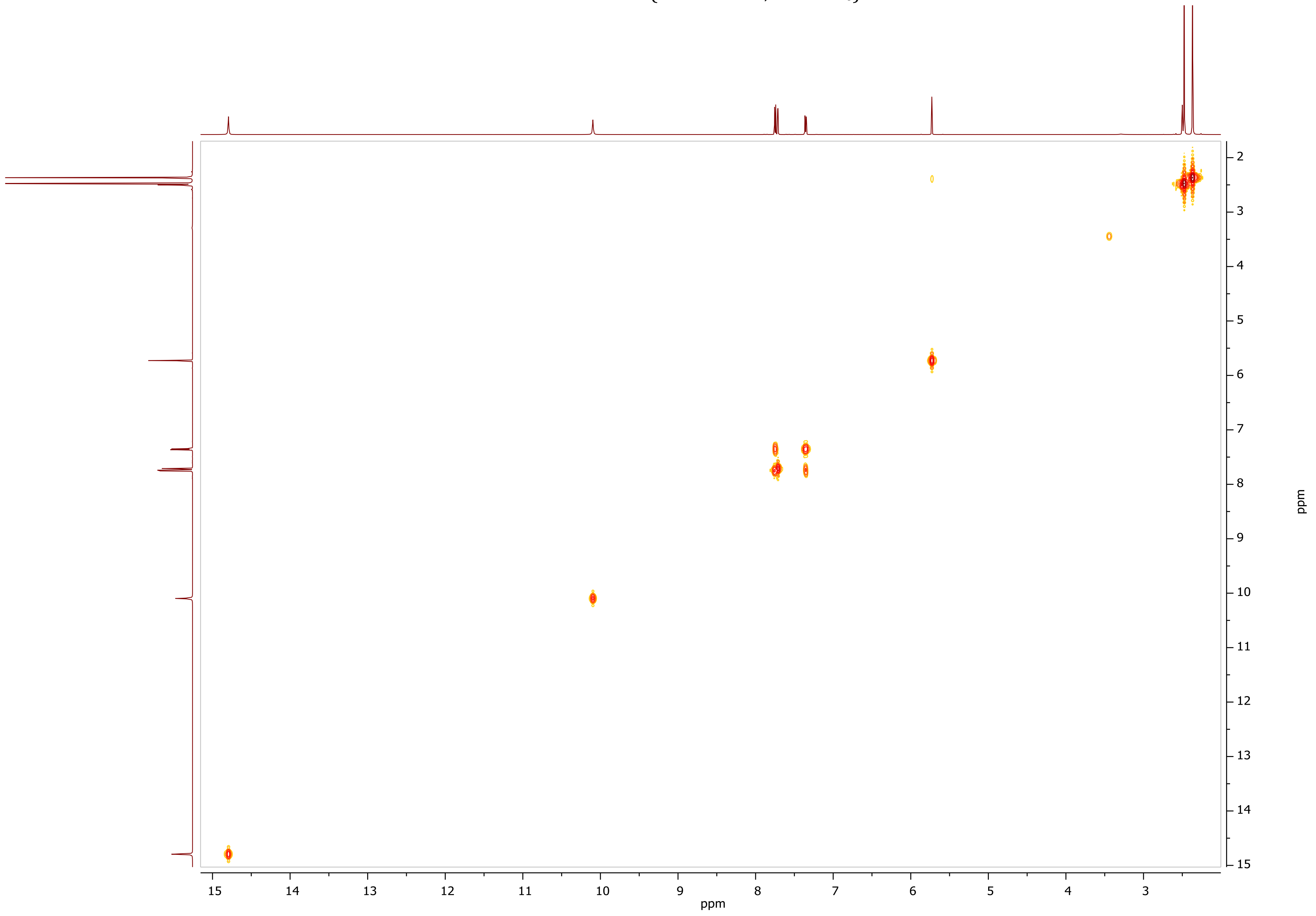

HSQC-DEPT NMR of **ZEV-V7** (600.11 MHz, DMSO-$d_6$)

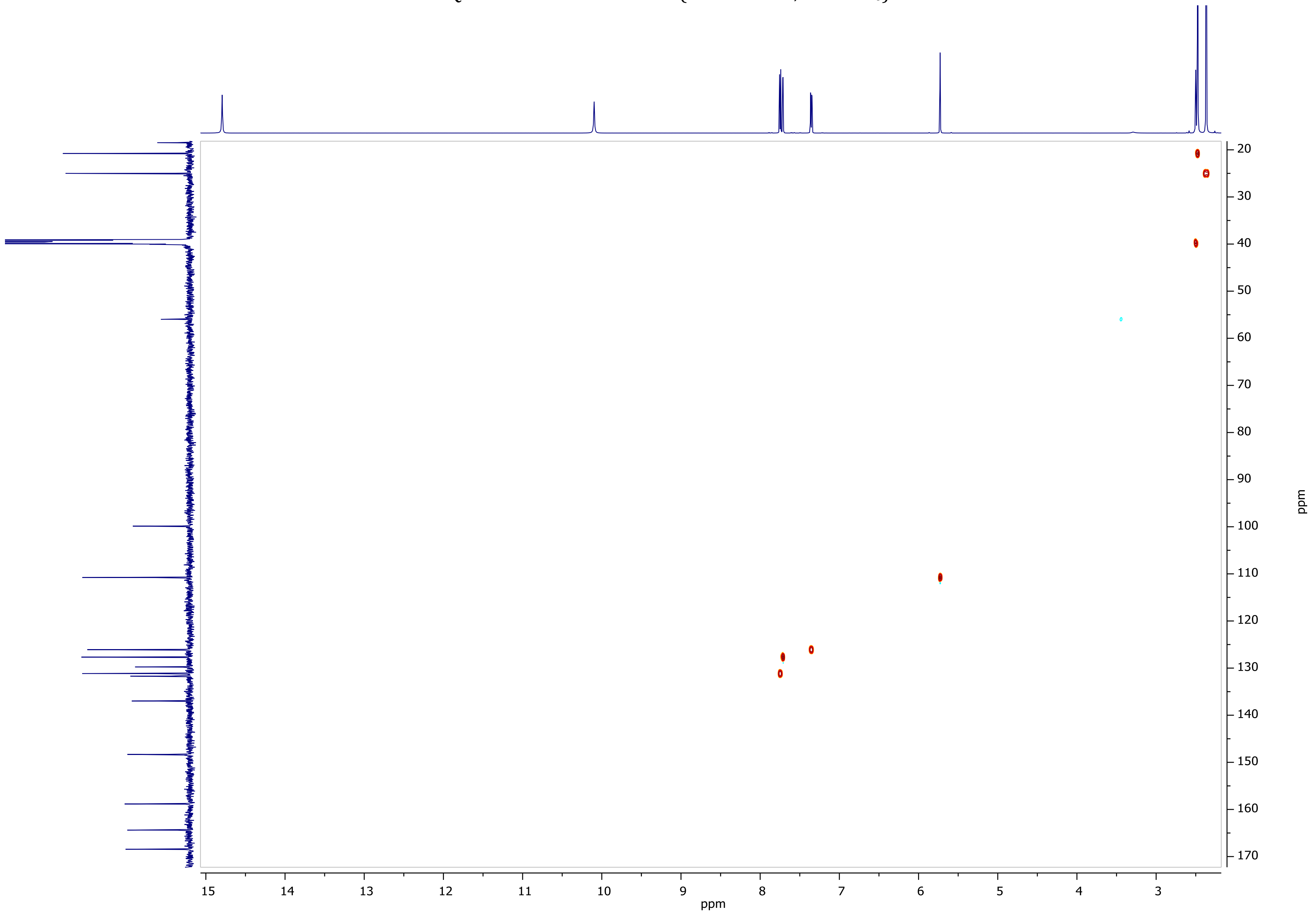

HMBC NMR of **ZEV-V7** (600.11 MHz, DMSO-$d_6$)

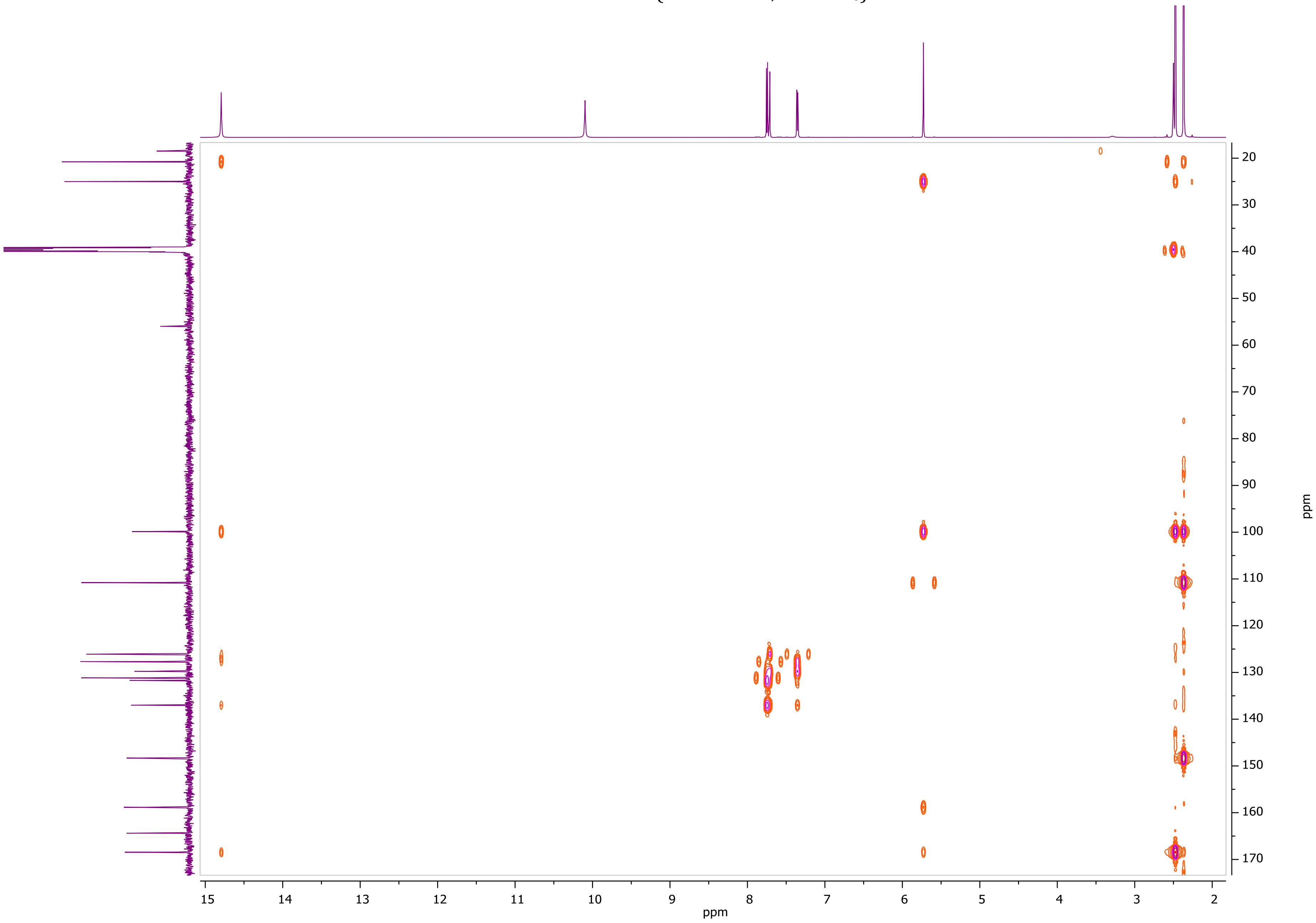

$^1$H NMR of **ZEV-E2** (600.11 MHz, DMSO-$d_6$)

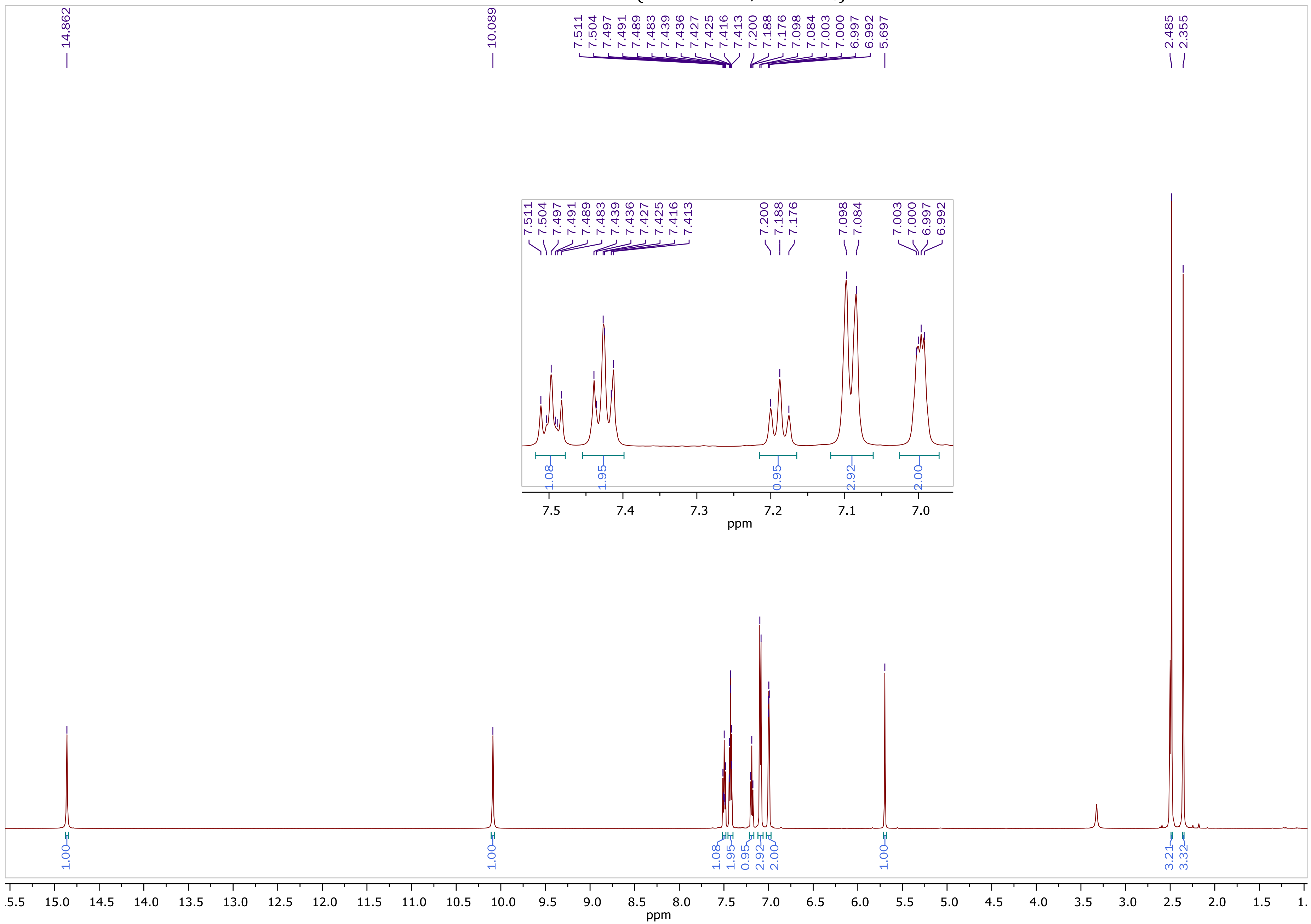

$^{13}$C NMR of **ZEV-E2** (150.9 MHz, DMSO-$d_6$)

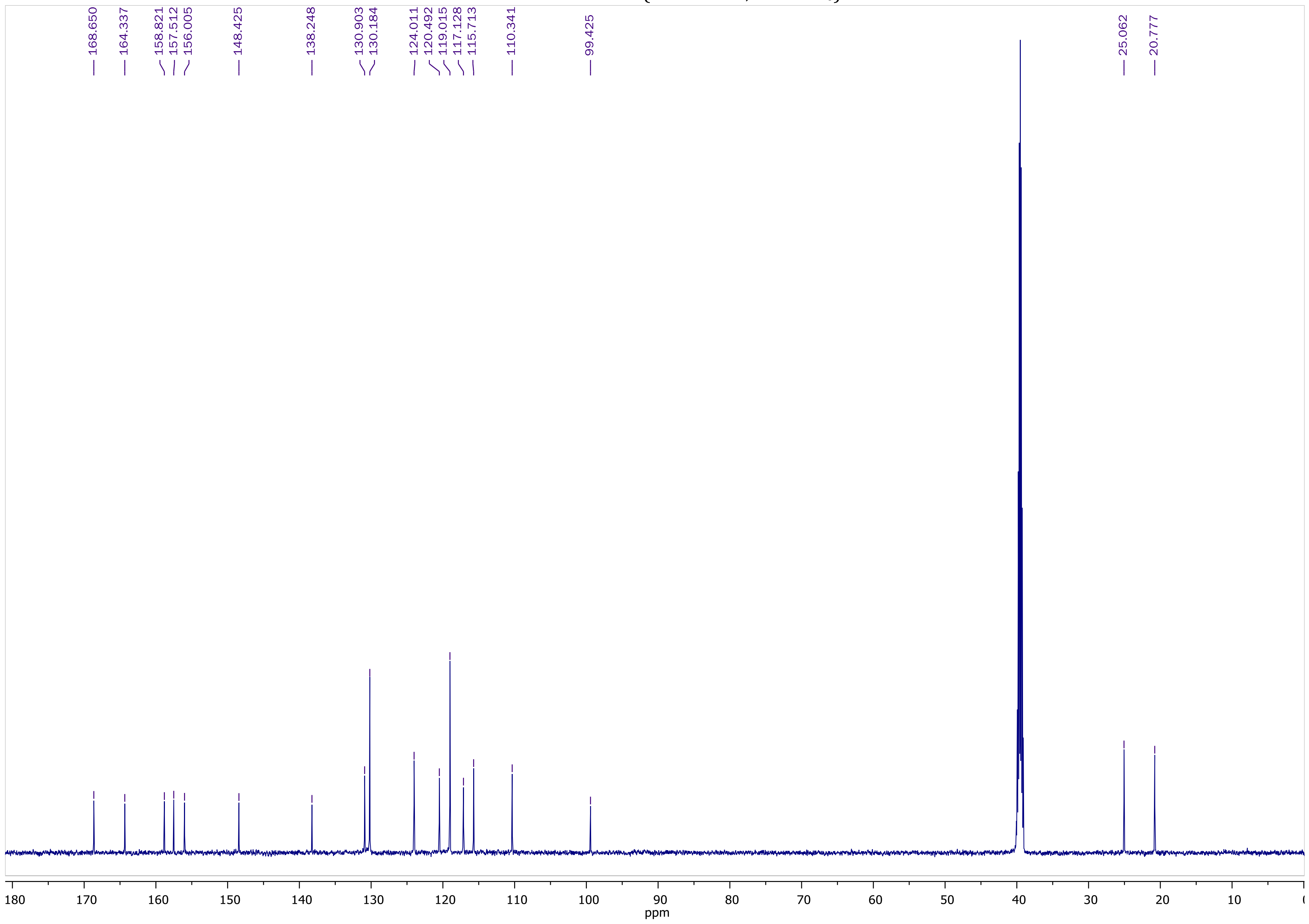

COSY NMR of **ZEV-E2** (600.11 MHz, DMSO-$d_6$)

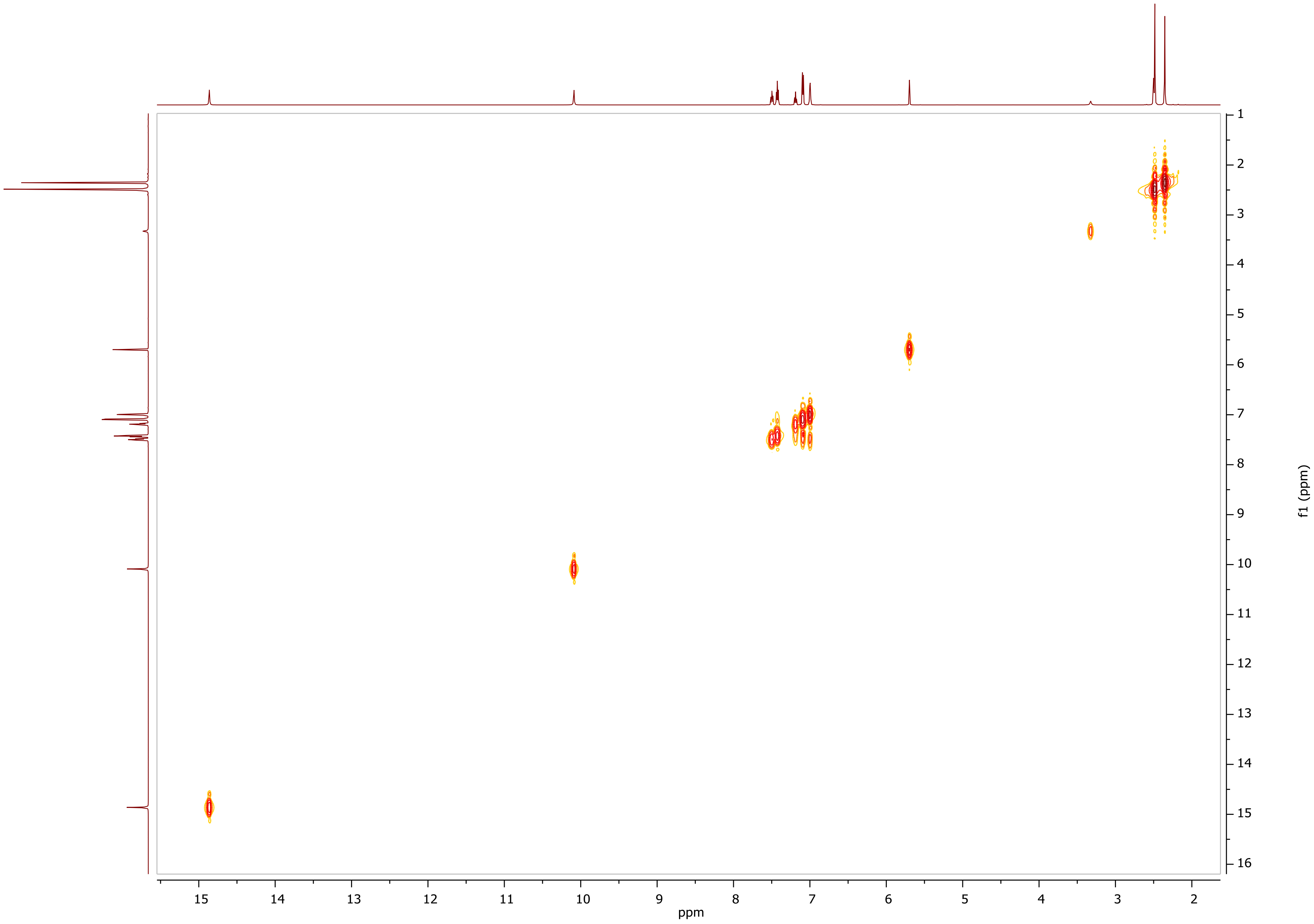

HSQC-DEPT NMR of **ZEV-E2** (600.11 MHz, DMSO-$d_6$)

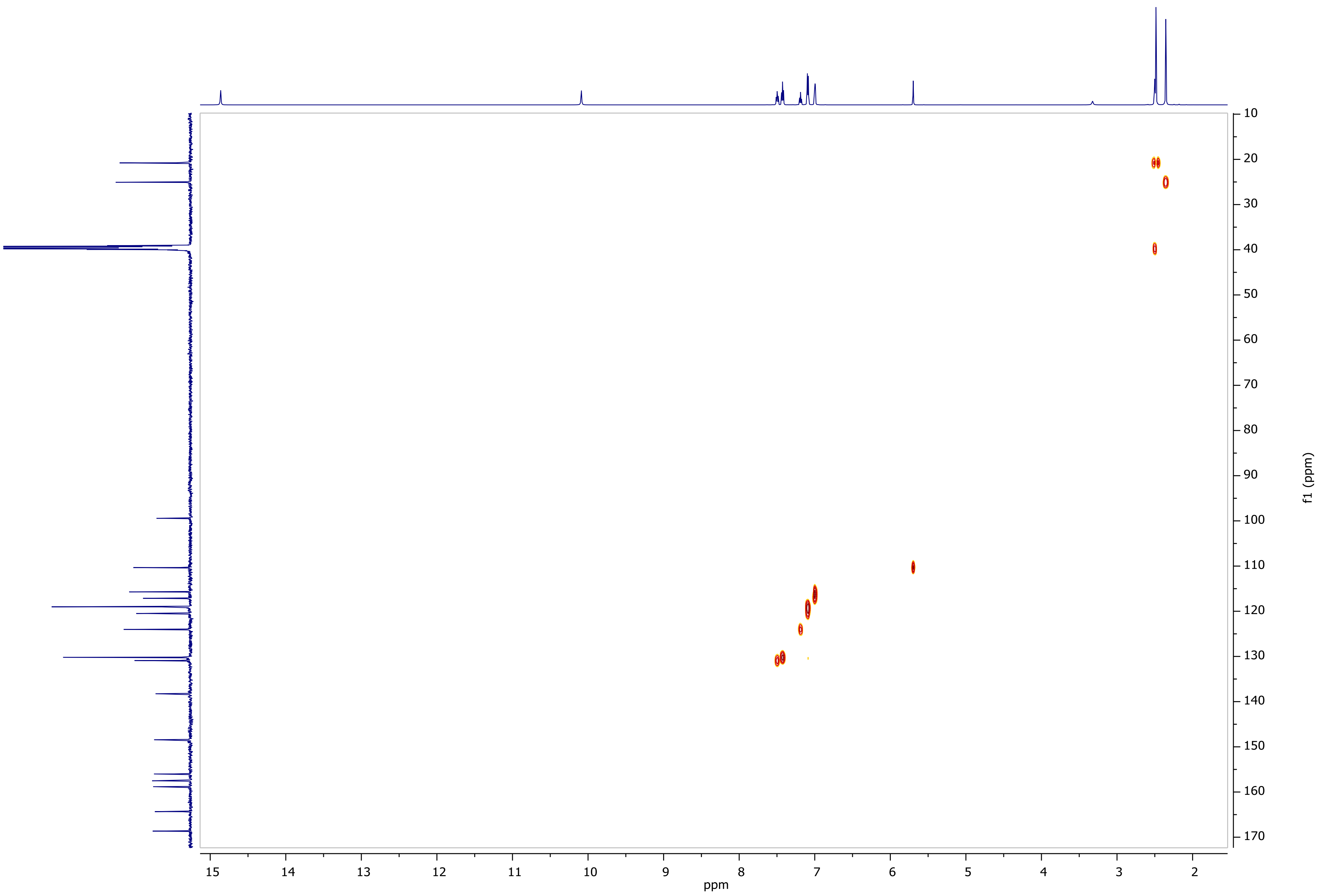

HMBC NMR of **ZEV-E2** (600.11 MHz, DMSO-$d_6$)

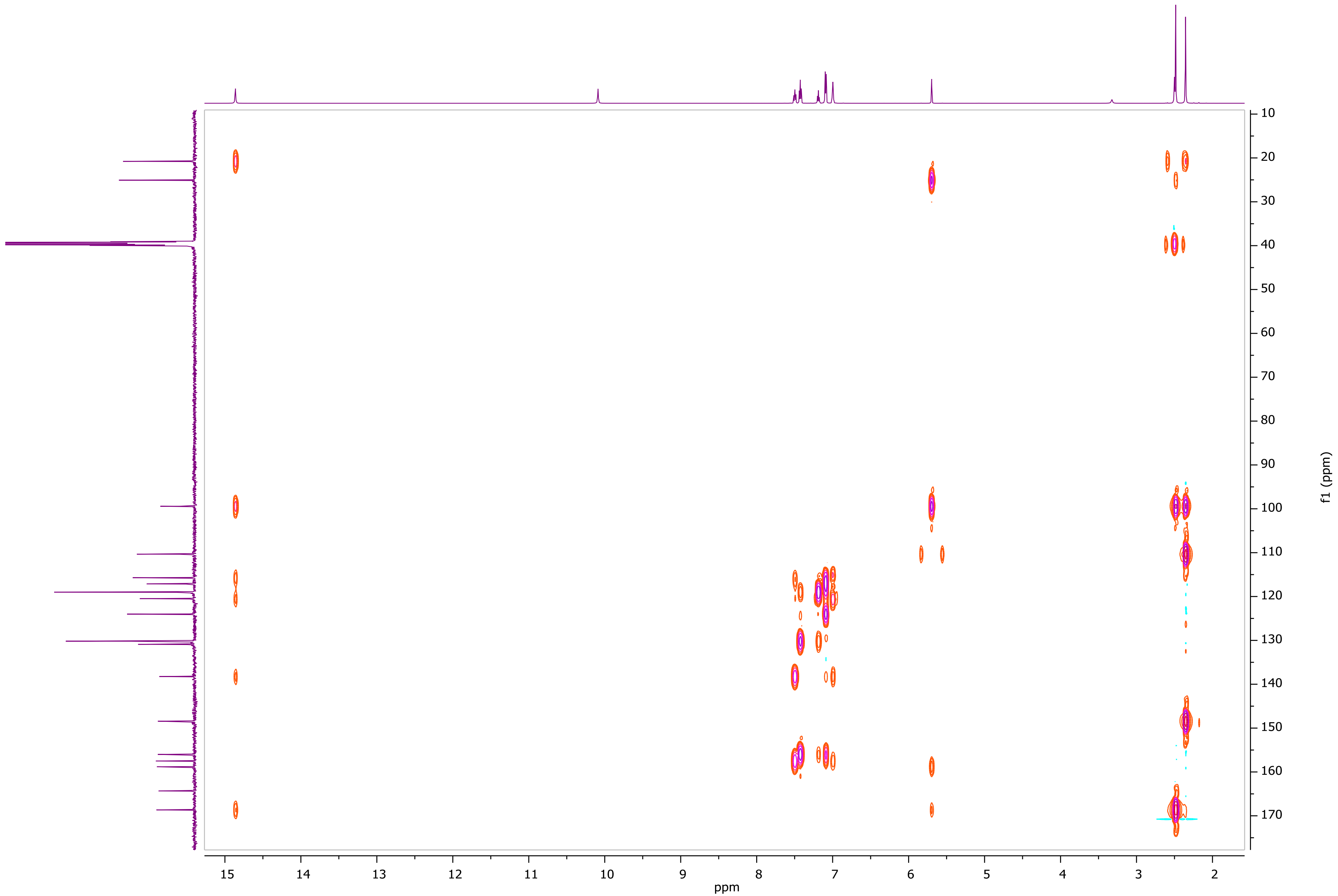

### Supplemental file 2. Structures of compounds in Tables 1 and 2.

#### α-Hydroxytropolones

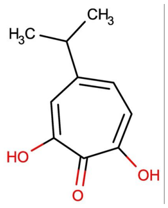

**46**

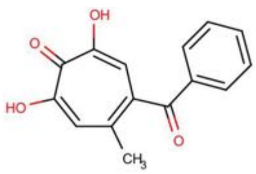

**111**

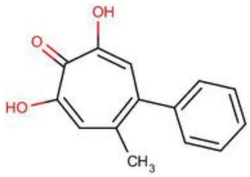

**113**

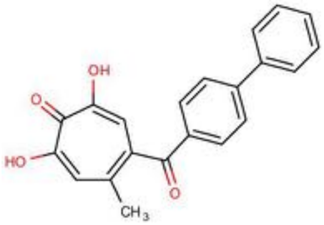

**118**

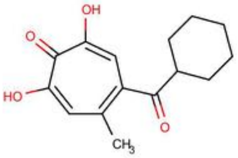

**120**

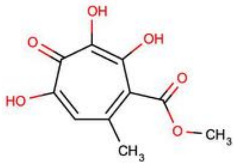

**196**

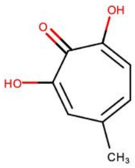

**210**

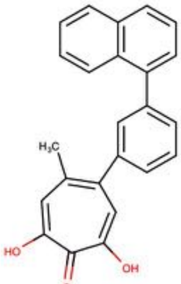

**234**

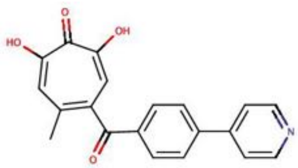

**260**

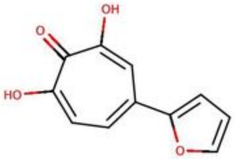

**265**

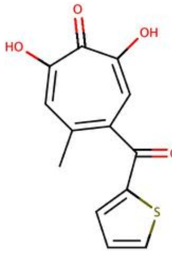

**308**

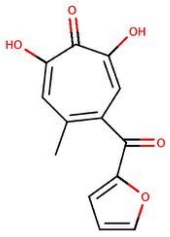

**309**

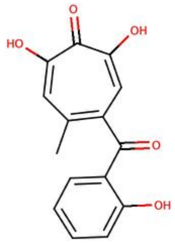

**311**

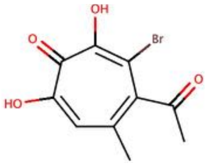

**320**

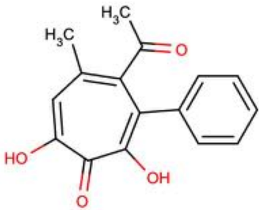

**330**

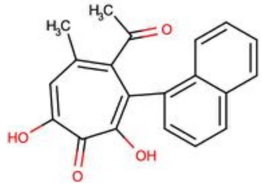

**331**

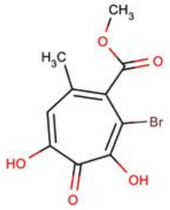

**335**

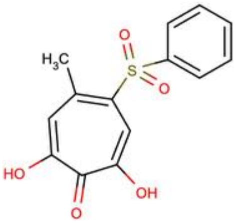

**336**

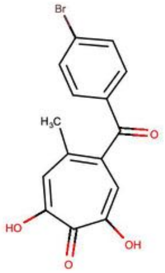

**358**

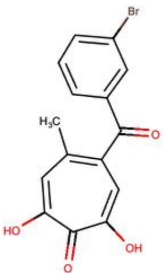

**359**

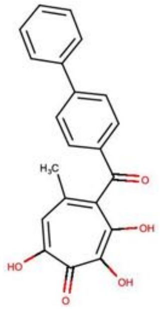

**362**

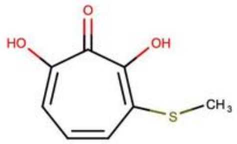

**385**

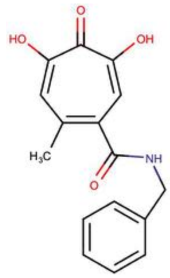

**388**

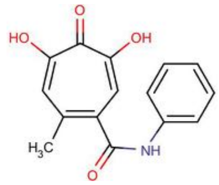

**389**

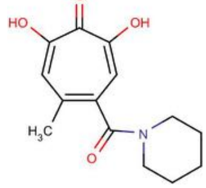

**390**

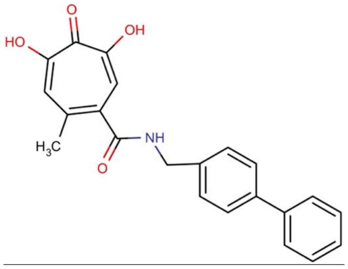

**539**

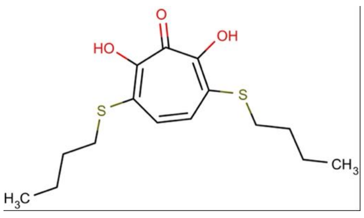

**694**

### α-Hydroxytropolones (Continued)

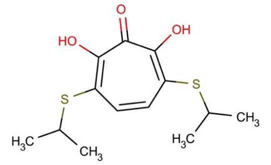

696

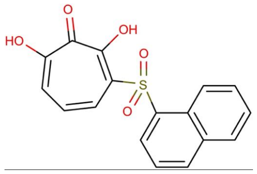

698

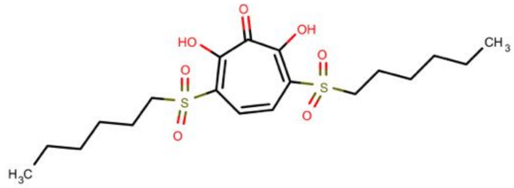

700

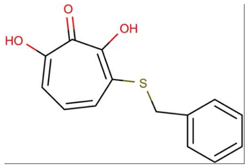

702

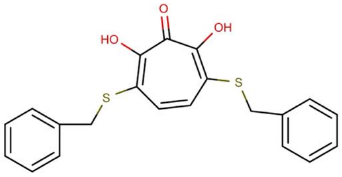

703

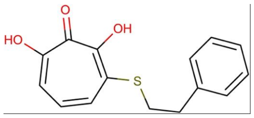

704

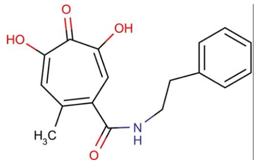

710

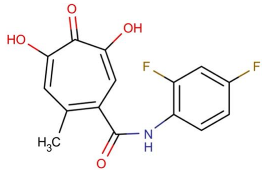

711

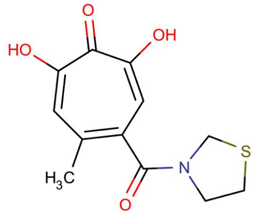

712

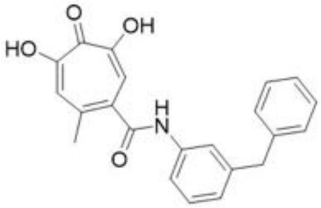

799

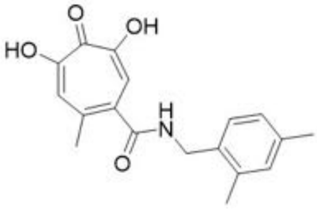

809

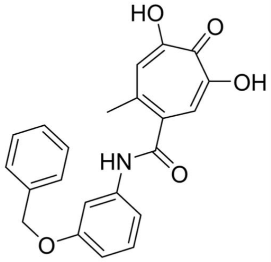

836

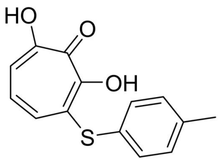

838

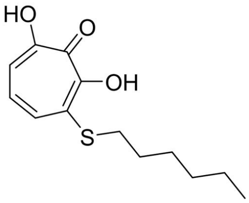

840

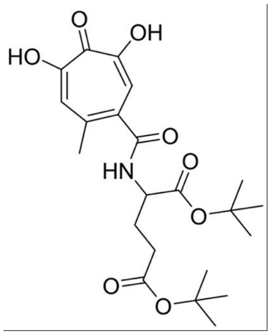

867

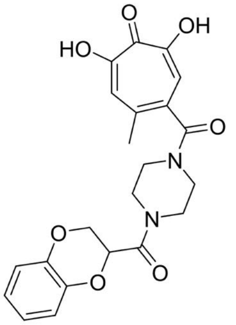

876

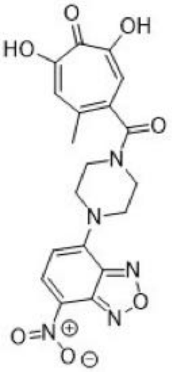

920

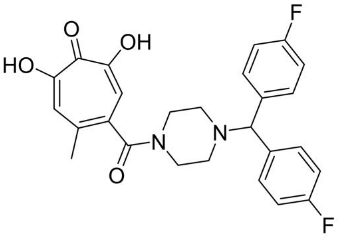

1017

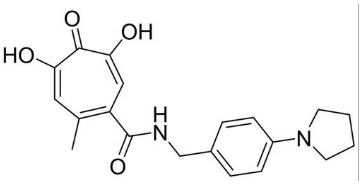

1019

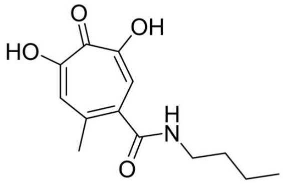

1039

Tropolone and thiotropolones

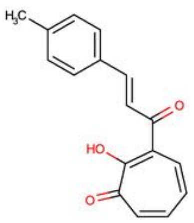

**340**

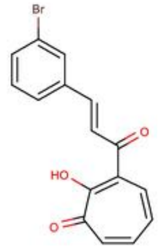

**341**

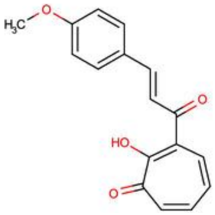

**342**

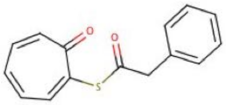

**680**

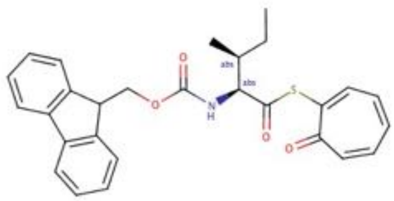

**686**

N-Hydroxypyridinediones

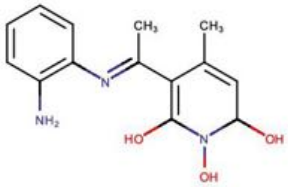

**208**

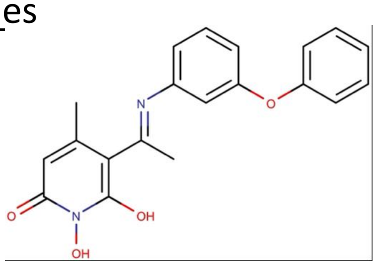

**515**

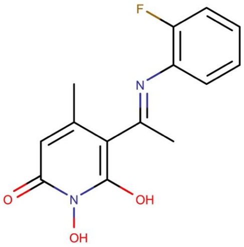

**516**

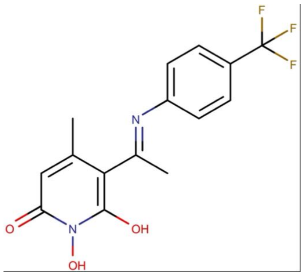

**517**

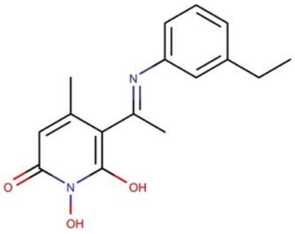

**518**

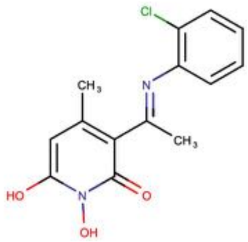

**668**

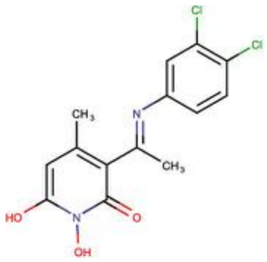

**670**

Dihydronapthalene

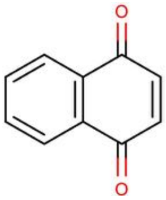

**327**

Example non-hit compounds

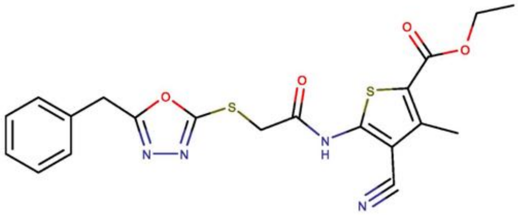

**6**

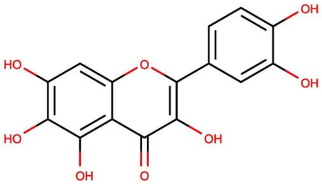

**7**

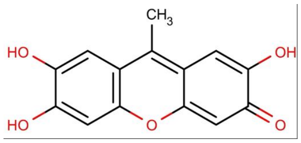

**8**

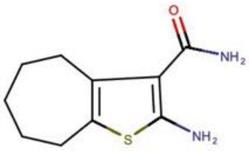

**22**

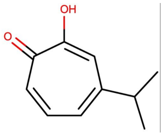

**47**

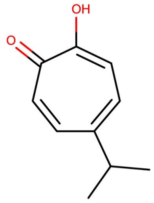

**48**

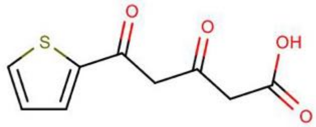

**129**

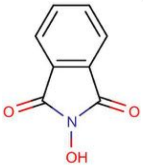

**138**

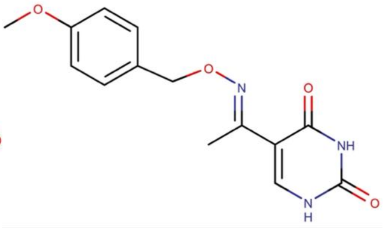

**522**

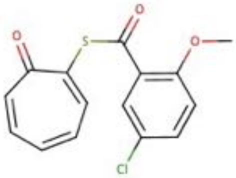

**681**

### Supplemental file 3. Confirmed hit compounds.

α-Hydroxytropolones

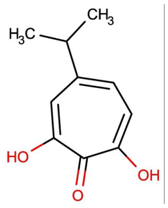

46

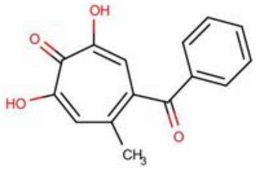

111

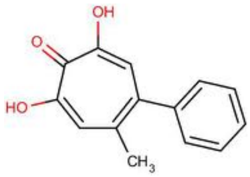

113

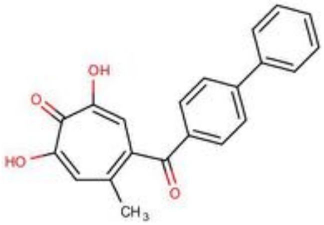

118

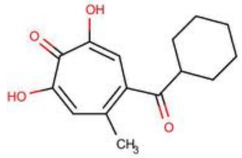

120

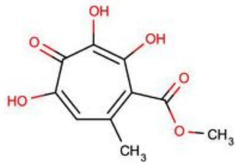

196

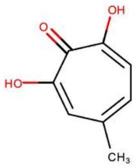

210

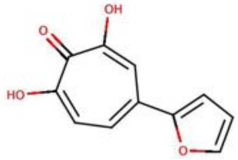

265

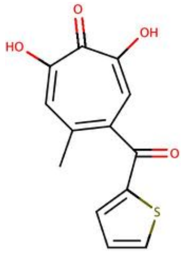

308

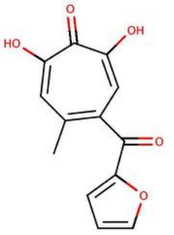

309

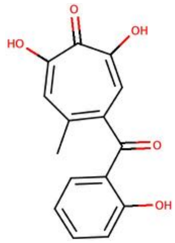

311

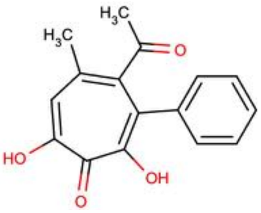

330

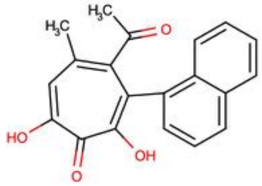

331

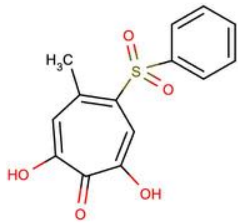

336

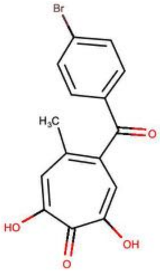

358

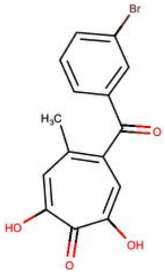

359

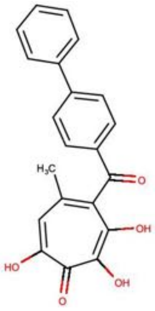

362

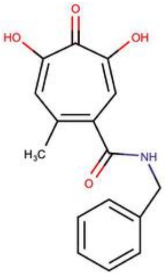

388

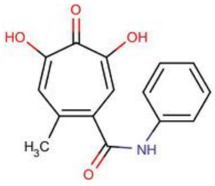

389

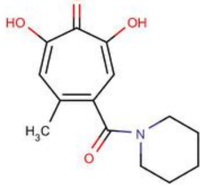

390

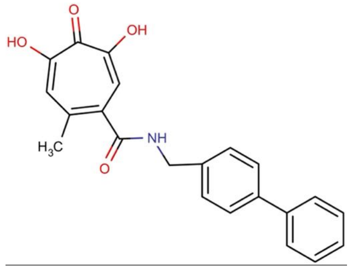

539

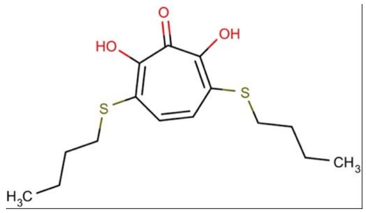

694

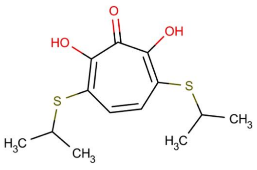

696

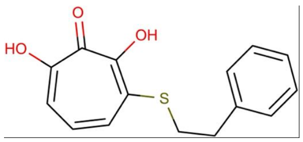

704

#### α-Hydroxytropolones (continued)

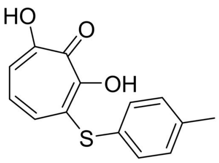


**838**

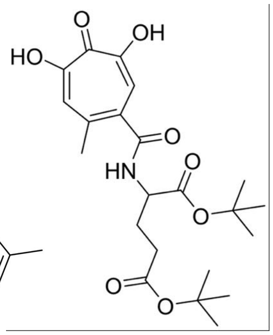


**867**

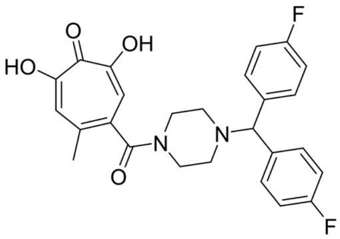


**1017**

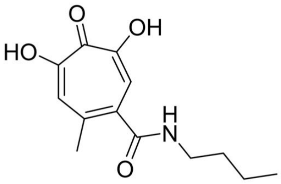


**1039**

#### N-Hydroxypyridinediones

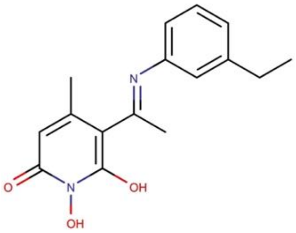


**518**

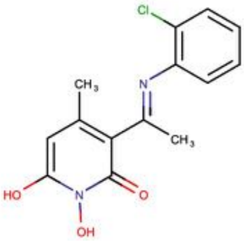


**668**

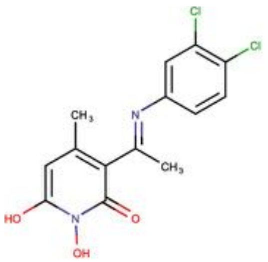


**670**
